## Supplementary material for "Genome mining unveils a class of ribosomal peptides with two amino termini": SI document

[1]Department of Chemical and Biomolecular Engineering, [2]Department of Chemistry, [3]Carl R. Woese Institute for Genomic Biology, [4]Department of Molecular and Cellular Biology, [5]School of Chemical Sciences, NMR Laboratory, [6]Department of Microbiology, [7]Department of Bioengineering, University of Illinois at Urbana-Champaign, Urbana, IL, USA. [8]These authors contributed equally to this work: Hengqian Ren, Shravan R. Dommaraju.

**Supplementary Table 1: Daptide protein co-occurrence table.** RODEO[1] annotation used the Pfam[2], TIGRFAM[3], and custom RiPP recognition element (RRE) profile hidden Markov models (pHMMs)[4]. The top three hits for each annotated CDS were compiled and counted for frequency. The top 32 highest occurring pHMM hits in daptide BGCs are listed with custom pHMM models for daptide detection shown in red.

| Count | % Co-occurrence | pHMM ID | pHMM description |
|---|---|---|---|
| **486** | 101% | PF00202 | Aminotransferase class-III |
| **486** | 101% | TIGR00707 | argD: transaminase, acetylornithine/succinylornithine family |
| **484** | 100% | TIGR00508 | bioA: adenosylmethionine-8-amino-7-oxononanoate transaminase |
| **456** | 94% | PF13649 | Methyltransferase domain |
| **452** | 94% | TIGR01188 | drrA: daunorubicin resistance ABC transporter, ATP-binding protein |
| **381** | 79% | PF08241 | Methyltransferase domain |
| **377** | 78% | TIGR03740 | galliderm_ABC: lantibiotic protection ABC transporter, ATP-binding subunit |
| **375** | 78% | TIGR03522 | GldA_ABC_ATP: gliding motility-associated ABC transporter ATP-binding subunit GldA |
| **319** | 66% | **Actino_DapB_RRE** | C-terminal RRE domain from Actinomycetota DapB proteins |
| **307** | 64% | PF00465 | Iron-containing alcohol dehydrogenase |
| **302** | 63% | TIGR03405 | Phn_Fe-ADH: phosphonate metabolism-associated iron-containing alcohol dehydrogenase |
| **289** | 60% | **Actino_DapP_RRE** | N-terminal RRE domain from Actinomycetota DapP proteins |
| **192** | 40% | PF13561 | Enoyl-(Acyl carrier protein) reductase |
| **185** | 38% | TIGR02638 | lactal_redase: lactaldehyde reductase |
| **183** | 38% | PF12698 | ABC-2 family transporter protein |
| **178** | 37% | PF13489 | Methyltransferase domain |
| **176** | 36% | **Bacill_DapB_RRE** | C-terminal RRE domain from Bacillota DapB proteins |
| **163** | 34% | TIGR01830 | 3oxo_ACP_reduc: 3-oxoacyl-[acyl-carrier-protein] reductase |
| **154** | 32% | PF00528 | Binding-protein-dependent transport system inner membrane component |
| **145** | 30% | PF02163 | Peptidase family M50 |
| **142** | 29% | PF00583 | Acetyltransferase (GNAT) family |
| **142** | 29% | **Bacill_DapP_RRE** | N-terminal RRE domain from Bacillota DapP proteins |
| **139** | 29% | PF02624 | YcaO cyclodehydratase, ATP-ad Mg2+-binding |
| **134** | 28% | TIGR03604 | TOMM_cyclo_SagD: thiazole/oxazole-forming peptide maturase, SagD family component |
| **124** | 26% | PF00069 | Protein kinase domain |
| **123** | 26% | PF00296 | Luciferase-like monooxygenase |
| **120** | 25% | TIGR03564 | F420_MSMEG_4879: F420-dependent oxidoreductase, MSMEG_4879 family |
| **120** | 25% | TIGR03841 | F420_Rv3093c: probable F420-dependent oxidoreductase, Rv3093c family |
| **120** | 25% | PF08242 | Methyltransferase domain |
| **105** | 22% | TIGR01575 | rimI: ribosomal-protein-alanine acetyltransferase |
| **104** | 22% | PF12730 | ABC-2 family transporter protein |
| **96** | 20% | PF05147 | Lanthionine synthetase C-like protein |

**Supplementary Table 2: Comparison of the Dmp chemical shifts observed in 1 (daptide) and hominicin[5].** Data was obtained in 1,1,1,3,3,3-hexafluoroisopropanol-$d_2$ (HFIP-$d_2$) for **1** and $H_2O$ + $D_2O$ for hominicin, respectively. [§]Data not available. *Signals assigned to *N*-methyl groups are not distinguished in the original paper. Dmp: (*S*)-$N_2$,$N_2$-dimethyl-1,2-propanediamine.

| No. | **1 (daptide)** | | **Hominicin** | |
|---|---|---|---|---|
| | $\delta_C$ type | $\delta_H$ multi | $\delta_C$ type | $\delta_H$ multi |
| 1 | 39.1 $CH_2$ | 3.91 overlapped | 39.9 $CH_2$ | 3.46 d (6.3) |
| | | 3.38 d (15.2) | | 3.41 m |
| 2 | 61.7 CH | 3.75 m | 60.9 CH | 3.48 m |
| 3 | 6.9 $CH_3$ | 1.34 brs | 10.9 $CH_3$ | 1.23 d (6.7) |
| 4 | 34.1 $CH_3$ | 2.80 s | 38.2 $CH_3$ | 2.75* s |
| 5 | 41.6 $CH_3$ | 2.97 s | 41.0 $CH_3$ | 2.80* s |
| NH | | N/A[§] | | 8.24 t (5.9) |

**Supplementary Table 3: Comparison of the Ala-23 chemical shifts observed in 1 (daptide) and Ala-20 in hominicin[5].** Data was obtained in 1,1,1,3,3,3-hexafluoroisopropanol-*d*$_2$ (HFIP-*d*$_2$) for **1** and $H_2O$ + $D_2O$ for hominicin, respectively. [§]Data not available. [*]Carbonyl carbon of Ala-22 in **1** and Ile-19 in hominicin.

| No. | **1 (daptide)** | | **Hominicin** | |
|---|---|---|---|---|
| | $\delta_C$ type | $\delta_H$ multi | $\delta_C$ type | $\delta_H$ multi |
| 1’ | 51.0 CH | 4.22 overlapped | 50.0 CH | 4.18 m |
| 2’ | 13.7 $CH_3$ | 1.57 d (6.9) | 16.4 $CH_3$ | 1.29 d (7.4) |
| NH | | N/A[§] | | 8.32 d (5.8) |
| CO | 175.8 C | | 175.5 C | |
| CO[*] | 176.4 C | | 173.3 C | |

**Supplementary Table 4: Indicator strains and corresponding culturing conditions used in the antimicrobial activity screening**. Recipes for mediums used in the bioassay are as the following: lysogeny broth (LB): 10 g/L tryptone, 10 g/L NaCl, and 5 g/L yeast extract; brain heart infusion broth (BHI): 5 g/L beef heart (infusion from 250 g), 12.5 g/L calf brains (infusion from 200 g), 2.5 g/L $Na_2HPO_4$, 2 g/L glucose, 10 g/L peptone, and 5 g/L NaCl; international Streptomyces project-2 medium (ISP2): 10 g/L malt extract, 4 g/L yeast extract, and 4 g/L glucose; ATCC medium 172 (ATCC 172): 10 g/L glucose, 20 g/L soluble starch, 5 g/L yeast extract, 5 g/L N-Z Amine Type A (Sigma C0626), and 1 g/L $CaCO_3$; yeast extract–peptone–dextrose medium (YPD): 10 g/L yeast extract, 20 g/L peptone, and 20 g/L glucose.

| Phylum | Strain | Medium | Temperature (°C) |
|---|---|---|---|
| Bacillota | *Bacillus cereus* TZ417 | LB | 30 |
| | *Bacillus subtilis* ATCC 6633 | LB | 30 |
| | *Lactococcus lactis* CNRZ 481 | LB | 30 |
| | *Staphylococcus epidermidis* 15X154 | LB | 37 |
| | *Staphylococcus aureus* USA300 | BHI | 37 |
| | *Streptococcus mutans* ATCC 25175 | BHI | 37 |
| Actinomycetota | *Micrococcus luteus* ATCC 4698 | LB | 30 |
| | *Streptomyces albus* J1074 | ISP2 | 30 |
| | *Streptomyces lividans* TK24 | ISP2 | 30 |
| | *Streptomyces coelicolor* M145 | ISP2 | 30 |
| | *Microbacterium aurum* B-24210 | ATCC 172 | 30 |
| | *Microbacterium esteraromaticum* B-24213 | ATCC 172 | 30 |
| | *Microbacterium hominis* B-24220 | ATCC 172 | 30 |
| | *Microbacterium ketosireducens* B-24221 | ATCC 172 | 30 |
| | *Microbacterium kitamiense* B-24226 | ATCC 172 | 30 |
| Pseudomonadota | *Escherichia coli* DH5a | LB | 37 |
| | *Enterobacter cloacae* | LB | 30 |
| | *Pseudomonas fluorescens* Pf-5 | LB | 30 |
| | *Pseudomonas putida* mt-2 | LB | 30 |
| Ascomycota | *Saccharomyces cerevisiae* YSG50 | YPD | 30 |
| | *Aspergillus terreus* | YPD | 30 |

**Supplementary Table 5: Oligonucleotides used for cloning and site-directed mutagenesis**. The primers are named for the annealing region and whether the primer was for the forward (F) or reverse (R) direction.

| Name | Nucleotide Sequence (5' to 3' direction) |
|---|---|
| pBE44 F | ttaccaatgcttaatcagtgaggcacc |
| pBE45 R | atctttatagtcctgtcgggtttcg |
| pBE44-DSM15019-ami R | accgtgttcgtcttcccggatctcaacaccggcaacaacaacttatatcgtatggggctg |
| pBE45-DSM15019-ami F | cactcccccgccttcccaccgttctccacacccttctctgacgctcagtggaacgaaaac |
| DSM15019-g1 F | aattaatacgactcactatagggaatttctactgttgtagatccggatctcaacaccggc |
| DSM15019-g1 R | gccggtgttgagatccggatctacaacagtagaaattccctatagtgagtcgtattaatt |
| DSM15019-g2 F | aattaatacgactcactatagggaatttctactgttgtagatccaccgttctccacaccc |
| DSM15019-g2 R | gggtgtggagaacggtggatctacaacagtagaaattccctatagtgagtcgtattaatt |
| pBE44-B3501-ami R | ggtcacctgcgcttcttcgaacatgtccgagccggtacaacaacttatatcgtatggggctgacttc |
| pBE45-B3501-ami F | cgcacagaagctttgaaggtcgaaagggatagggatatcgacgctcagtggaacgaaaac |
| B3501-g1 F | aattaatacgactcactatagggaatttctactgttgtagatttcgaacatgtccgagcc |
| B3501-g1 R | ggctcggacatgttcgaaatctacaacagtagaaattccctatagtgagtcgtattaatt |
| B3501-g2 F | aattaatacgactcactatagggaatttctactgttgtagataaggtcgaaagggatagg |
| B3501-g2 R | cctatccctttcgaccttatctacaacagtagaaattccctatagtgagtcgtattaatt |
| pSET (XbaI)-mpaB F | gacggccagtgccaagcttgggctgcaggtcgactctagacgccgcccgaaacgagagaa |
| mpaP R | gcgacgccgatcgggtcgag |
| mpaM F | tcctcacgacctcccgcgcc |
| Yeast-mpaT3 R | ttatagcacgtgatgaaaaggacccaggtggcacttttcgccaggaatcctacgtcgcct |
| mpaT3-Yeast F | acgtgtcgtagcgcttcaggaggcgacgtaggattcctggcgaaaagtgccacctgggtc |
| pSET (EcoRI)-Yeast R | cacacaggaaacagctatgacatgattacgaattcgcatcacagtatcgtgatgacagtg |
| Yeast-mpaP R | tagcacgtgatgaaaaggacccaggtggcacttttcgtcatcctctctcctcctcgctcg |
| mpaP-Yeast F | cccgctggcagctctcacgagcgaggaggagagaggatgacgaaaagtgccacctgggtc |
| Yeast-mpaM R | ttatagcacgtgatgaaaaggacccaggtggcacttttcgtcatcgccggccccttccat |
| mpaM-Yeast F | cacgacctcccgcgccggggatggaaggggccggcgatgacgaaaagtgccacctgggtc |
| Yeast-mpaA1 R | tatagcacgtgatgaaaaggacccaggtggcacttttcggcgtccctccttcctttctcc |
| mpaA1-Yeast F | cgtccggcagcacccgatcggagaaaggaaggagggacgccgaaaagtgccacctgggtc |
| BbsI-mpaA1 F | acgtgaagacaaaatgatgaatgaatccacagaaatgtcgatccg |
| BbsI-mpaA1 R | acgtgaagacaaaccgtcaggtcgctgccgctgc |
| mpaA1 E13A R | ttccatcgggtccagcgcttgaaagcggatcga |
| mpaA1 E13A F | tcgatccgctttcaagcgctggacccgatggaa |
| mpaA1 L14A R | ggcttccatcgggtccgcctcttgaaagcggat |
| mpaA1 L14A F | atccgctttcaagaggcggacccgatggaagcc |
| mpaA1 D15A R | gggggcttccatcggcgccagctcttgaaagcg |
| mpaA1 D15A F | cgctttcaagagctggcgccgatggaagccccc |
| mpaA1 P16A R | actgggggcttccatcgcgtccagctcttgaaa |
| mpaA1 P16A F | tttcaagagctggacgcgatggaagcccccagt |
| mpaA1 M17A R | ccaactgggggcttccgccgggtccagctcttg |
| mpaA1 M17A F | caagagctggacccggcggaagcccccagttgg |
| mpaA1 E18A R | atcccaactgggggccgccatcgggtccagctc |
| mpaA1 E18A F | gagctggacccgatggcggcccccagttgggat |

| | |
|---|---|
| mpaA1 P20A R | aaaggaatcccaactcgcggcttccatcgggtc |
| mpaA1 P20A F | gacccgatggaagccgcgagttgggattccttt |
| BbsI-mpaA1 Ser R | acgtgaagacaaaccgtcagctcgctgccgctgcgccaatac |
| BbsI-mpaA1 Cys R | acgtgaagacaaaccgtcagcacgctgccgctgcgccaatac |
| BbsI-mpaA1 Asp R | acgtgaagacaaaccgtcaatccgctgccgctgcgccaatac |
| BbsI-mpaA1 Asn R | acgtgaagacaaaccgtcagttcgctgccgctgcgccaatac |
| BbsI-mpaA1 Glu R | acgtgaagacaaaccgtcattccgctgccgctgcgccaatac |
| BbsI-mpaA1 Gln R | acgtgaagacaaaccgtcactgcgctgccgctgcgccaatac |
| BbsI-mpaA1 Gly R | acgtgaagacaaaccgtcagcccgctgccgctgcgccaatac |
| BbsI-mpaA1 Ala R | acgtgaagacaaaccgtcacgccgctgccgctgcgccaatac |
| BbsI-mpaA1 Phe R | acgtgaagacaaaccgtcaaaacgctgccgctgcgccaatac |
| BbsI-mpaA1 Lys R | acgtgaagacaaaccgtcatttcgctgccgctgcgccaatac |
| BbsI-mpaA1 Arg R | acgtgaagacaaaccgtcaacgcgctgccgctgcgccaatac |
| BbsI-mpaA1 Val R | acgtgaagacaaaccgtcacaccgctgccgctgcgccaatac |
| BbsI-mpaA1 +44A R | acgtgaagacaaaccgtcaggtcgccgctgccgctgcgccaatac |
| BbsI-mpaA1 -43A R | acgtgaagacaaaccgtcaggttgccgctgcgccaataccga |
| BbsI-mpaA1 +45A R | acgtgaagacaaaccgtcacgcggtcgctgccgctgcgccaa |
| BbsI-mpaA1 G39P R | acgtgaagacaaaccgtcaggtcgctgccgctgccggaataccgatc |
| BbsI-mpaA1 G37P R | acgtgaagacaaaccgtcaggtcgctgccgctgcgccaatcgggatcaggaca |
| BbsI-mpaA1 V34P R | acgtgaagacaaaccgtcaggtcgctgccgctgcgccaataccgatcagcggaagcgctc |
| BbsI-mpaA1 G31P R | acgtgaagacaaaccgtcaggtcgctgccgctgcgccaataccgatcaggacaagcgccgggatgac acct |
| BbsI-mpaA1 G28P R | acgtgaagacaaaccgtcaggtcgctgccgctgcgccaataccgatcaggacaagcgctccgatgac cggttgataaaag |

**Supplementary Table 6: DNA sequences of the *E. coli* codon-optimized *mpa* genes**.

| Gene | Nucleotide Sequence (5' to 3' direction) |
|---|---|
| *mpaA1* | atgaatgaatccacagaaatgtcgatccgctttcaagagctggacccgatggaagcccccagttgggattccttttatc<br>aaggtgtcatcggagcgcttgtcctgatcggtattggcgcagcggcagcgacctga |
| *mpaA2* | atgatgaatgccgcgctgcctgccttagagtttacggagctggaagctatggatgcgccgggcgatgcagatgattata<br>tccgtggcttcgctgtgggcgttatcatcgggattctcgccctgacctaa |
| *mpaA3* | atgcaatcaacatcccttgaatttctggaattagaacagatcgacactccattggaatggtgggaacatgcgagttaca<br>ttattgcaattattggtggtgcagctgccattgcgacgtga |
| *mpaB* | atgggaacctcagctacgaccgcacgccaagacgacggctcaaccggcgaagatccgttattaggccctcatgtccttc<br>gcgcatgggtaacgggcggccacgcggcccgtggtcgcgttttcctgcgtgaggcagaaacgggtgcggaagcgttagc<br>tgcagcggctgatccggaagatgtcattctggagccagccggttctgcgcctcgtccaggagcagcccgtgtcgttggg<br>tatggcggacgcctggccgatttgggtgatgaactctttctcggcgaacgcggagtggagctgcaagactatatcgccg<br>ccagtttcgttcagatcgttggtcccacggccatgcgtttctttgatgaggcttcctggcgtgccttcctcgatgatgc<br>cgatctcgcacgtggcaccggcgtgtttgccgcggcaatgctggacccacgtgtcttattggccgaccgctcagcactg<br>gcgcgcccgcaggaagtggagacgccacgtgccgtacgcatcgacggggacggtggtgtgcacttaggcctgcagggcg<br>aggcaattggtcatgttgacgatcttccagacgccctgggttgggcgttacctcgtgcttcaatgcttggtggtgttcc<br>tggctcggccgatctcgttgccgaacttgatacgcgtccatggttacgccgctacctcgatgctgcggaccttcgtaaa<br>atgttacgcctgccgaatggcaccgcacgtattgcaggctacggttgggcattactggatgacgatctggcggaagcgg<br>aagcccttgcggatgaccctttcttttagataccgccgaaggttttctgttggcggatgtgcgtacccttcgccgtca<br>cttactttcgcctttaacagccgcggtggtggatgccatccagactagtagcacgcgtggtcgtgctgcggcacgcgtg<br>gctcgcgcctgtggcgtgccggatgcgcacgcccgccatttgtgtttggaagcgttagcagctctgggcgttcacttag<br>gggcacgcaccgaggacgcatctggtgatggggcacgttga |
| *mpaC* | atgactgtacccttcacagcccgcgttccgaccttatggcacggcagtcgcgcacgcgagcgcatggtaggtctgactg<br>cgggccgtgataccctggtgctggctgacgcggcggtagcgcatcccttgcctactcctcaatcgggccgcactatgac<br>cgtcgacgccgccgcggtcgacgagttgggaattcttctgatcgcggaagaactggtccgtcgtccgcctgaagtgatc<br>gtggcggtcggtggcggctcagttctggatgcctctaagattgccgcgctggctctctctcctgggcgcttgcttgatt<br>tcgcacttcgtcacgcagccacgtctgctcttaccatcttaccggacgctcccccgccagtggacattgtggcgatgcc<br>aacgacccttggaacctcttcggaaaccaacagcgtggccatccttcgtaatcgttgtggcgctcgtctgcttgtggga<br>cgtcccttacgtccgcgccacgcagtcttggacccggaccagctggccactttgacccctgcagcggttcgtgagggcg<br>ctcttgaggcgtttctccgcgtcgccggagcgagtacctcaccacgtacagcccgcgccgaccgtgatgcgcgtatctt<br>aggccgcgcgttgctggaagcggctgcacgtgatgtaatgtctacgaccggtcgtctccgtcttgctcgccttagcgcg<br>gccactcagcgcactgccgctctccgtggccctgacccgtattctgcacgccactggtacttggccaatgaggtagcct<br>tccacctgggagtgcgcaaaatgacggcaactgcggcagtgattgcagccgtatggcaacgcatcggtgcgggcgaccc<br>gcgctggggtgaccgtgcgagcttggacgcattttgggctggagttgcagcaggaacggcgttacctcgtgatcctgca<br>gctggaatcgcggtagtggtagatcagggcgggttacccagcccaccacgccccagcgcgcgtgagattgatcgtattg<br>cggacgcagtcgaacgtgcgtggggagatcatcgccctatgttgcgcggcattcgcgccgctgatgtacgtgcagtttt<br>gcacgactcccgctggagcccttccccggtaggcgatcgtcgtcctcgtgtggcagagcgtcgttga |
| *mpaD* | atgagcgttccgcatgccctttggacctctatgcttccagcagacaccgcatttccgcccgaccgcgtcgccatcgggg<br>ccgcaggccaccgtttgttctacgcagacgggtctgatcgtttatgcgctacaagtgggttgtggaacgtaccgttggg<br>atttgggaatccagcggtcaccgaggctgtctcacgcgcgactcgtgacgcgtcctacctgagcttattccgtgctccc<br>caccgttacgcagaggatgcggcggatgcgttaattgcgctcgcaggaagcgaccgttaccgccgcgtcatctttagca<br>ctagtggcggcgccgcaaatgatgccgctatgaaactcgcccgtcaatattgggcacaacaaggtgctggtagccgttc<br>tttagtagttggtttacgcggtagctatcatggtacgatgtatggctcccacgttctttcgggcgacgatctgctgcag<br>tcggtgtatggcgtcgaccgtcgtacggtacgccacgtaagtcacacggatgaaggggctgagctggaagcgttgctgg<br>agcgcgaaggttcccgtgtcgccagtgtcgttgtcgagccggtactgggttcaggtgcccatgcgctttctgctgcatt<br>cgtgcgtcgcttgctcgagttgcgcgaacgccatgggtttctgttagttgcggacgaggtagccaccgggttcggtcgt<br>acgggcgcgatgttcgccactgaaggttgggcagccgctccagatgtccttcttgtatcaaaggcgctcacgaatggag<br>ctatgggcgcagcagcgctcttggtgggtgatcgcattgcaaccgcctttgcgcgtggcggctggacatttgttcacgg<br>tgagacacaagccgggacgcctgcctgtgcagcggctgtccttgcggttatcgacgagcttcgtcgtattgacgtcgag<br>gcgaccactcaagcgctggctagcacgttacatcgtttggcccgcacgtggcaagaggacggcatcgtgaccgatgtta<br>gcggaagtggttgcttcgtgggtgtaggattacgccgtcctgacggttcagctctgagcggaaccgacgtgctggaagt<br>ggttagtgccattgcggattccggcgttgttgtacatcctggcccgtcatcatttcagttaatccctggctacggtttt<br>acacctgcagaggtagcggaaactgaccgcgcggtgcgtggcggcttggcacgtgcccgtgaggcggtggcatga |
| *mpaM* | atgacctcgttgattaccgagactgtcgccgcccgtctggaccttgtgggtgctaccgcccgtgcgcaagacttgtatg<br>caggcgcgggtacagacttttatgatcgcctcgtaggtcccgatcgtgccgaaattcgtgaggtactgggtttagccca<br>caccgcacccggtccggtactggacctcgcagcgggttctggccgtttgacaattcctcttgcgcgtagtggccaccgc<br>gtgacagccgtggaccttagtgcggatatgttagcccgtttgcgtggcgcagtcccgcatggcgccaccgtagaatgcg<br>ttgtggctgatatgcgtgacttagcacttggcggacacttcggcctggtagttctcggagcaacgtcgatcaccctgct<br>cgaccgcgaagatcgcgcacgcctctacgccggtgtgcgtcgtcatcttgcatcagcaggtgtctttgcattgaccatt<br>gcggacggtgtcagtgcagacacgcttgcgatcccaacggaccgcgaaattacggttcctggttccgcaggtgatgagc<br>cttacttattcgcacaacaaatcgaggatgatggggctgcgcgtcttgtaaactgggtccgcctggctgacatcgcccc<br>acgtgcagaagttccggtgttaacatcacgcttgcacgtcttaaccccagaggtcctggctggcgagttagtagcagcc<br>gggtttacagcacccgagcgctcgccggttcgcactccggcaggtgttgagattttgttgttaacgacctcgcgcgccg<br>gggacggacgcggtcgtcgctaa |

**Supplementary Table 7: Naming scheme for daptide biosynthetic genes.** Custom pHMM models for daptide detection are shown in red.

| Gene Name | Function | pHMM annotations for identification |
|---|---|---|
| **DapA** | Precursor peptide | N/A |
| **DapB** | DUF-RRE fusion protein | Actino_DapB_RRE, Bacill_DapB_RRE |
| **DapC** | Oxidative decarboxylase (alcohol dehydrogenase) | PF00465, PF13561 |
| **DapD** | Aminotransferase | PF00202 |
| **DapM** | Methyltransferase | PF13649, PF08241, PF13489, PF08242, *etc*. |
| **DapP** | Intramembrane protease | PF02163, Actino_DapP_RRE, Bacill_DapP_RRE |
| **DapT** | ABC transporter | TIGR01188, TIGR03740, PF00528, *etc*. |

**Supplementary Note: Algorithm for pairing of RRE families with precursor peptide families.**

```
import argparse
import sys
import os
import re

def __main__():

    input_file = open(RRE_ALL_BY_ALL_BLAST_FILE, 'r')
    input_precursor_file = open(PRECURSOR_ALL_BY_ALL_BLAST_FILE, 'r')
    enzyme_edge_data = input_file.read().split("\n")
    precursor_edge_data = input_precursor_file.read().split("\n")
    input_file.close()
    input_precursor_file.close()
    cutoff = 50
    print("Starting enzyme edge filtering...")

    enzyme_edges_withAS = []
    for line in enzyme_edge_data:
        values = line.split()
        try:
            enzyme_edges_withAS.append([values[0], values[1], float(values[11])])
        except:
            pass

    print("Starting ID mapping...")

    precursor_edges_withids = []
    for line in precursor_edge_data:
        try:
            node1, node2, BS = re.split("\t", line)
            if float(BS) > 10:
                precursor_edges_withids.append([node1, node2])
        except:
            pass

    open("temp.tsv", 'w').close()

    enzyme_cluster_generate(output_file_name=”temp.tsv", enzyme_edges_withAS=enzyme_edges_withAS,
cutoff=cutoff, precursor_edges_withids=precursor_edges_withids)

def enzyme_cluster_generate(output_file_name, enzyme_edges_withAS=[], cutoff=1000,
precursor_edges_withids=[]):

    print("Starting enzyme graph generation... cutoff = ", cutoff)

    enzyme_edges = []
    for edge in enzyme_edges_withAS:
        if edge[2] > cutoff and edge[0] < edge[1]:
            enzyme_edges.append([edge[0], edge[1]])
    enzyme_clusters = sorted(connected_components(enzyme_edges), key=len, reverse=True)

    print("Starting precursor graph generation...")

    for se in enzyme_clusters:
        print("Cutoff: ", cutoff, "Enzyme cluster size: ", len(se))
        if len(se) < 30:
            print("Too small:", len(se))
            break
        precursor_edges = []
        new_precursor_edges_withids = []
        for edge in precursor_edges_withids:
            if edge[0].split("_")[0] in se and edge[1].split("_")[0] in se:
                precursor_edges.append([edge[0], edge[1]])
            new_precursor_edges_withids.append(edge)
        edge_count = 0
        for enzyme_edge in enzyme_edges:
            if enzyme_edge[0] in se and enzyme_edge[1] in se:
                edge_count += 1
        enzyme_connectivity = (((2 * edge_count) / float(len(se))) / float((len(se)-1)))
        print("Enzyme connectivity: " + str(enzyme_connectivity))

        precursor_clusters = sorted(connected_components(precursor_edges), key=len, reverse=True)
        try:
            x = precursor_clusters[0]
        except:
            return
        if len(x) < 5:
            return
        print("Precursor cluster size: ", len(x))
        cluster_found=False

        if len(x) > 0.1 * len(se) and enzyme_connectivity>0.2:
```

```
            print("Precursor cluster size: " + str(len(x)) + "    Enzyme cluster size: " + str(len(se)) + "
BS cutoff: " + str(cutoff))
            cluster_found=True
            precursor_edge_count = 0
            for z in precursor_edges:
                if z[0] in x and z[1] in x and z[0] != z[1]:
                    precursor_edge_count+=1
            precursor_connectivity = precursor_edge_count / len(x) / (len(x)-1)
            print("Precursor connectivity: " + str(precursor_connectivity))

        new_enzyme_edges=[]
        real_node_list = []
        for edge in enzyme_edges_withAS:
            if edge[0] in se and edge[1] in se:
                new_enzyme_edges.append(edge)
                if edge[0] not in real_node_list:
                    real_node_list.append(edge[0])
                if edge[1] not in real_node_list:
                    real_node_list.append(edge[1])
        if cluster_found==False:
            enzyme_cluster_generate(output_file_name=output_file_name,
enzyme_edges_withAS=new_enzyme_edges, cutoff=cutoff+10,
precursor_edges_withids=new_precursor_edges_withids)
        else:
            precursor_sequences = open(PRECURSOR_FASTA_FILE, 'r').read().split(">")
            output_file = open(output_file_name, 'a')
            output_file.write(str(cutoff) + "\t" + ",".join(se) + "\t" + str(len(se)) + "\t" +
str(enzyme_connectivity)+"\t")
            for precursor in precursor_sequences:
                header = precursor.split("\n")[0]
                if (header) in x:
                    output_file.write(">" + header + " " + "".join(precursor.split("\n")[1:]) + ",")
                continue
            output_file.write("\t" + str(len(x)) +"\t"+str(precursor_connectivity) + "\t" +
",".join(real_node_list) + "\t" + str(len(real_node_list)) + "\n")
            output_file.close()

def connected_components(pairs):
    components = []
    for a, b in pairs:
        a_num = -1
        b_num = -2
        for i, component in enumerate(components):
            if a in component:
                a_num = i
            if b in component:
                b_num = i
            if a_num != -1 and b_num != -2:
                break
        if a_num == -1:
            if b_num == -2:
                components.append([a,b])
                #print(a,b)
            else:
                components[b_num].append(a)
                #print(a,b,components[b_num])
        elif b_num == -2:
            components[a_num].append(b)
            #print(a,b,components[a_num])
        elif a_num != b_num:
            components[a_num].extend(components[b_num])
            components[b_num] = []
    return components

if __name__=="__main__":
    __main__()
```

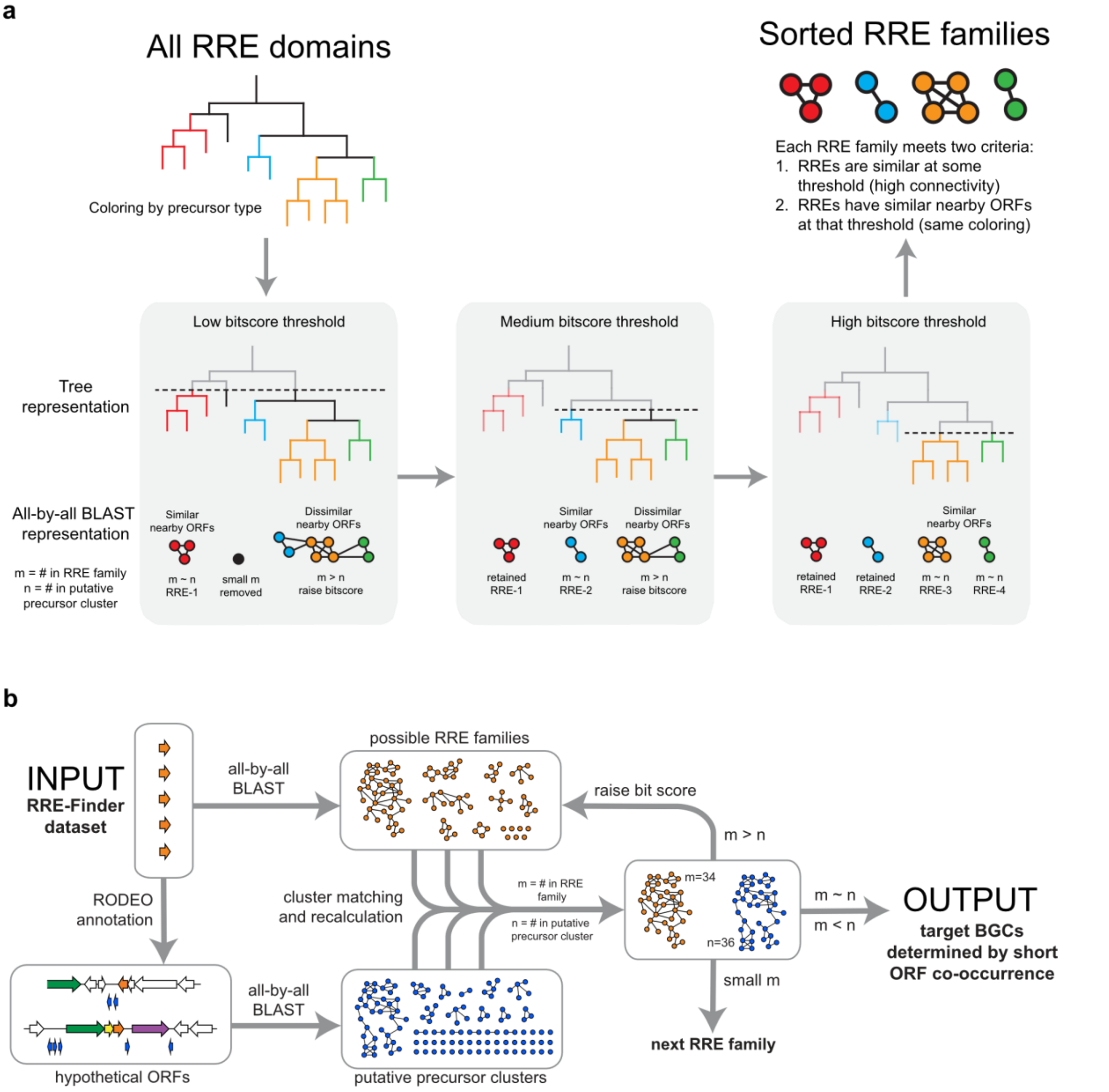

**Supplementary Figure 1: Generalized workflow for bioinformatic analysis of the RRE-Finder exploratory mode dataset**. The comprehensive description for this workflow is in the Methods section, and code used for implementation is on Github (https://github.com/the-mitchell-lab/rodeo2) and in the **Supplementary Note**. Phylogenetic trees, sequence similarity networks, and biosynthetic gene clusters depicted in this figure are stylized. (**a**) Logical basis for finding new RRE-dependent RiPP pathways. Analysis of RRE domain clusters is performed iteratively with increasing bitscore thresholds. Examination of each cluster for high similarity nearby open-reading frames (ORFs) allows categorization of RRE families based on sequence similarity of the RRE domains and co-occurrence with high-likelihood precursor peptides. (**b**) Workflow of bioinformatics and algorithmic logic. RRE domains predicted by RRE-Finder were annotated by RODEO to identify hypothetical precursors in intergenic ORFs. The RRE domains were sorted into families using an all-by-all BLAST search and the predicted ORFs were also sorted into clusters. For each RRE family, putative precursor peptide clusters were filtered for ORFs in the same genomic neighborhood, and the sizes of the families were compared. Small RRE families were discarded from the algorithm. Cases where the size of the largest precursor peptide cluster matched the RRE family were output from the algorithm for further analysis. All other cases were re-entered into the algorithm with a higher bitscore threshold for RRE family generation.

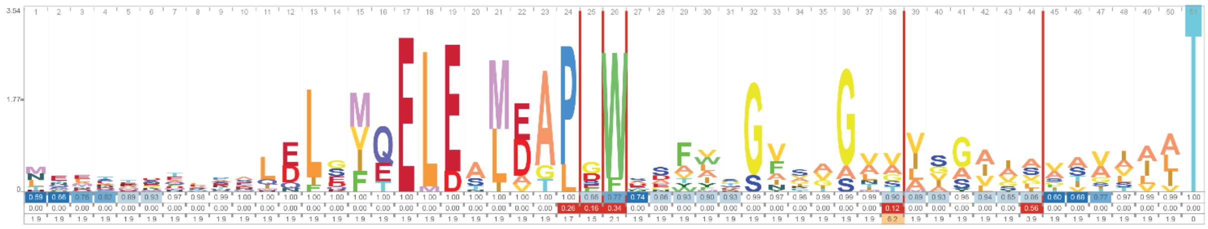

**Supplementary Figure 2: Daptide precursor peptide sequence logo**. The daptide precursor peptide pHMM ($n$ = 184) was generated from precursor peptides identified by the workflow in **Supplementary Figure 1**. The pHMM was converted to a sequence logo using Skylign[6] with the information content above background setting, and the direct output is provided. The invariant Thr residue at the C-terminus is evident (cyan color), along with highly conserved features in the N-terminal leader region.

**a**

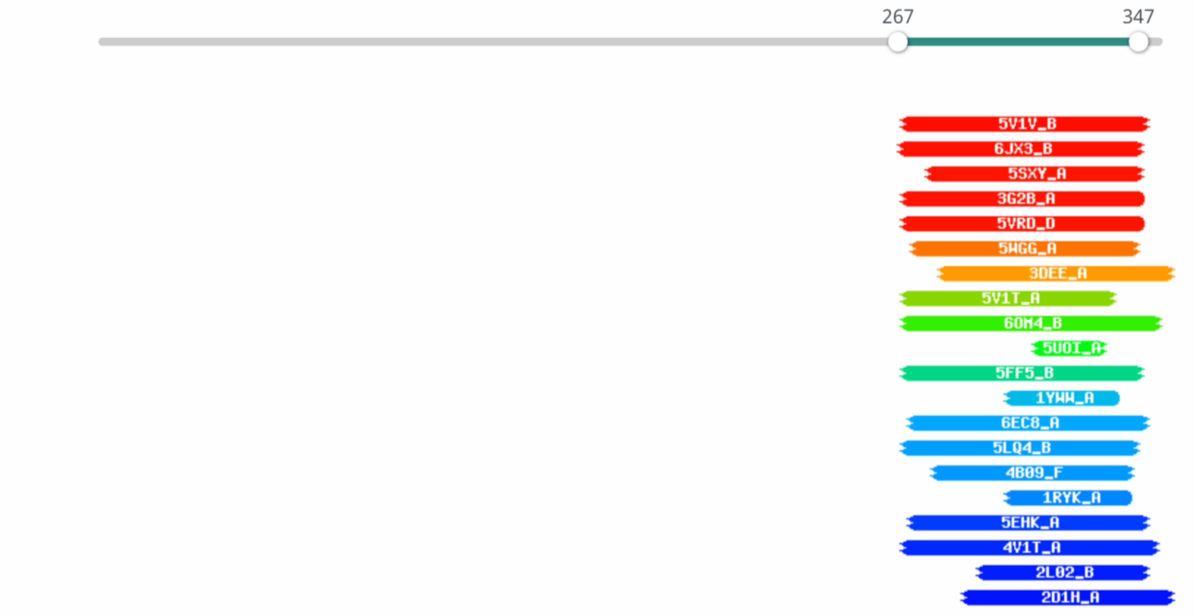

**b**

Hitlist

Show 25 Entries Search:

| Nr | Hit | Name | Probability | E-value | Score | SS | Aligned cols | Target Length |
|---|---|---|---|---|---|---|---|---|
| 1 | 5V1V_B | TbiB1; Lasso Peptide, RiPP, Peptide Binding, PROTEIN BINDING; 1.35A {Thermobaculum terrenum (strain ATCC BAA-798 / YNP1) | 98.85 | 1e-7 | 72.32 | 12.4 | 81 | 91 |
| 2 | 6JX3_B | TfuB1; lasso peptide, RRE, PEPTIDE BINDING PROTEIN; 1.7A {Thermobifida fusca} | 98.64 | 8e-7 | 67.55 | 11.7 | 80 | 99 |
| 3 | 5SXY_A | Bifunctional coenzyme PQQ synthesis protein C/D; RiPP RRE peptide scaffolding, CHAPERONE; NMR {Methylobacterium extorque | 98.35 | 0.0000077 | 61.76 | 10.3 | 69 | 94 |
| 4 | 3G2B_A | Coenzyme PQQ synthesis protein D; helix-turn-helix, PQQ biosynthesis, BIOSYNTHETIC PROTEIN; 1.66A {Xanthomonas campestri | 98.34 | 0.0000082 | 62.04 | 10.3 | 78 | 95 |
| 5 | 5VRD_D | Bifunctional coenzyme PQQ synthesis protein C/D; PQQ, oxidase, alpha-helical bundle, OXIDOREDUCTASE; 2.85A {Methylobacte | 98.33 | 0.0000043 | 82.98 | 11.1 | 78 | 392 |

**Supplementary Figure 3: HHpred results for MpaB**. (**a**) Depicted annotation of MpaB (RefSeq ID: WP_060922562.1) analyzed using HHpred[7] with the PDB_mmCIF70_12_Oct database. The *N*-terminal domain of the protein is not annotated by HHpred, while the C-terminal domain is annotated with RREs. (**b**) Hitlist table of HHpred analysis.

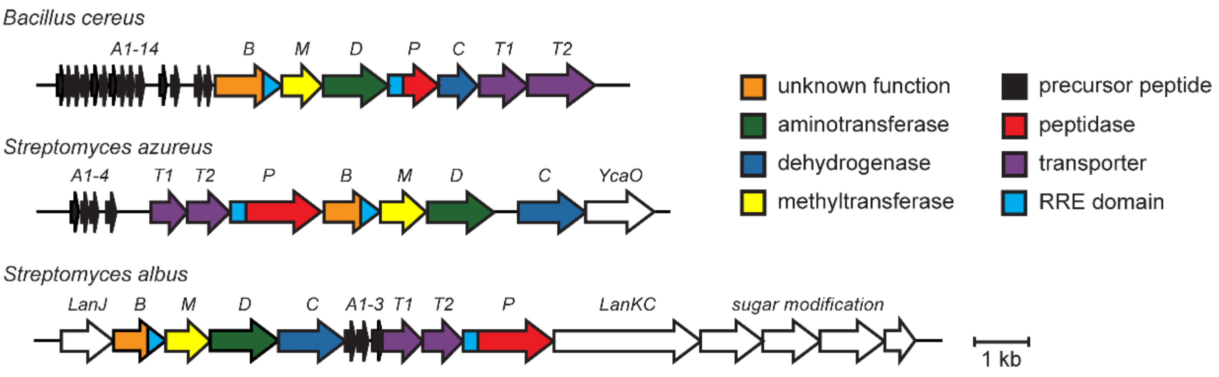

**Supplementary Figure 4: Alternative identified daptide BGCs**. Depicted BGCs are from two bacterial phyla (Bacillota and Actinomycetota) with differing genomic architectures, tailoring genes, and precursor peptide count.

**a**

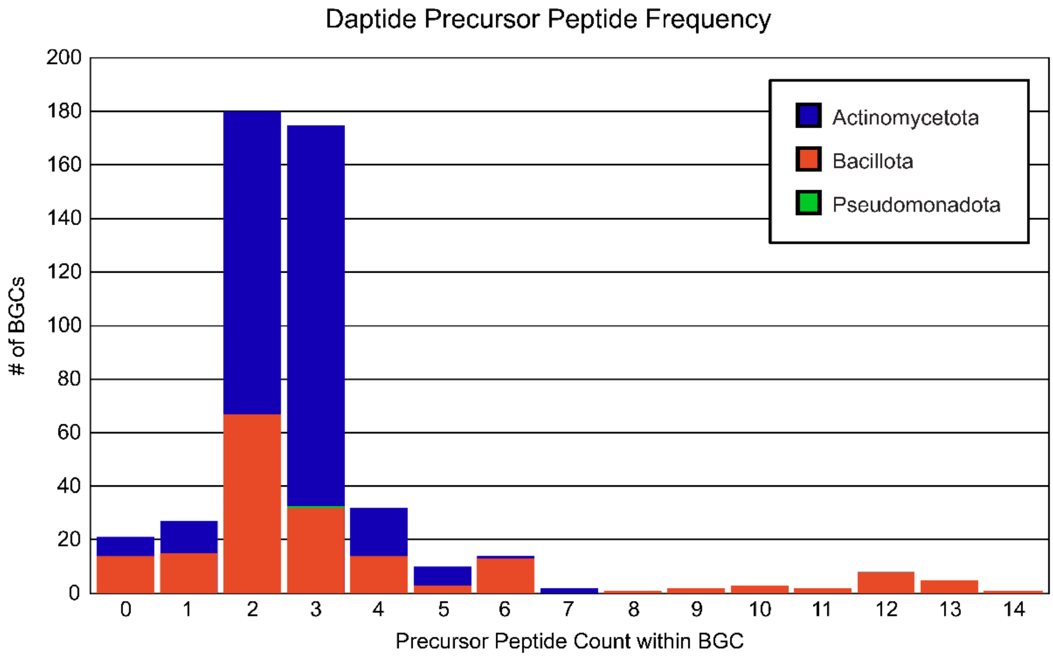

**b**

Precursor Peptide Count within BGC

| | 0 | 1 | 2 | 3 | 4 | 5 | 6 | 7 | 8 | 9 | 10 | 11 | 12 | 13 | 14 | Total |
|---|---|---|---|---|---|---|---|---|---|---|---|---|---|---|---|---|
| **Actinomycetota** | 7 | 12 | 113 | 142 | 18 | 7 | 1 | 2 | | | | | | | | 302 |
| **Bacillota** | 14 | 15 | 67 | 32 | 14 | 3 | 13 | | 1 | 2 | 3 | 2 | 8 | 5 | 1 | 180 |
| **Pseudomonadota** | | | | 1 | | | | | | | | | | | | 1 |
| **Total** | 21 | 27 | 180 | 175 | 32 | 10 | 14 | 2 | 1 | 2 | 3 | 2 | 8 | 5 | 1 | 483 |

**Supplementary Figure 5: Daptide precursor peptide frequency within a BGC**. (**a**) Histogram of daptide precursor peptides per BGC. (**b**) Numerical values used to generate the histogram are given in table format. Each BGC that lacked an identified precursor peptide originated from a short contig, and the BGC was presumed to be incomplete.

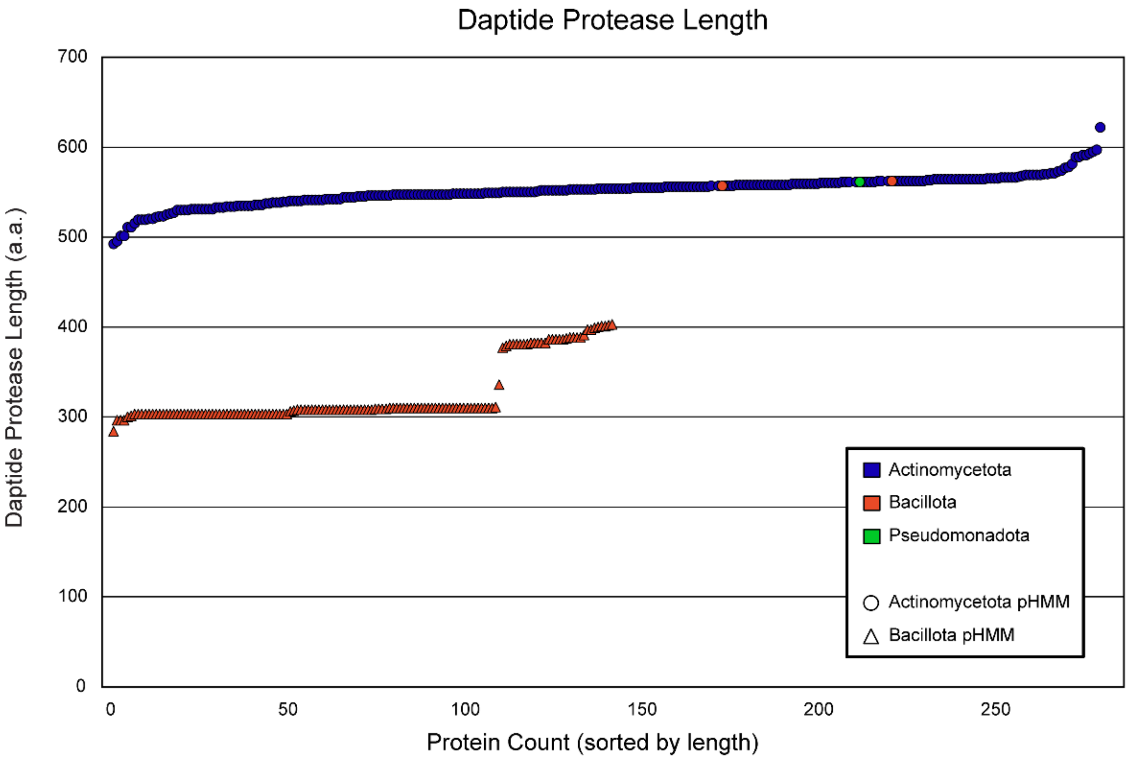

**Supplementary Figure 6: Taxonomic comparison of daptide protease lengths**. All full-length hits for the Actino_DapP_RRE and Bacill_DapP_RRE found using RODEO and within daptide BGCs were compiled. Protein lengths in amino acids (a.a.) were sorted into two series based on which pHMM best retrieved each protein (circle and triangle) prior to sorting by length. The color coding is based on GenBank taxonomic classification.

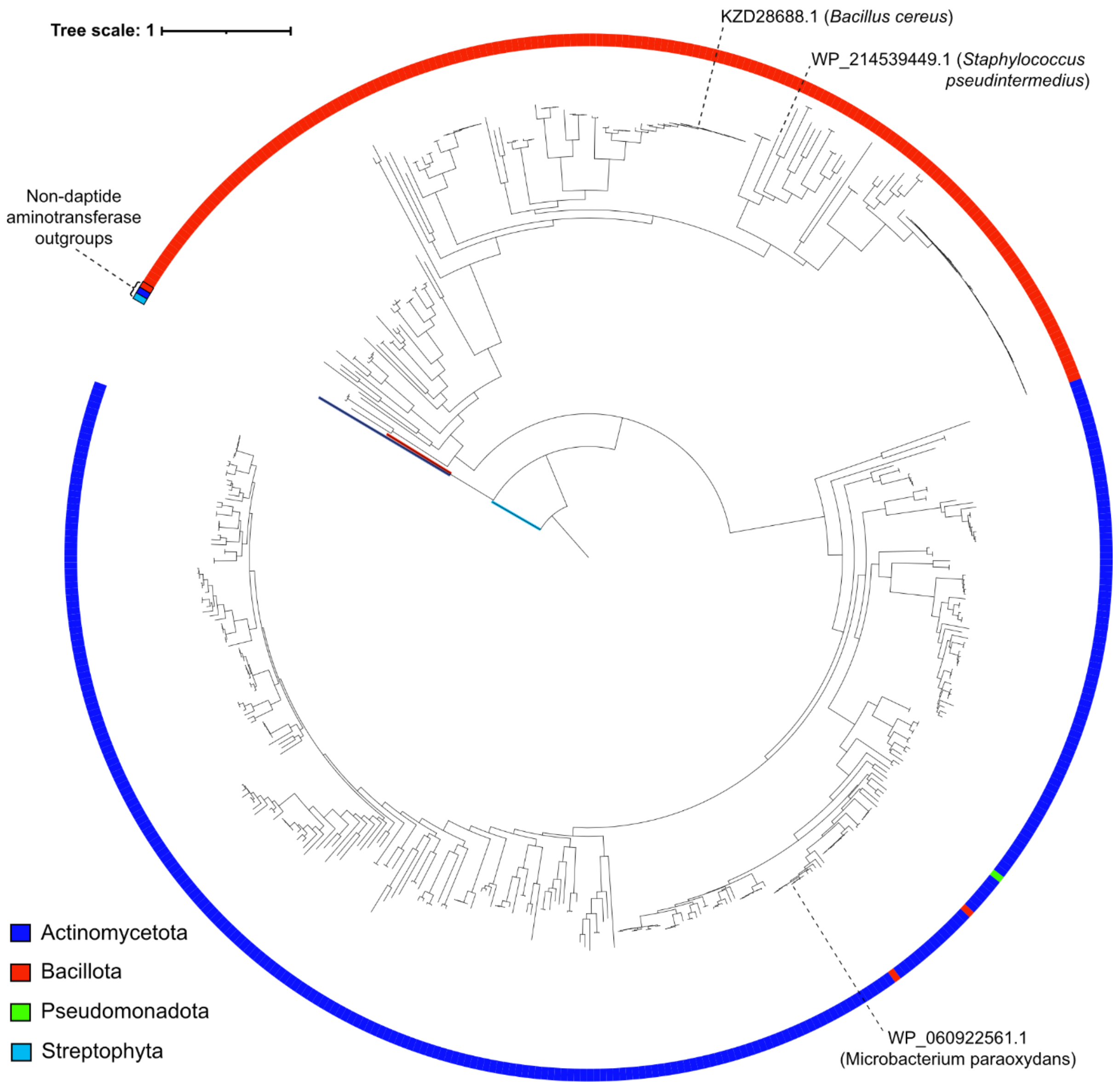

**Supplementary Figure 7: Phylogenetic tree of aminotransferases**. Maximum-likelihood phylogenetic tree of all 483 identified daptide aminotransferases and three outgroup aminotransferases (from aminotransferase class-III[8]). The tree is rooted using gamma-aminobutyrate transaminase 3 (UniProt/GenBank CDS identifiers: Q84P52/BAG16484.1 - Streptophyta). Two additional outgroups are included (P9WQ81/CCP44332.1 - Actinomycetota and P53555/AAB17458.1 - Bacillota) that perform related reactions in biotin biosynthesis.

**a**

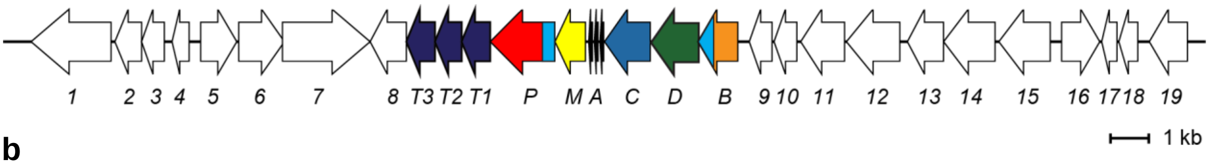

**b**

| Gene | NCBI Accession | Description (per NCBI listing) |
|---|---|---|
| *orf1* | WP_060923289.1 | Phosphate acetyltransferase |
| *orf2* | WP_157547021.1 | Hypothetical protein |
| *orf3* | WP_060922551.1 | Rhomboid family intramembrane serine protease |
| *orf4* | WP_157547022.1 | Hypothetical protein |
| *orf5* | WP_060922553.1 | MoxR family ATPase |
| *orf6* | WP_060922554.1 | DUF58 domain-containing protein |
| *orf7* | WP_060922555.1 | DUF3488 and transglutaminase-like domain-containing protein |
| *orf8* | WP_060922556.1 | Rhodanese-related sulfurtransferase |
| *mpaT3* | WP_083370907.1 | Hypothetical protein |
| *mpaT2* | WP_060922557.1 | Hypothetical protein |
| *mpaT1* | WP_157547023.1 | ATP-binding cassette domain-containing protein |
| *mpaP* | WP_060922558.1 | Hypothetical protein |
| *mpaM* | WP_060922559.1 | Class I SAM-dependent methyltransferase |
| *mpaA3* | N/A[a] | Precursor peptide |
| *mpaA2* | N/A[a] | Precursor peptide |
| *mpaA1* | N/A[a] | Precursor peptide |
| *mpaC* | WP_060922560.1 | Iron-containing alcohol dehydrogenase |
| *mpaD* | WP_060922561.1 | Aminotransferase class III-fold pyridoxal phosphate-dependent enzyme |
| *mpaB* | WP_060922562.1 | Hypothetical protein |
| *orf9* | WP_174521438.1 | GNAT family N-acetyltransferase |
| *orf10* | WP_060922563.1 | Phosphatase PAP2 family protein |
| *orf11* | WP_060922564.1 | Cystathionine gamma-synthase |
| *orf12* | WP_025104232.1 | Cystathionine beta-synthase |
| *orf13* | WP_025104231.1 | Carbohydrate ABC transporter permease |
| *orf14* | WP_025104230.1 | Sugar ABC transporter permease |
| *orf15* | WP_060923551.1 | Bacterial extracellular solute-binding protein |
| *orf16* | WP_083371072.1 | LacI family transcriptional regulator |
| *orf17* | WP_025104227.1 | Ribosomal protein |
| *orf18* | WP_025104226.1 | Ribosomal protein |
| *orf19* | WP_157547024.1 | Hypothetical protein |

**Supplementary Figure 8: Annotation of the cloned region from *M. paraoxydans* DSM 15019**. (**a**) Gene arrangement of the cloned region. (**b**) Accession numbers and annotation of corresponding genes in NCBI. [a]Not annotated as gene.

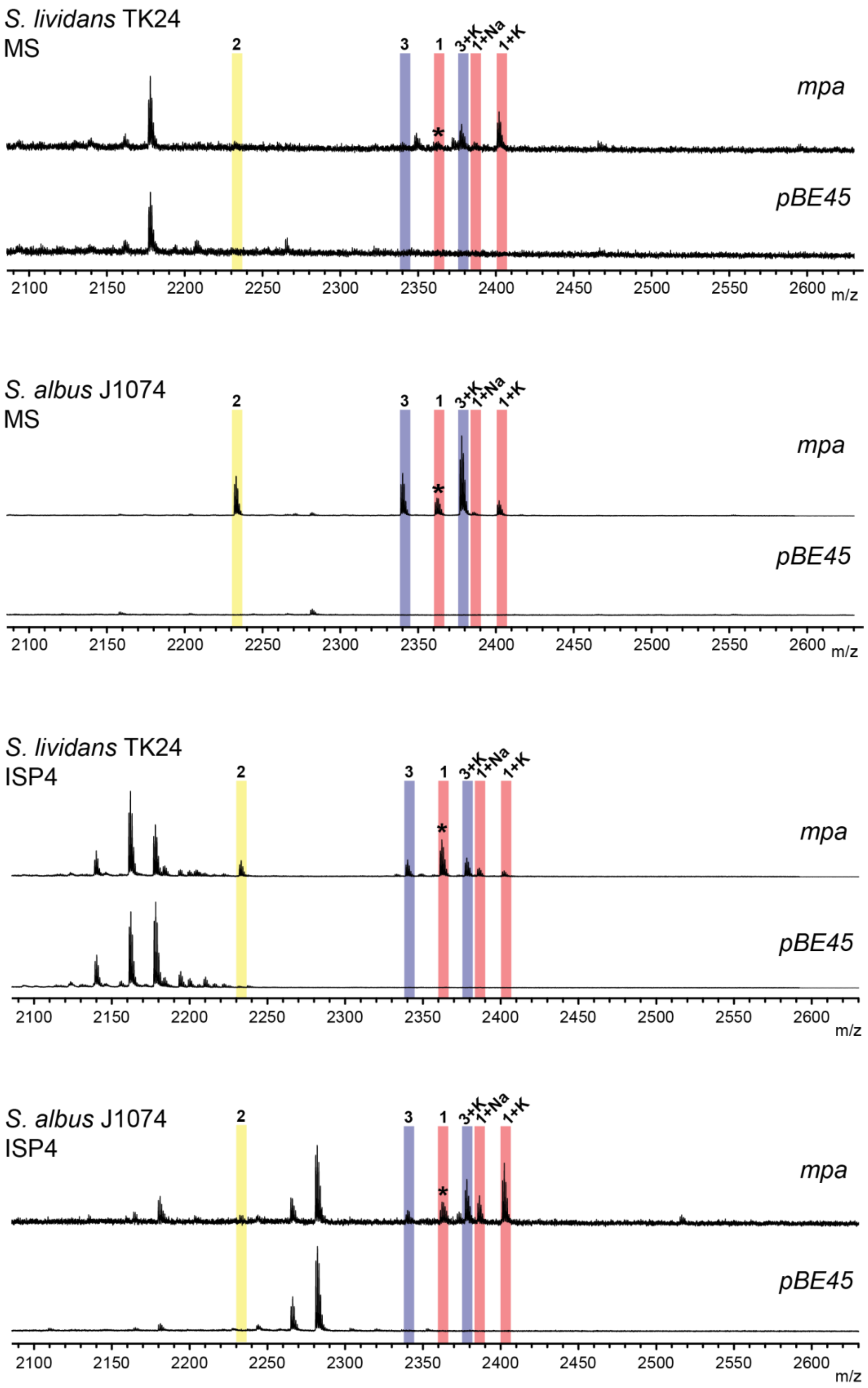

**Supplementary Figure 9: Media and host screen for *mpa* BGC expression**. MALDI-TOF mass spectra of methanolic extracts of *S. lividans* TK24 and *S. albus* J1074 colonies that were transformed with *mpa* or the pBE45 empty vector and cultivated on MS and ISP4 agar media. Possible overlapped peaks of $[\mathbf{1}+H]^+$ and $[\mathbf{3}+Na]^+$ are indicated with an asterisk.

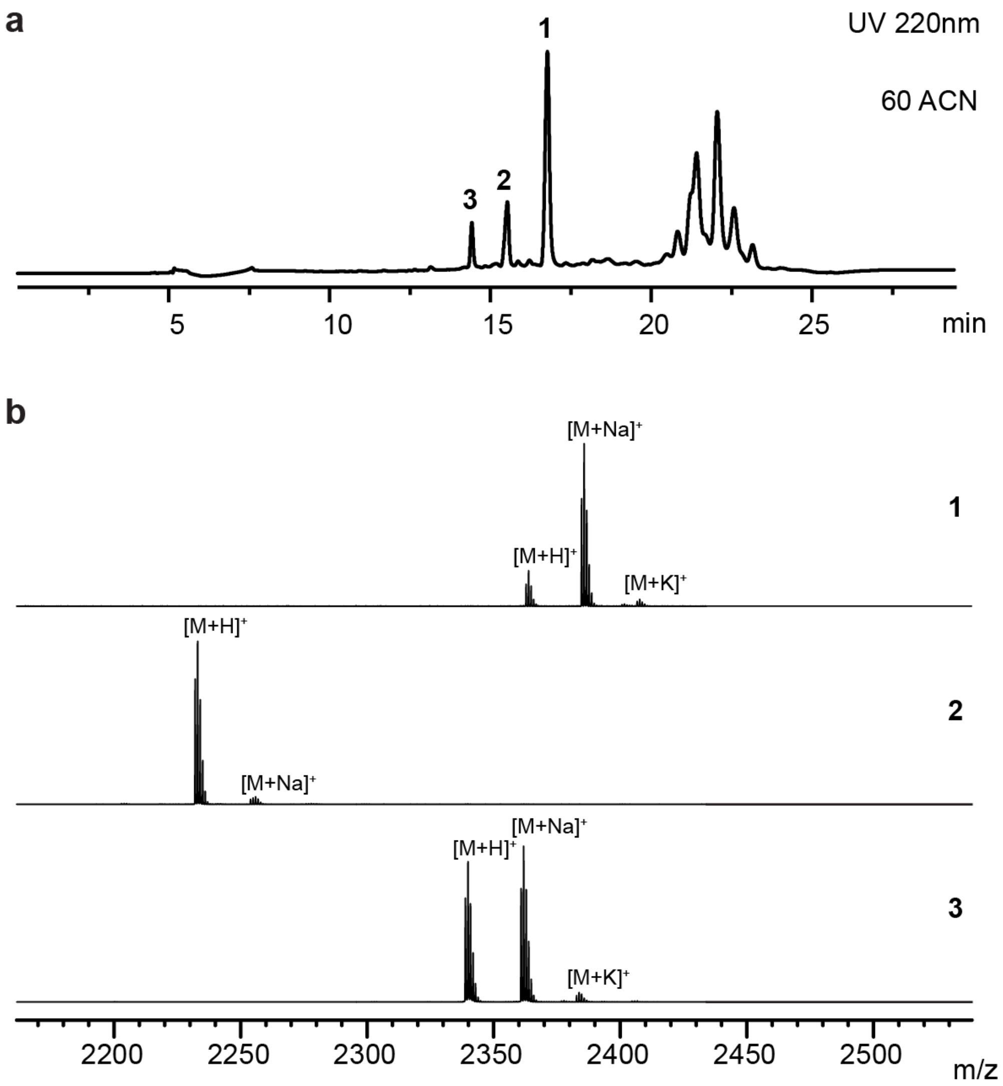

**Supplementary Figure 10: Purification of daptides 1-3.** (**a**) Reverse phase (C18) HPLC chromatograms with UV absorbance (220 nm) monitoring of solid phase extraction eluents using 60% acetonitrile. (**b**) MALDI-TOF-MS analysis of the collected fractions.

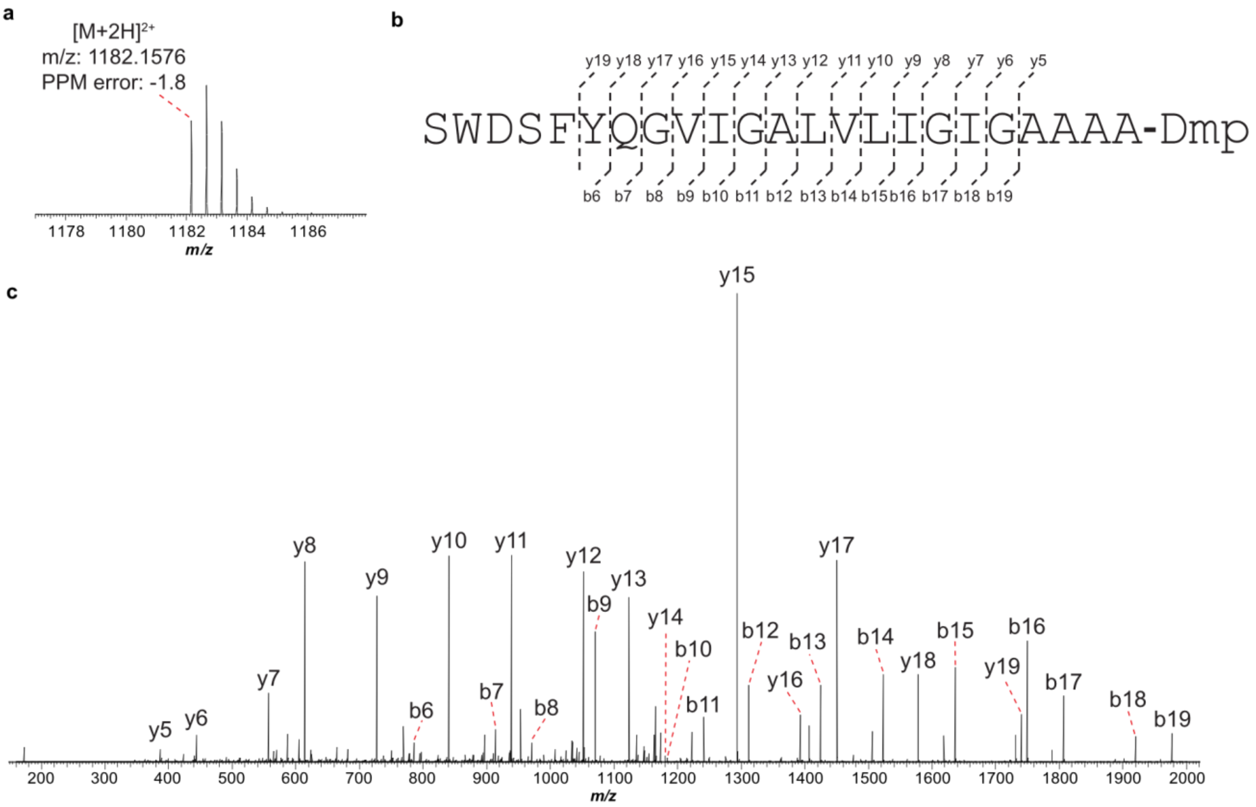

d

| Ion | Calc. Mass | Obs. Mass | PPM Error |
|---|---|---|---|
| **[M+2H]$^{2+}$** | 1182.1597 | 1182.1576 | -1.8 |
| **b6** | 786.3093 | 786.3082 | -1.4 |
| **b7** | 914.3679 | 914.3664 | -1.6 |
| **b8** | 971.3894 | 971.3877 | -1.8 |
| **b9** | 1070.4578 | 1070.4558 | -1.9 |
| **b10** | 1183.5419 | 1183.5393 | -2.2 |
| **b11** | 1240.5633 | 1240.5610 | -1.9 |
| **b12** | 1311.6004 | 1311.5981 | -1.8 |
| **b13** | 1424.6845 | 1424.6827 | -1.3 |
| **b14** | 1523.7529 | 1523.7514 | -1.0 |
| **b15** | 1636.8370 | 1636.8348 | -1.3 |
| **b16** | 1749.9210 | 1749.9180 | -1.7 |
| **b17** | 1806.9425 | 1806.9382 | -2.4 |
| **b18** | 1920.0266 | 1920.0216 | -2.6 |
| **b19** | 1977.0480 | 1977.0431 | -2.5 |

| Ion | Calc. Mass | Obs. Mass | PPM Error |
|---|---|---|---|
| **y5** | 387.2714 | 387.2710 | -1.0 |
| **y6** | 444.2929 | 444.2922 | -1.6 |
| **y7** | 557.3770 | 557.3761 | -1.6 |
| **y8** | 614.3984 | 614.3975 | -1.5 |
| **y9** | 727.4825 | 727.4813 | -1.6 |
| **y10** | 840.5665 | 840.5650 | -1.8 |
| **y11** | 939.6350 | 939.6333 | -1.8 |
| **y12** | 1052.7190 | 1052.7171 | -1.8 |
| **y13** | 1123.7561 | 1123.7537 | -2.1 |
| **y14** | 1180.7776 | 1180.7751 | -2.1 |
| **y15** | 1293.8617 | 1293.8594 | -1.8 |
| **y16** | 1392.9301 | 1392.9279 | -1.6 |
| **y17** | 1449.9515 | 1449.9497 | -1.2 |
| **y18** | 1578.0101 | 1578.0079 | -1.4 |
| **y19** | 1741.0735 | 1741.0704 | -1.8 |

**Supplementary Figure 11: HR-MS/MS analysis of 1**. (**a**) Broad band high-resolution mass spectrum of compound **1**. (**b**) Summary of collision-induced dissociation (CID) daughter ions for **1**. Observed b- and y-ions are annotated. Ions are in the +1 charge state except where indicated. Dmp: $N_2$,$N_2$-dimethyl-1,2-propanediamine. (**c**) CID spectrum of **1**. (**d**) Table of daughter ion assignments for **1**.

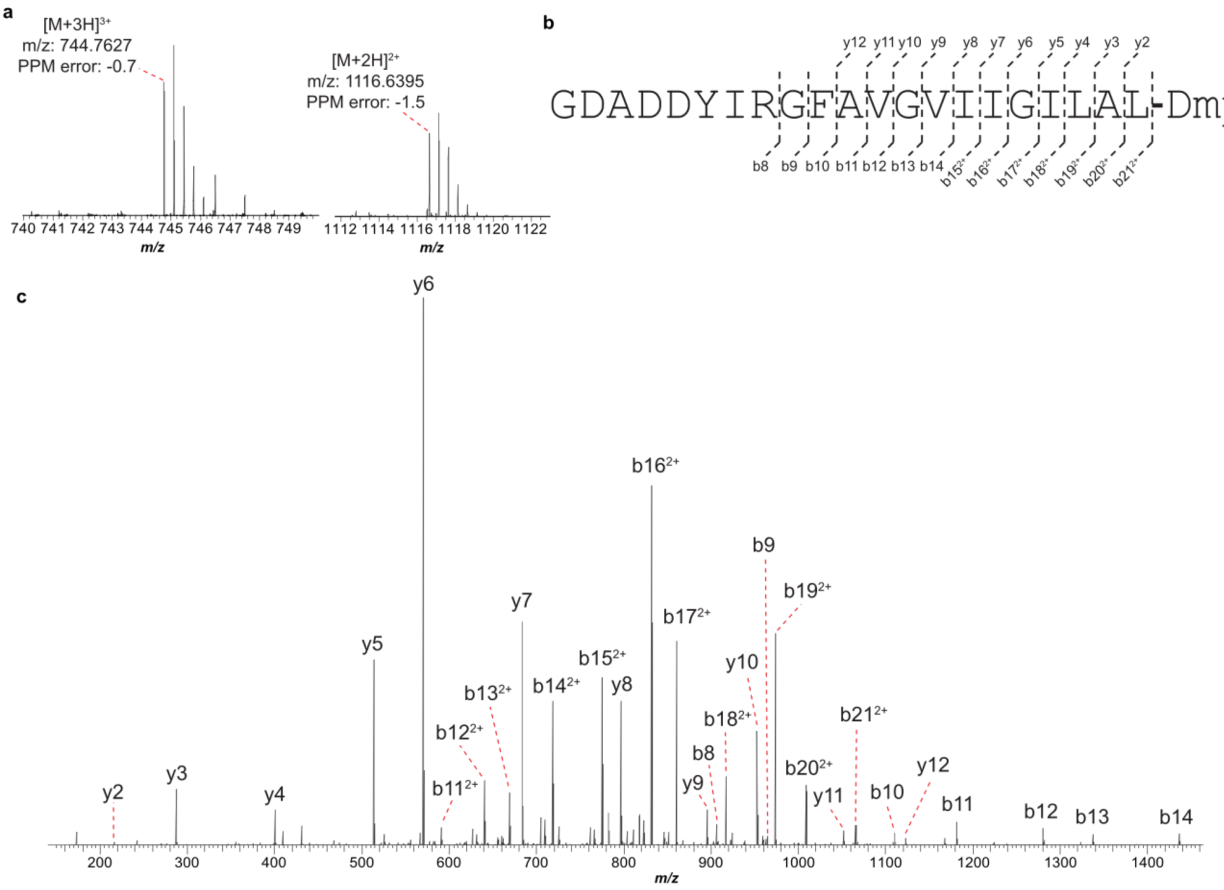

d

| Ion | Calc. Mass | Obs. Mass | PPM Error |
|---|---|---|---|
| **[M+3H]³⁺** | 744.7632 | 744.7627 | -0.7 |
| **[M+2H]²⁺** | 1116.6412 | 1116.6395 | -1.5 |
| **b8** | 906.3952 | 906.3939 | -1.4 |
| **b9** | 963.4167 | 963.4150 | -1.8 |
| **b10** | 1110.4851 | 1110.4829 | -2.0 |
| **b11** | 1181.5222 | 1181.5196 | -2.2 |
| **b11²⁺** | 591.2647 | 591.2639 | -1.4 |
| **b12** | 1280.5906 | 1280.5882 | -1.9 |
| **b12²⁺** | 640.7989 | 640.7981 | -1.3 |
| **b13** | 1337.6121 | 1337.6095 | -1.9 |
| **b13²⁺** | 669.3097 | 669.3088 | -1.3 |
| **b14** | 1436.6805 | 1436.6791 | -1.0 |
| **b14²⁺** | 718.8439 | 718.8428 | -1.5 |
| **b15²⁺** | 775.3859 | 775.3849 | -1.3 |
| **b16²⁺** | 831.9279 | 831.9269 | -1.2 |
| **b17²⁺** | 860.4387 | 860.4372 | -1.7 |

| Ion | Calc. Mass | Obs. Mass | PPM Error |
|---|---|---|---|
| **b18²⁺** | 916.9807 | 916.9793 | -1.5 |
| **b19²⁺** | 973.5227 | 973.5211 | -1.7 |
| **b20²⁺** | 1009.0413 | 1009.0397 | -1.6 |
| **b21²⁺** | 1065.5833 | 1065.5812 | -2.0 |
| **y2** | 216.2067 | 216.2067 | -1.4 |
| **y3** | 287.2438 | 287.2438 | -1.4 |
| **y4** | 400.3277 | 400.3277 | -1.2 |
| **y5** | 513.4116 | 513.4116 | -1.4 |
| **y6** | 570.4330 | 570.4330 | -1.2 |
| **y7** | 683.5168 | 683.5168 | -1.5 |
| **y8** | 796.6007 | 796.6007 | -1.5 |
| **y9** | 895.6689 | 895.6689 | -1.6 |
| **y10** | 952.6902 | 952.6902 | -1.6 |
| **y11** | 1051.7580 | 1051.7580 | -2.1 |
| **y12** | 1122.7951 | 1122.7951 | -2.0 |

**Supplementary Figure 12: HR-MS/MS analysis of 2**. (**a**) Broad band high-resolution mass spectrum of compound **2**. The $[M+3H]^{3+}$ ion was selected for fragmentation. (**b**) Summary of collision-induced dissociation (CID) daughter ions for **2**. Observed b- and y-ions are annotated. Ions are in the +1 charge state except where indicated. Dmp: $N_2$,$N_2$-dimethyl-1,2-propanediamine. (**c**) CID spectrum of **2**. (**d**) Table of daughter ion assignments for **2**.

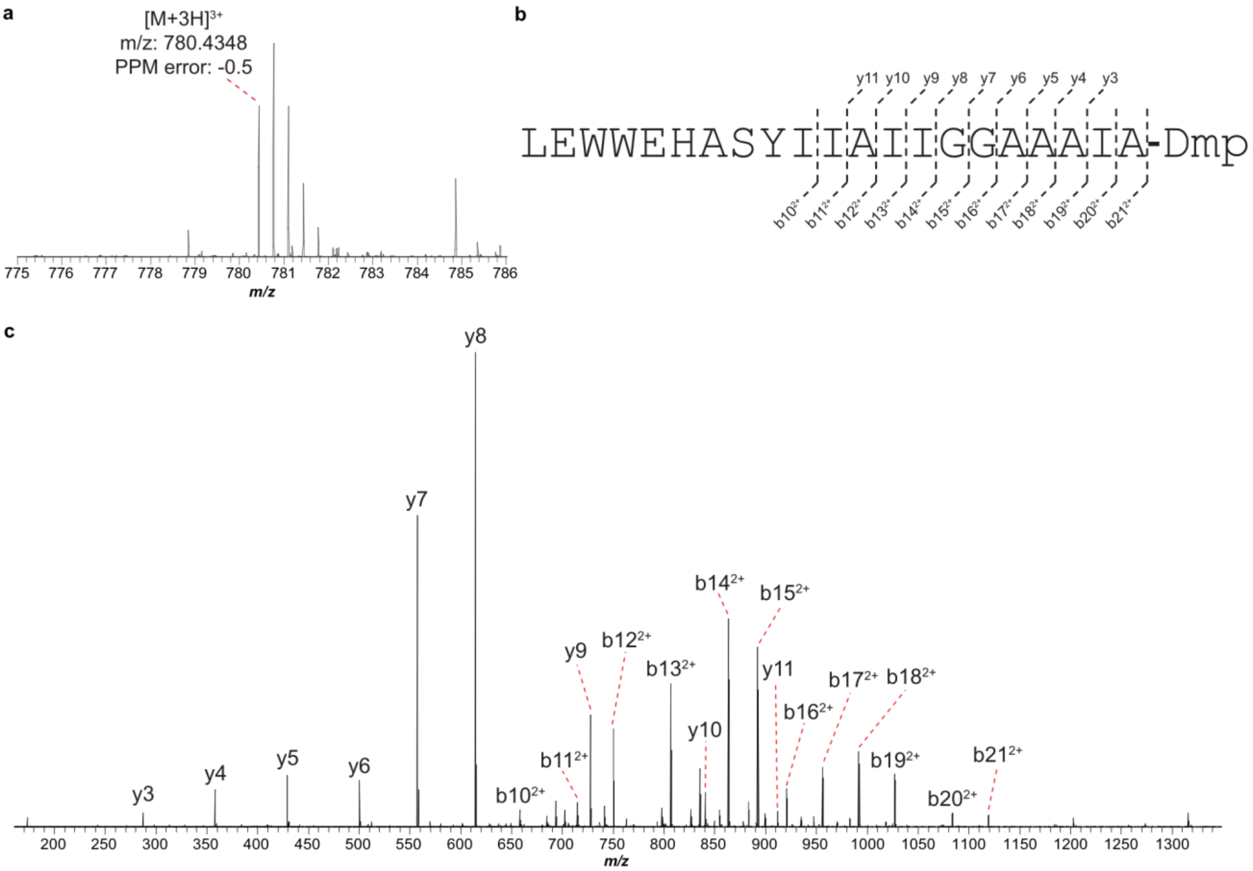

d

| Ion | Calc. Mass | Obs. Mass | PPM Error |
|---|---|---|---|
| **$[M+3H]^{3+}$** | 780.4352 | 780.4348 | -0.5 |
| **$b10^{2+}$** | 658.3089 | 658.3079 | -1.6 |
| **$b11^{2+}$** | 714.8510 | 714.8497 | -1.8 |
| **$b12^{2+}$** | 750.3695 | 750.3681 | -1.9 |
| **$b13^{2+}$** | 806.9115 | 806.9101 | -1.8 |
| **$b14^{2+}$** | 863.4536 | 863.4519 | -2.0 |
| **$b15^{2+}$** | 891.9643 | 891.9625 | -2.1 |
| **$b16^{2+}$** | 920.4750 | 920.4732 | -2.0 |
| **$b17^{2+}$** | 955.9936 | 955.9916 | -2.1 |
| **$b18^{2+}$** | 991.5122 | 991.5101 | -2.1 |
| **$b19^{2+}$** | 1027.0307 | 1027.0284 | -2.3 |

| Ion | Calc. Mass | Obs. Mass | PPM Error |
|---|---|---|---|
| **$b20^{2+}$** | 1083.5727 | 1083.5702 | -2.3 |
| **$b21^{2+}$** | 1119.0913 | 1119.0885 | -2.5 |
| **y3** | 287.2442 | 287.2436 | -2.1 |
| **y4** | 358.2813 | 358.2807 | -1.7 |
| **y5** | 429.3184 | 429.3176 | -1.9 |
| **y6** | 500.3555 | 500.3546 | -1.8 |
| **y7** | 557.3770 | 557.3759 | -2.0 |
| **y8** | 614.3984 | 614.3972 | -2.0 |
| **y9** | 727.4825 | 727.4812 | -1.8 |
| **y10** | 840.5665 | 840.5648 | -2.0 |
| **y11** | 911.6037 | 911.6019 | -2.0 |

**Supplementary Figure 13: HR-MS/MS analysis of 3**. (**a**) Broad band high-resolution mass spectrum of compound **3**. (**b**) Summary of collision-induced dissociation (CID) daughter ions for **3**. Observed b- and y-ions are annotated. Ions are in the +1 charge state except where indicated. Dmp: $N_2$,$N_2$-dimethyl-1,2-propanediamine. (**c**) CID spectrum of **3**. (**d**) Table of daughter ion assignments for **3**.

**a**

$^{1}H$ NMR, 1,1,1,3,3,3-Hexafluoroisopropanol-$d_2$ (HFIP-$d_2$), 750 MHz. The solvent peak appears at 4.41 ppm.

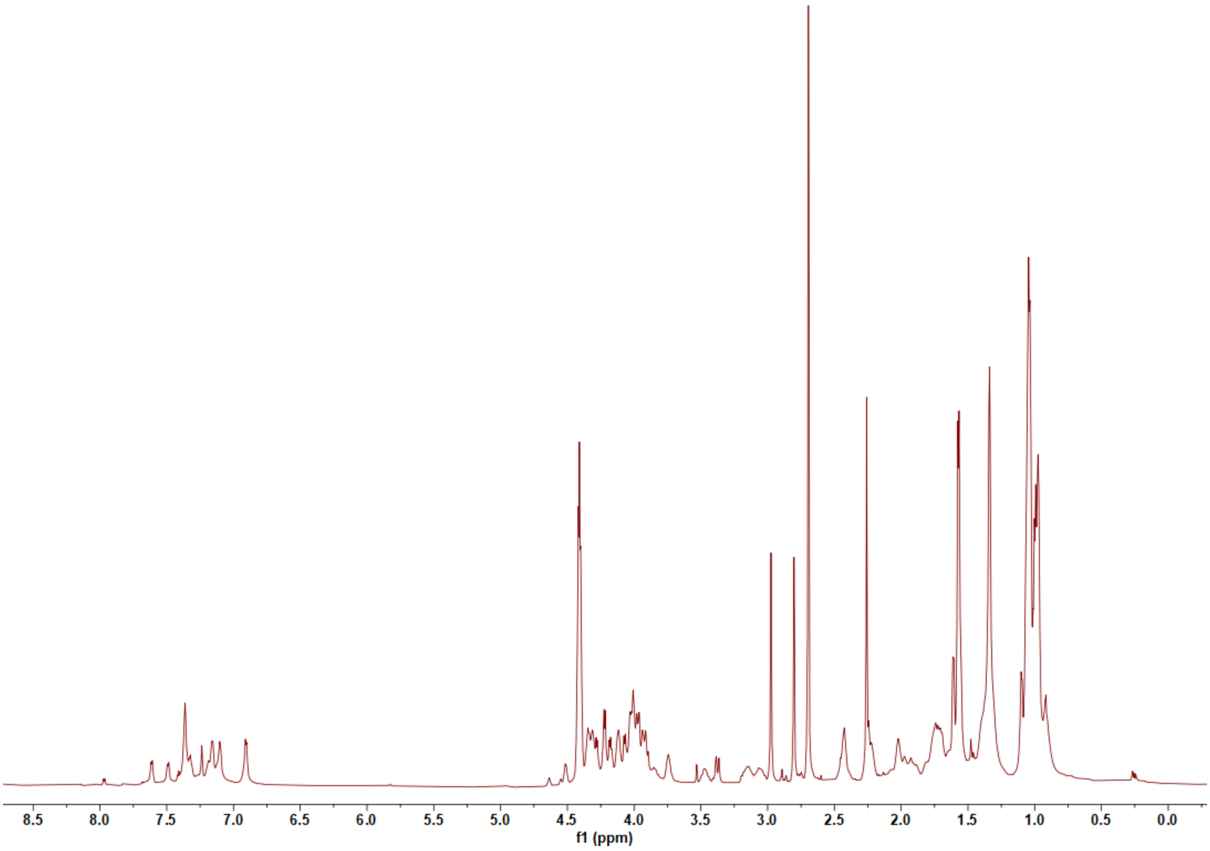

**b**

$^{13}$C NMR, 1,1,1,3,3,3-Hexafluoroisopropanol-$d_2$ (HFIP-$d_2$), 750 MHz. The solvent peaks appear at 68.1 and 120.7 ppm.

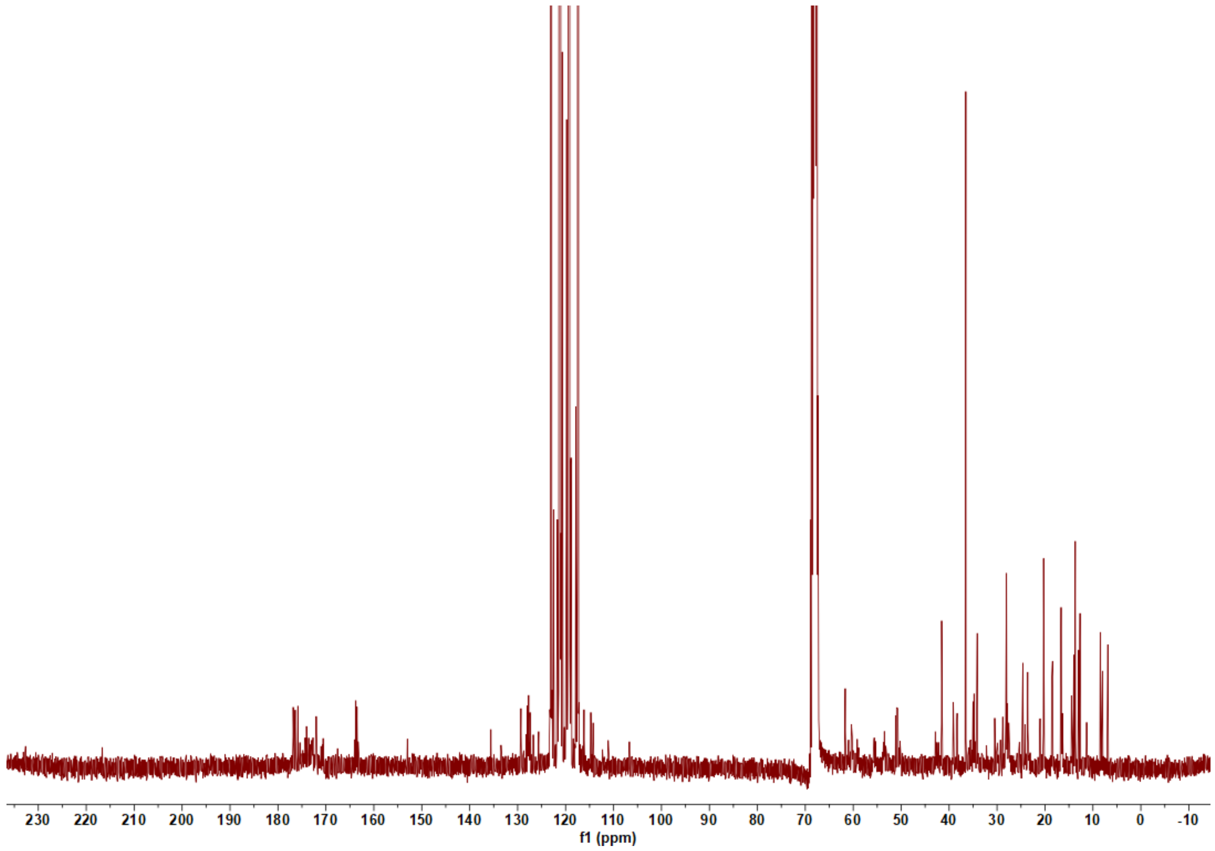

**c**

$^{1}$H-$^{1}$H COSY NMR, 1,1,1,3,3,3-Hexafluoroisopropanol-$d_2$ (HFIP-$d_2$), 750 MHz.

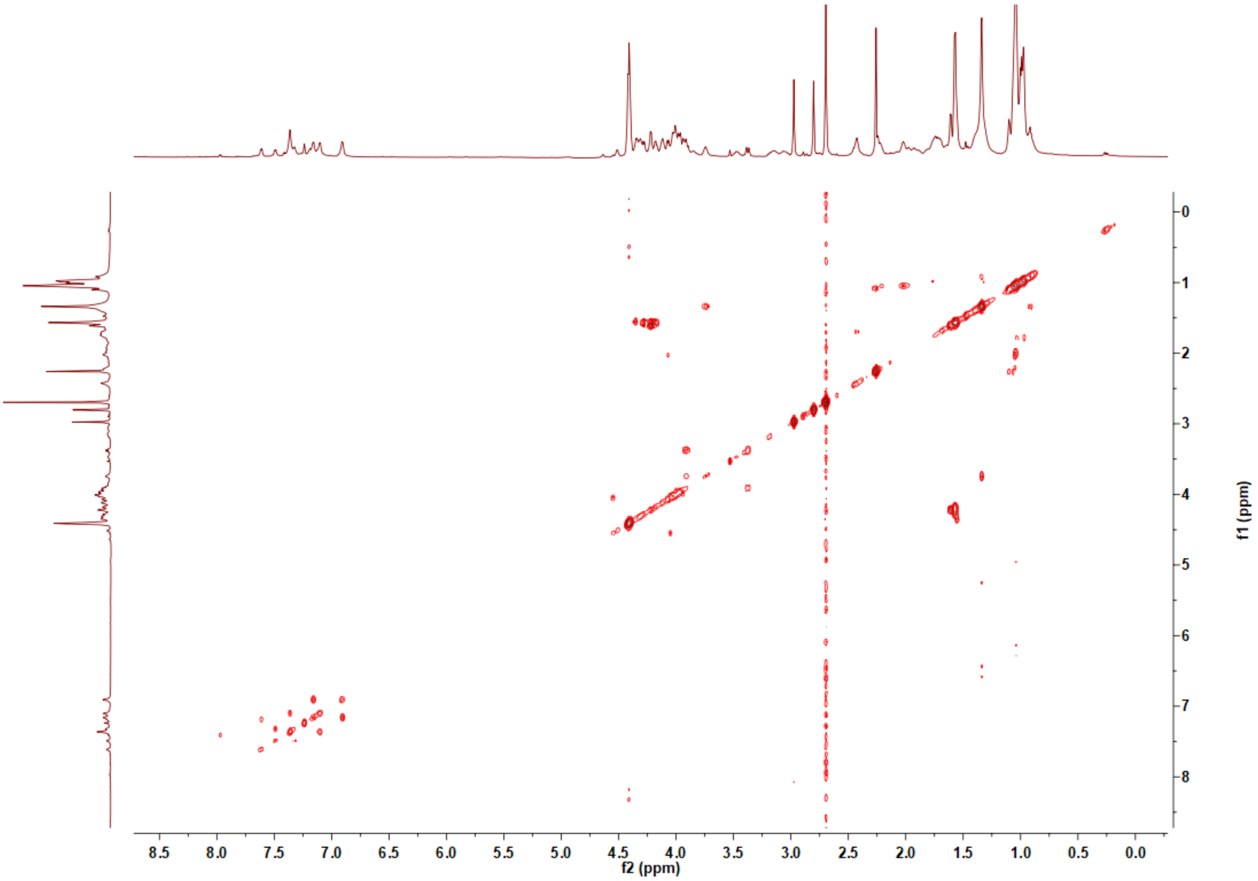

**d**

$^{1}H$-$^{13}C$ HSQC NMR, 1,1,1,3,3,3-Hexafluoroisopropanol-$d_2$ (HFIP-$d_2$), 750 MHz. Multiplicity-edited HSQC was performed. Signals of $CH_2$ and $CH/CH_3$ are shown in blue and red, respectively.

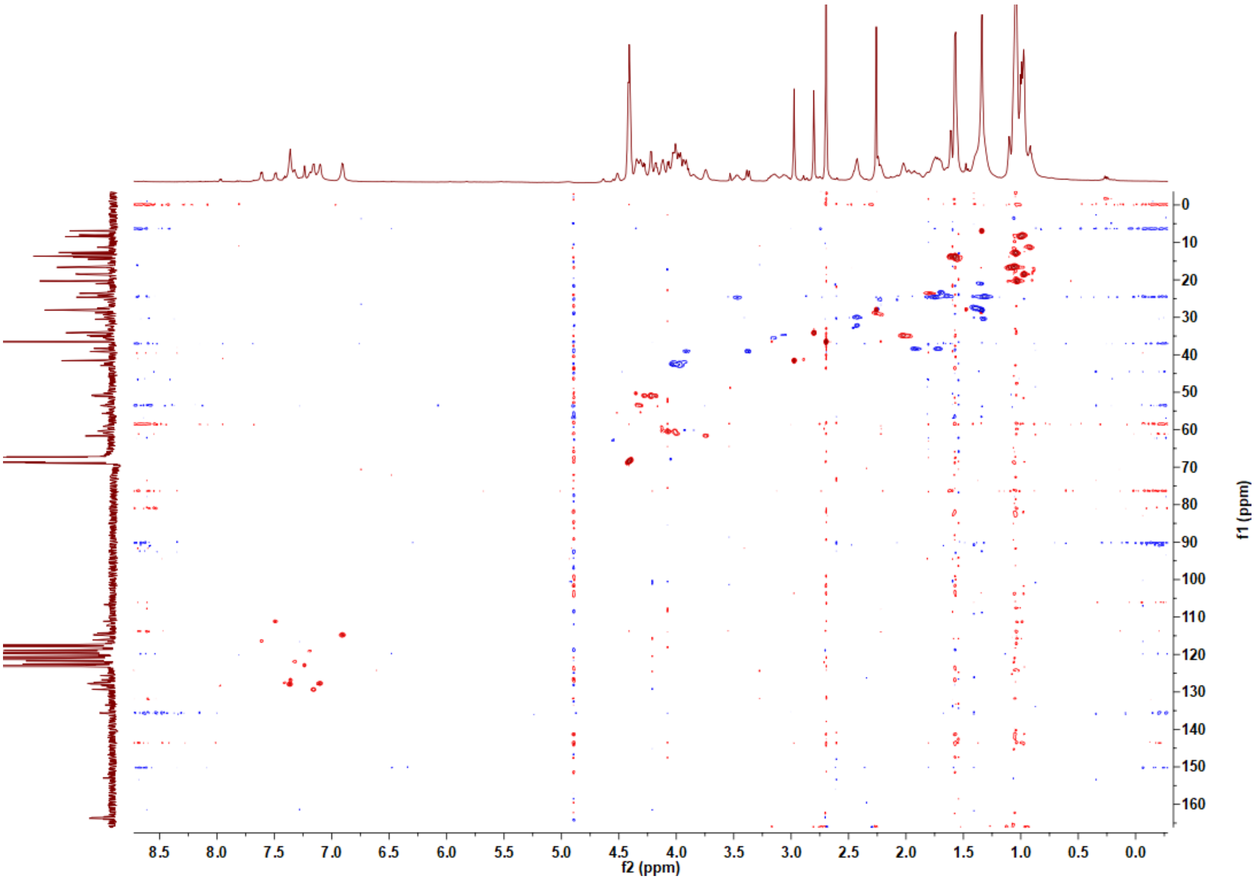

**e**

$^{1}H$-$^{13}C$ HMBC NMR, 1,1,1,3,3,3-Hexafluoroisopropanol-$d_2$ (HFIP-$d_2$), 750 MHz.

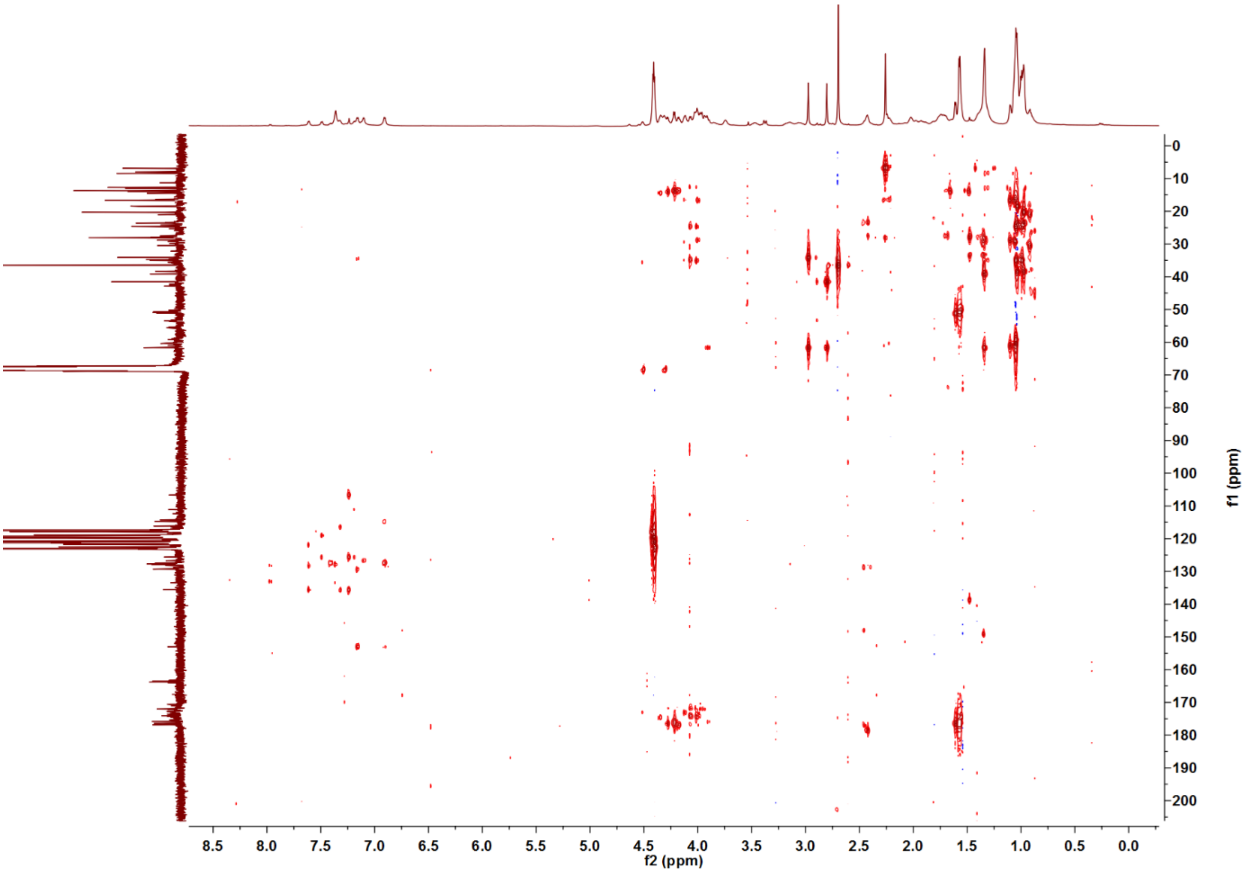

**f**

COSY focusing on the Dmp region. Dmp: $N_2$,$N_2$-dimethyl-1,2-propanediamine.

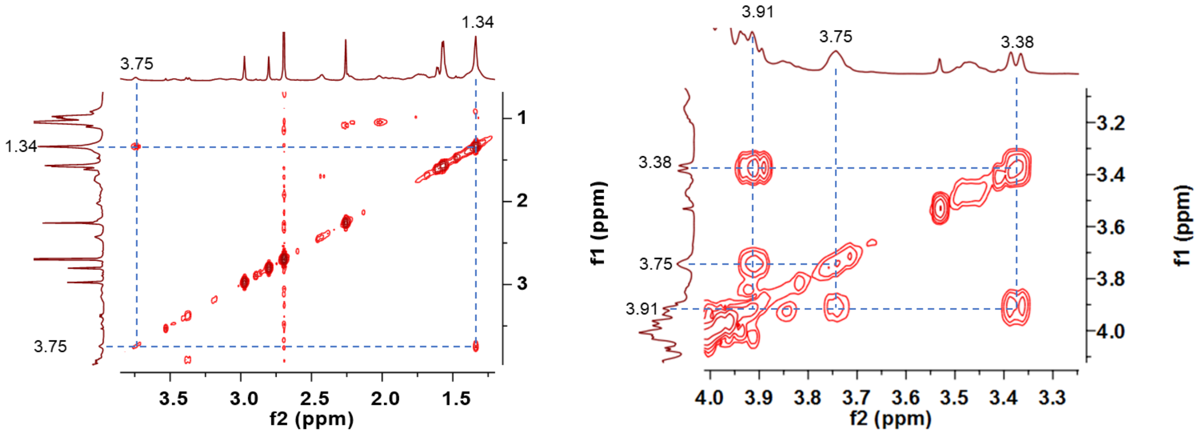

**g**

HSQC focusing on the Dmp region. Multiplicity-edited HSQC was performed. Signals of $CH_2$ and $CH/CH_3$ are shown in blue and red, respectively. Dmp: $N_2$,$N_2$-dimethyl-1,2-propanediamine.

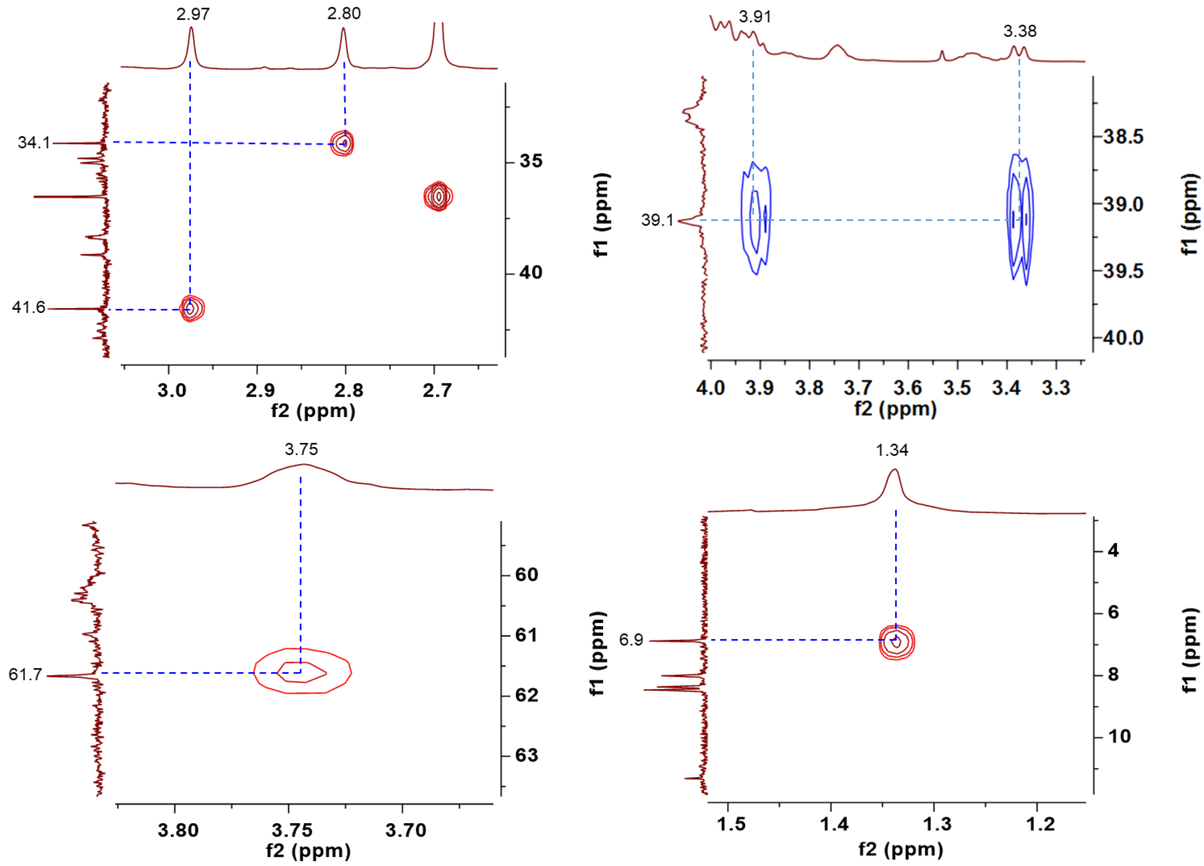

**h**

HMBC focusing on the Dmp region. Dmp: $N_2$,$N_2$-dimethyl-1,2-propanediamine.

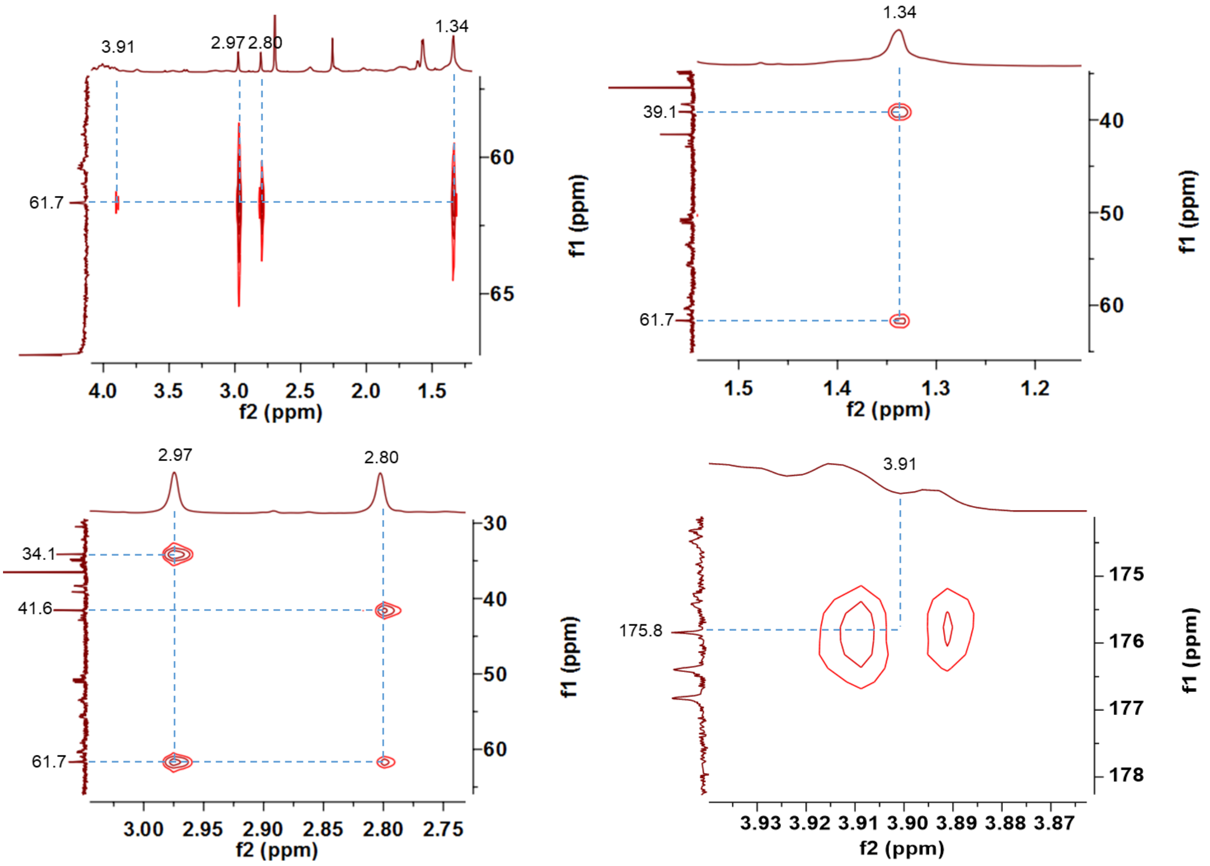

**i**

HSQC focusing on the Ala-23 region. Multiplicity-edited HSQC was performed. Signals of $CH_2$ and $CH/CH_3$ are shown in blue and red, respectively.

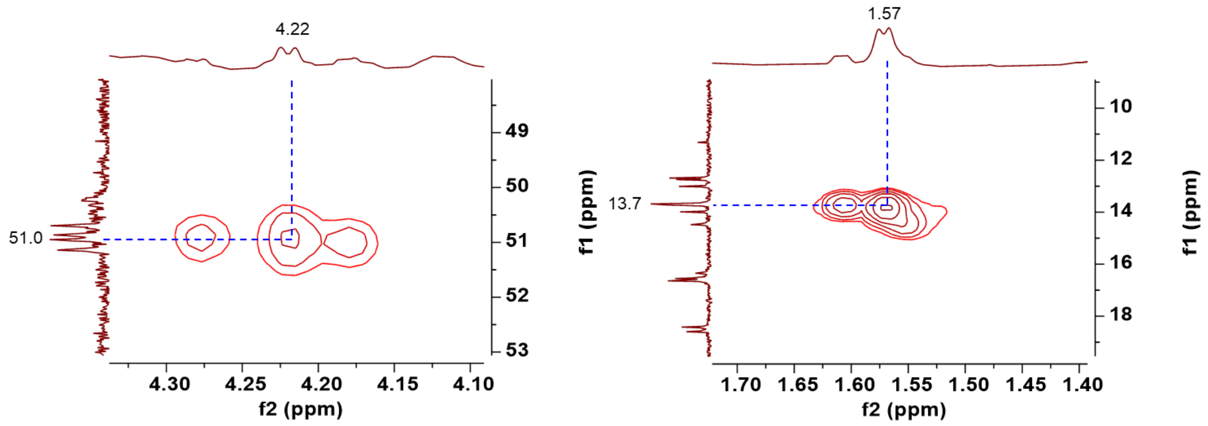

**j**

HMBC focusing on the Ala-23 region.

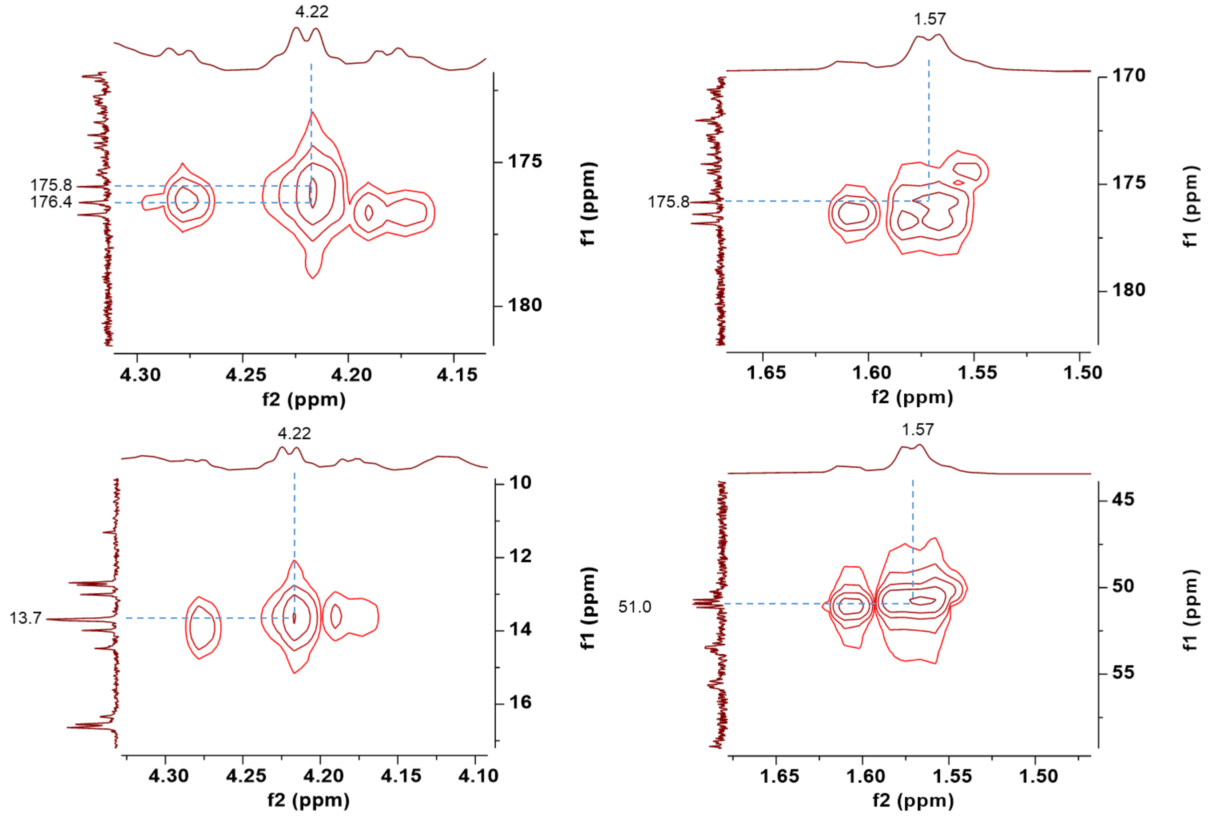

**k**

Summary of HMBC correlations in the Dmp and Ala-23 region. Dmp: $N_2$,$N_2$-dimethyl-1,2-propanediamine.

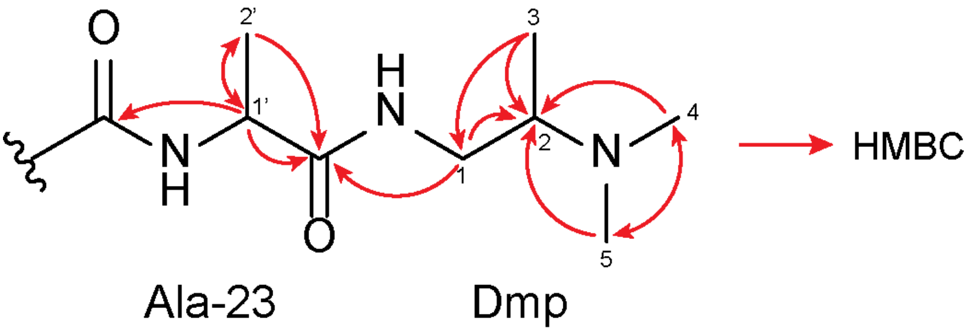

**I**

COSY focusing on the Trp-2 region

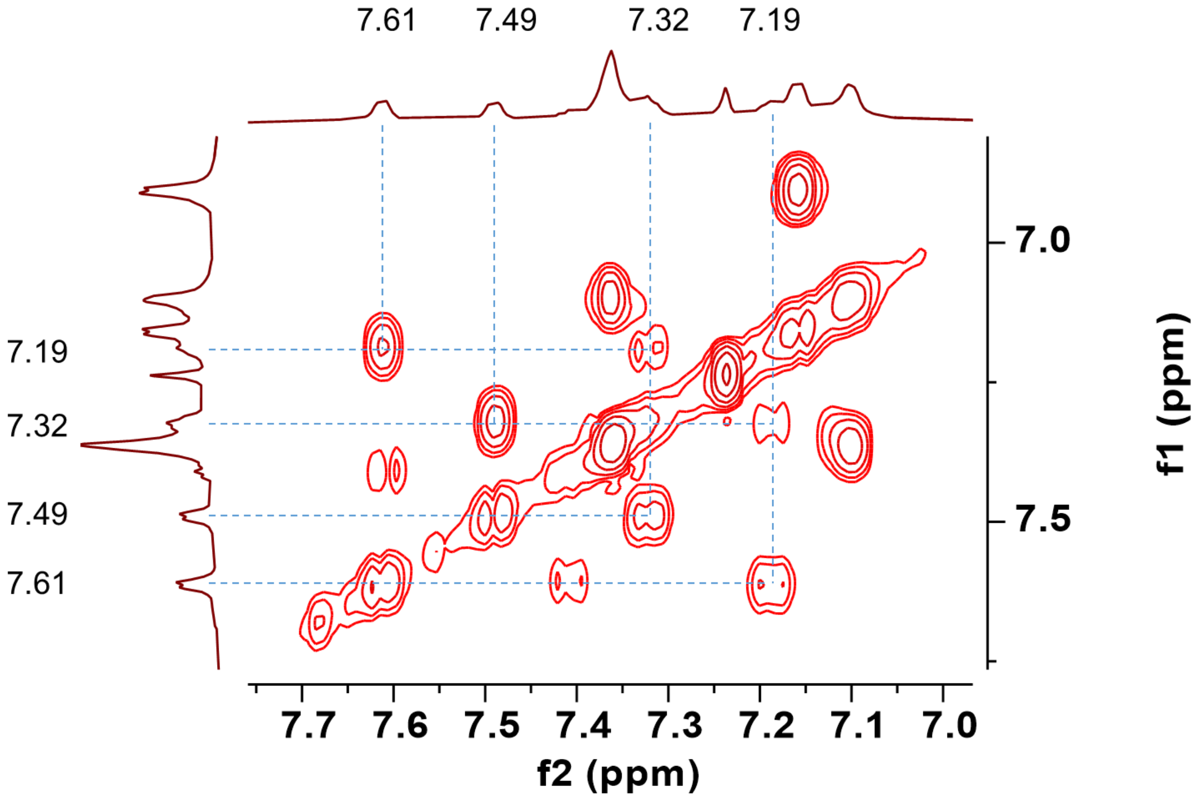

**m**

HSQC focusing on the Trp-2 region. Multiplicity-edited HSQC was performed. Signals of CH2 and CH/CH3 are shown in blue and red, respectively.

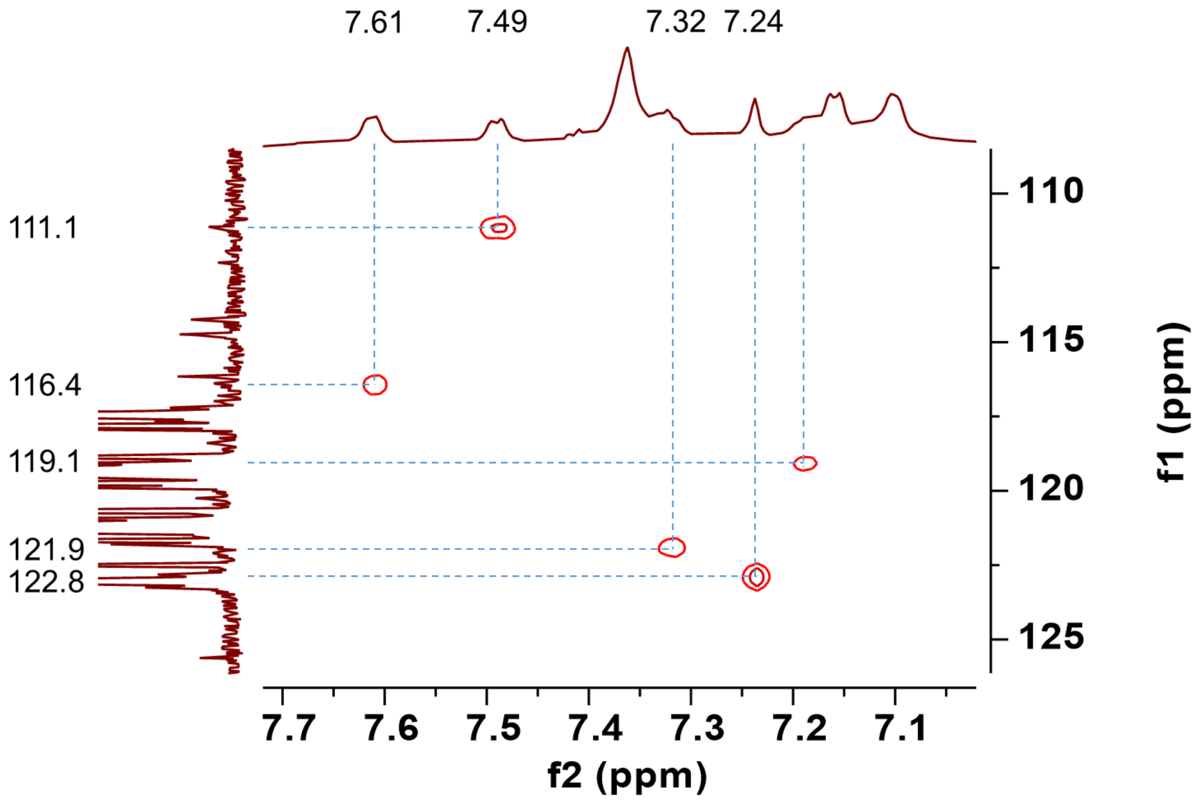

**n**

HMBC focusing on the Trp-2 region.

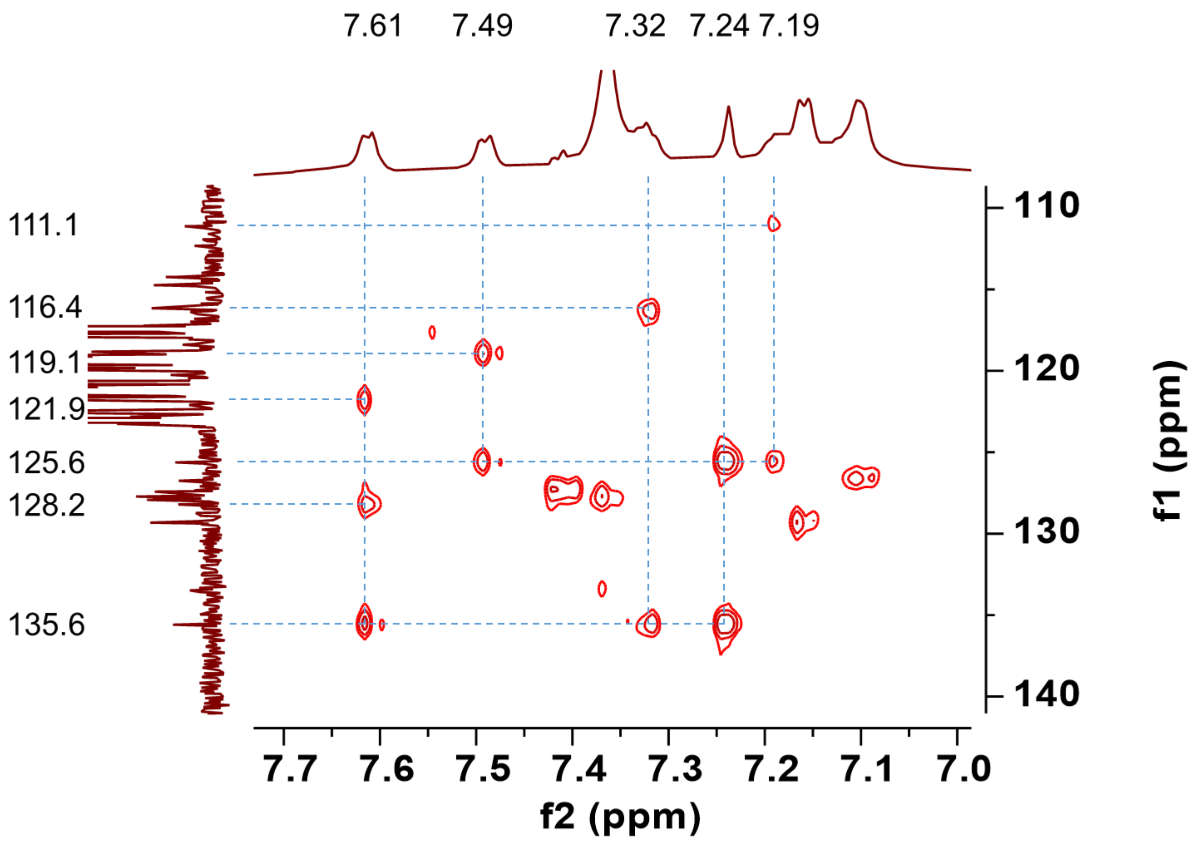

**o**

Summary of COSY and HMBC correlations in the Trp-2 region.

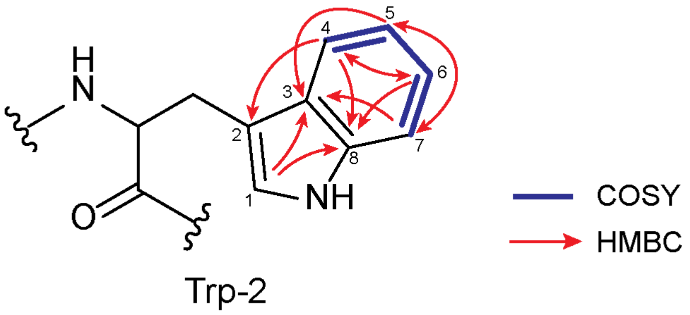

| No. | 1 (daptide) | |
|---|---|---|
| | $\delta_C$ type | $\delta_H$ multi |
| 1 | 122.8 CH | 7.24 s |
| 2 | 128.2 C | |
| 3 | 125.6 C | |
| 4 | 116.4 CH | 7.61 d (7.1) |
| 5 | 119.1 CH | 7.19 overlapped |
| 6 | 121.9 CH | 7.32 overlapped |
| 7 | 111.1 CH | 7.49 d (7.9) |
| 8 | 135.6 C | |

**p**

COSY focusing on the Tyr-6 region

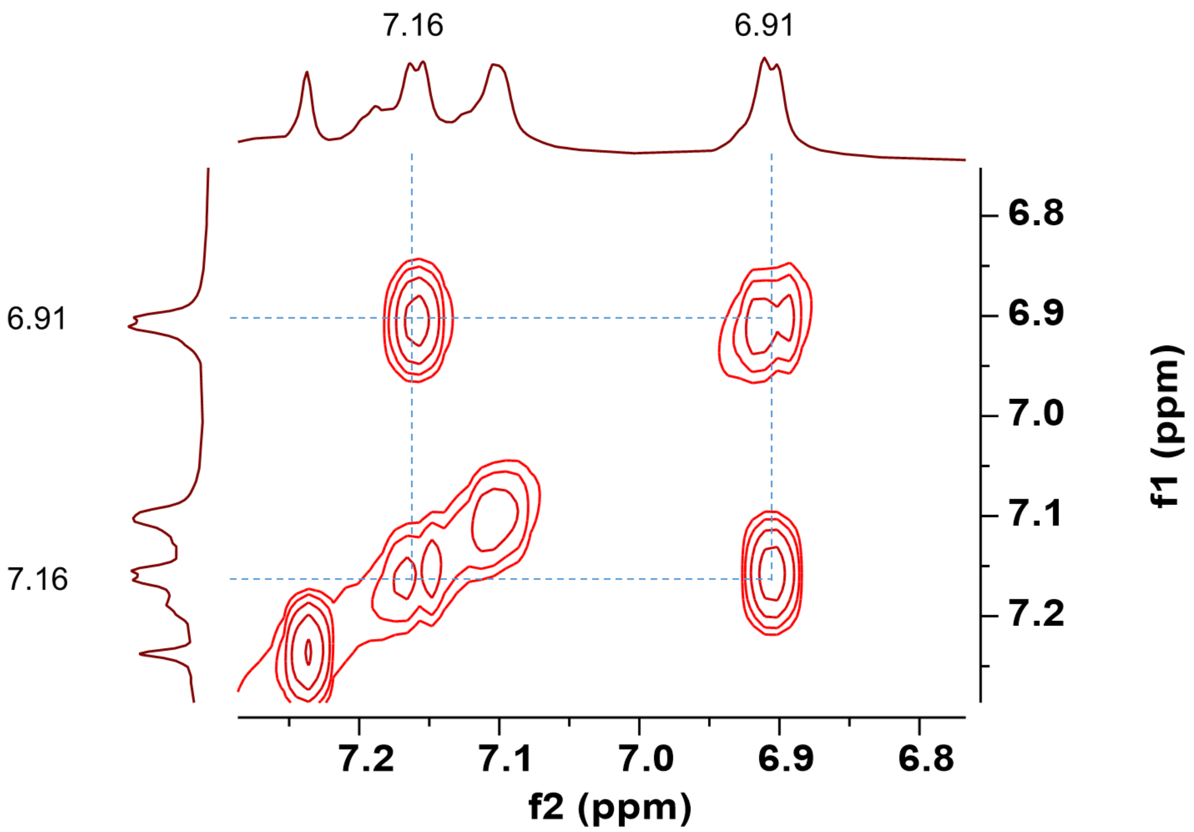

**q**

HSQC focusing on the Tyr-6 region. Multiplicity-edited HSQC was performed. Signals of CH2 and CH/CH3 are shown in blue and red, respectively.

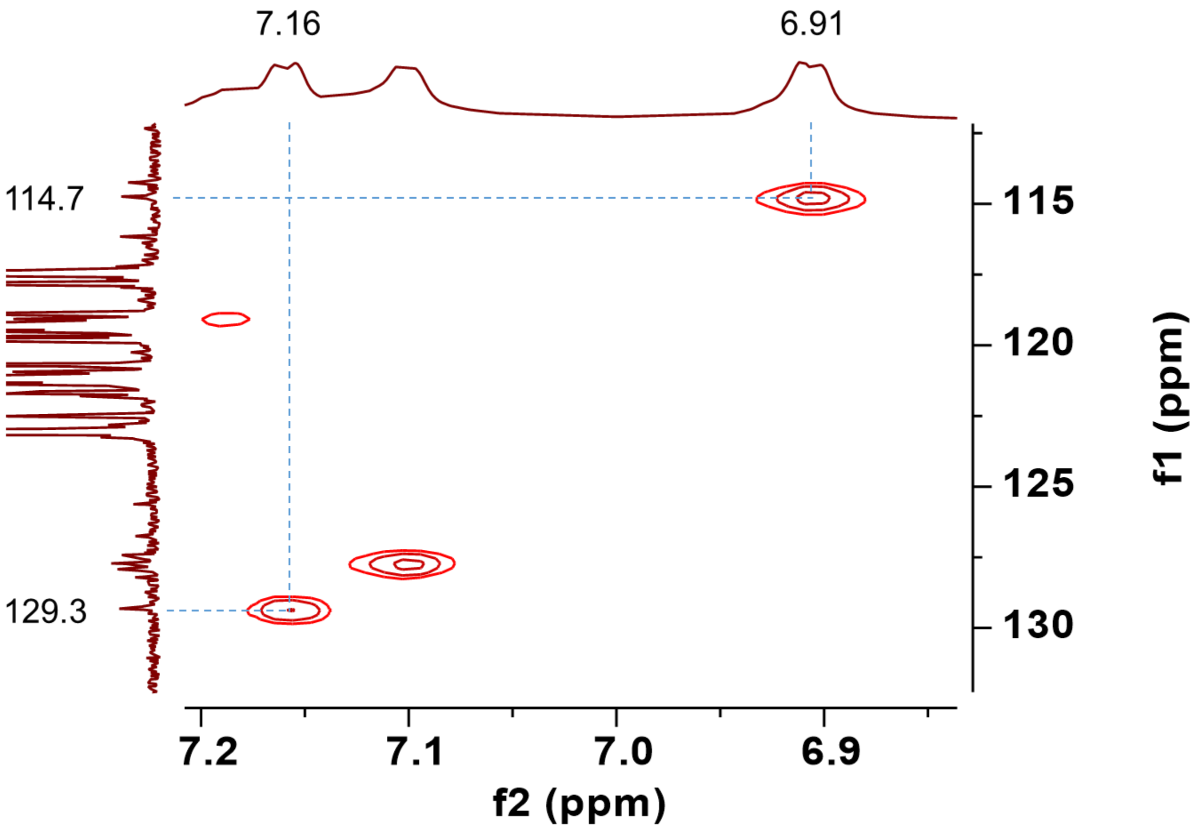

**r**

HMBC focusing on the Tyr-6 region.

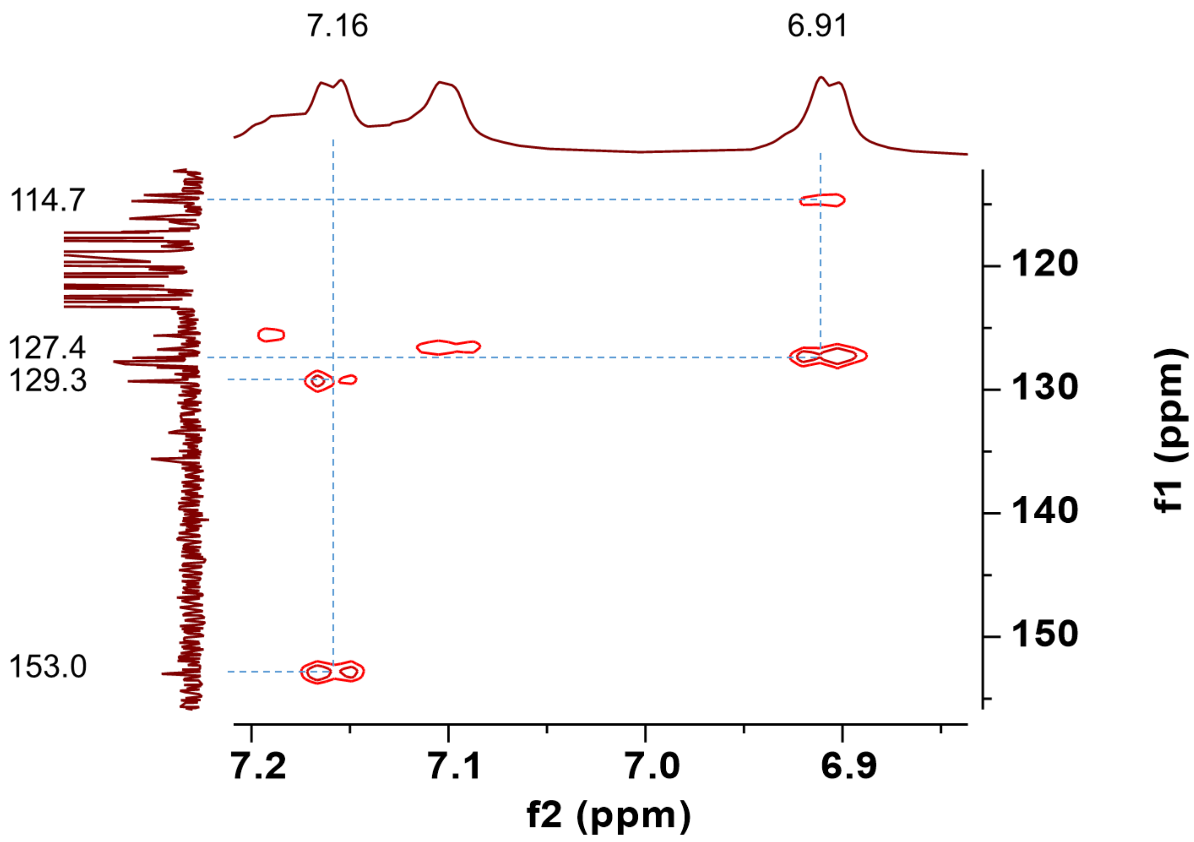

**s**

Summary of COSY and HMBC correlations in the Tyr-6 region.

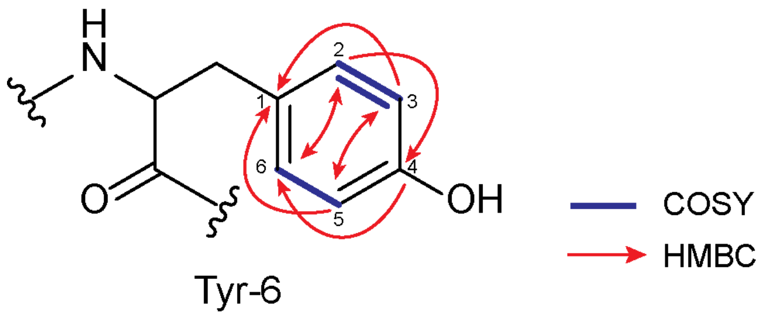

| No. | 1 (daptide) | |
|---|---|---|
| | δC type | δH multi |
| 1 | 127.4 C | |
| 2 | 129.3 CH | 7.16 overlapped |
| 3 | 114.7 CH | 6.91 d (7.1) |
| 4 | 153.0 C | |
| 5 | 114.7 CH | 6.91 d (7.1) |
| 6 | 129.3 CH | 7.16 overlapped |

**Supplementary Figure 14: NMR spectra of compound 1 in 1,1,3,3,3-hexafluoroisopropanol-d2 (HFIP-*d*2).** (**a**) $^{1}$H NMR. The solvent peak appears at 4.41 ppm. (**b**) $^{13}$C NMR. The solvent peaks appear at 68.1 and 120.7 ppm. (**c**) $^{1}$H-$^{1}$H COSY NMR. (**d**) $^{1}$H-$^{13}$C HSQC NMR. Multiplicity-edited HSQC was performed. Signals of $CH_2$ and $CH/CH_3$ are shown in blue and red, respectively. (**e**) $^{1}$H-$^{13}$C HMBC NMR. (**f**) COSY focusing on the Dmp region. Dmp: $N_2$,$N_2$-dimethyl-1,2-propanediamine. (**g**) HSQC focusing on the Dmp region. Multiplicity-edited HSQC was performed. Signals of $CH_2$ and $CH/CH_3$ are shown in blue and red, respectively. (**h**) HMBC focusing on the Dmp region. (**i**) HSQC focusing on the Ala-20 region. Multiplicity-edited HSQC was performed. Signals of $CH_2$ and $CH/CH_3$ are shown in blue and red, respectively. (**j**) HMBC focusing on the Ala-20 region. (**k**) Summary of HMBC correlations in the Dmp and Ala-20 region. Dmp: $N_2$,$N_2$-dimethyl-1,2-propanediamine. (**l**) COSY focusing on the Trp-2 region. (**m**) HSQC focusing on the Trp-2 region. Multiplicity-edited HSQC was performed. Signals of CH2 and CH/CH3 are shown in blue and red, respectively. (**n**) HMBC focusing on the Trp-2 region. (**o**) Summary of COSY and HMBC correlations in the Trp-2 region. (**p**) COSY focusing on the Tyr-6 region (**q**) HSQC focusing on the Tyr-6 region. Multiplicity-edited HSQC was performed. Signals of CH2 and CH/CH3 are shown in blue and red, respectively. (**r**) HMBC focusing on the Tyr-6 region. (**s**) Summary of COSY and HMBC correlations in the Tyr-6 region.

**a**

(*R*)-Dmp $^1$H NMR, DMSO-$d_6$, 500 MHz.

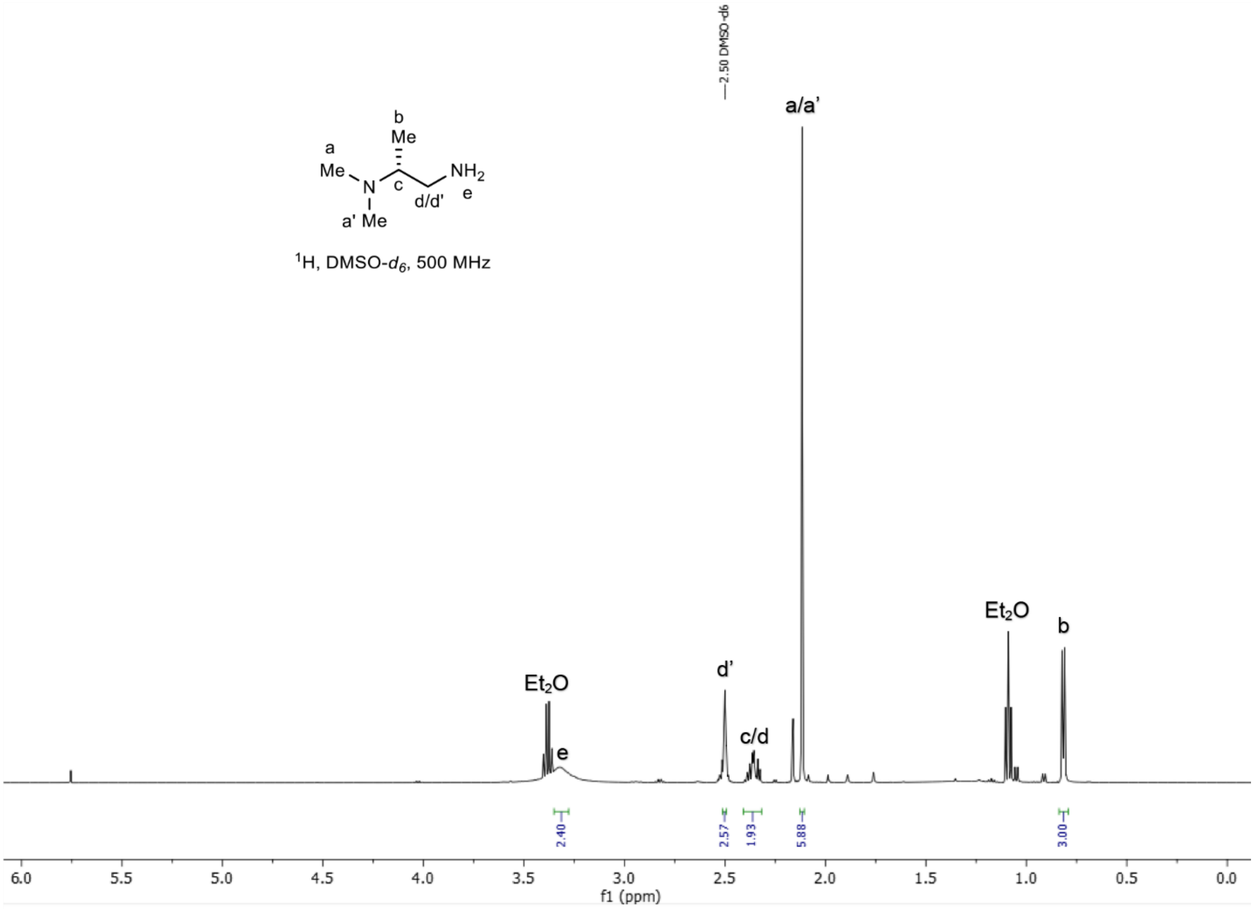

**b**

(*R*)-Dmp $^{13}$C NMR, DMSO-$d_6$, 126 MHz.

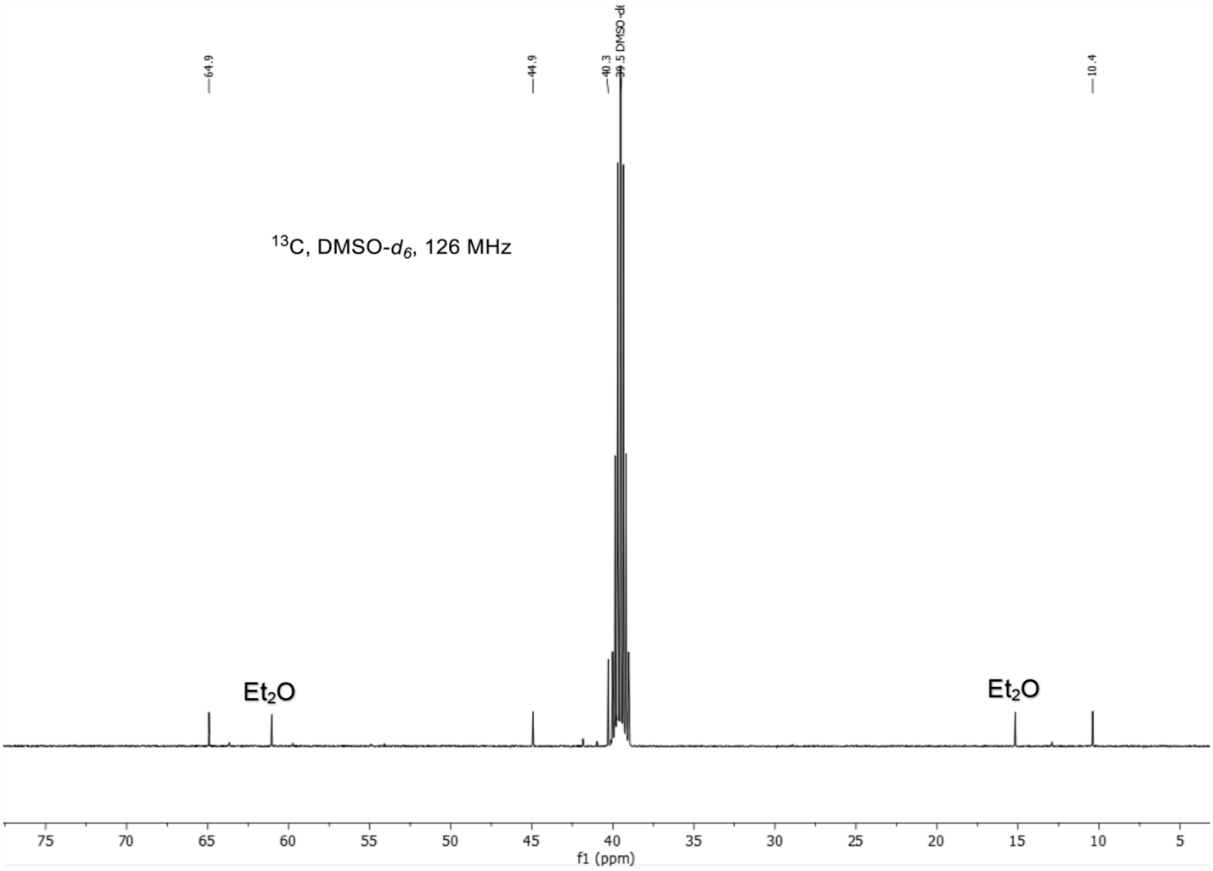

**c**

(*R*)-Dmp $^1$H-$^1$H COSY NMR, DMSO-$d_6$, 500 MHz.

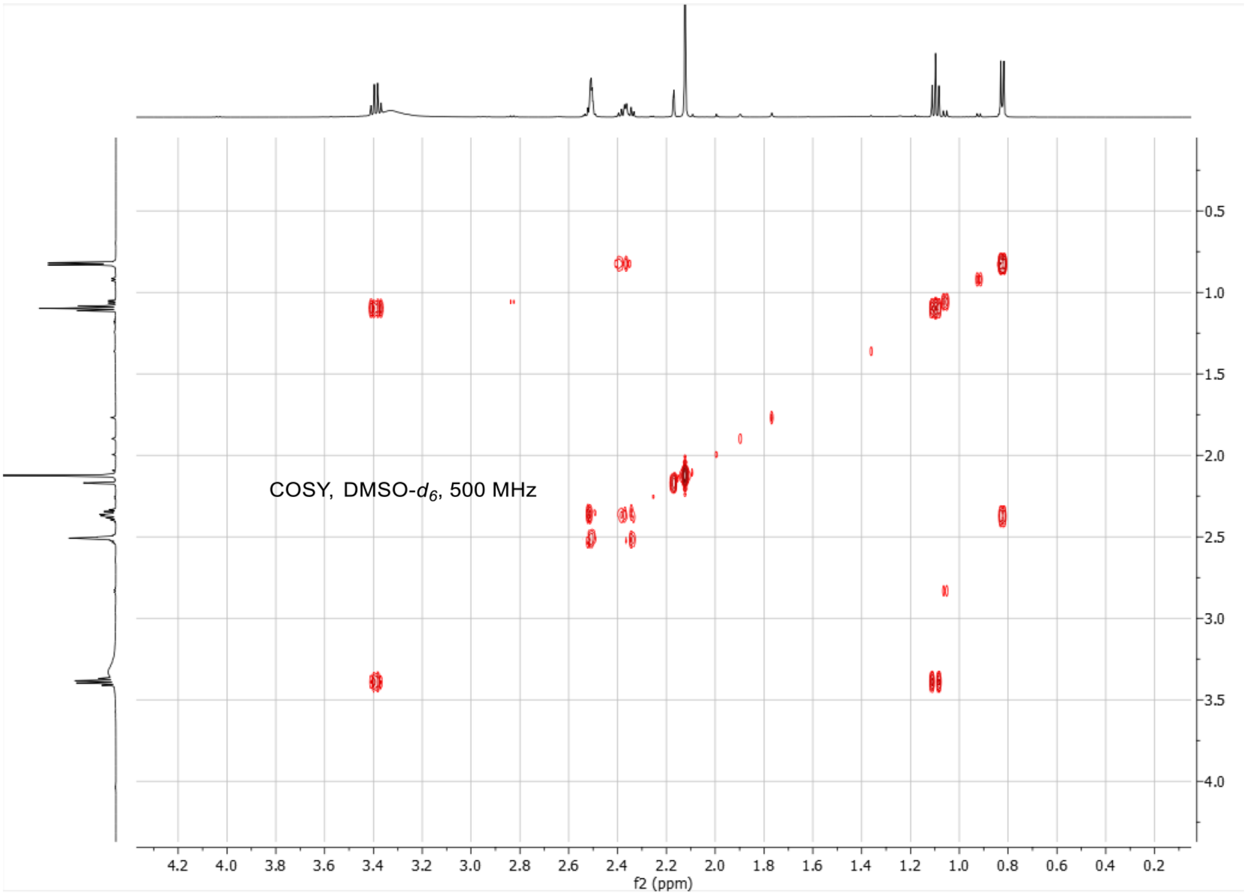

**d**

($S$)-Dmp $^1$H NMR, DMSO-$d_6$, 500 MHz.

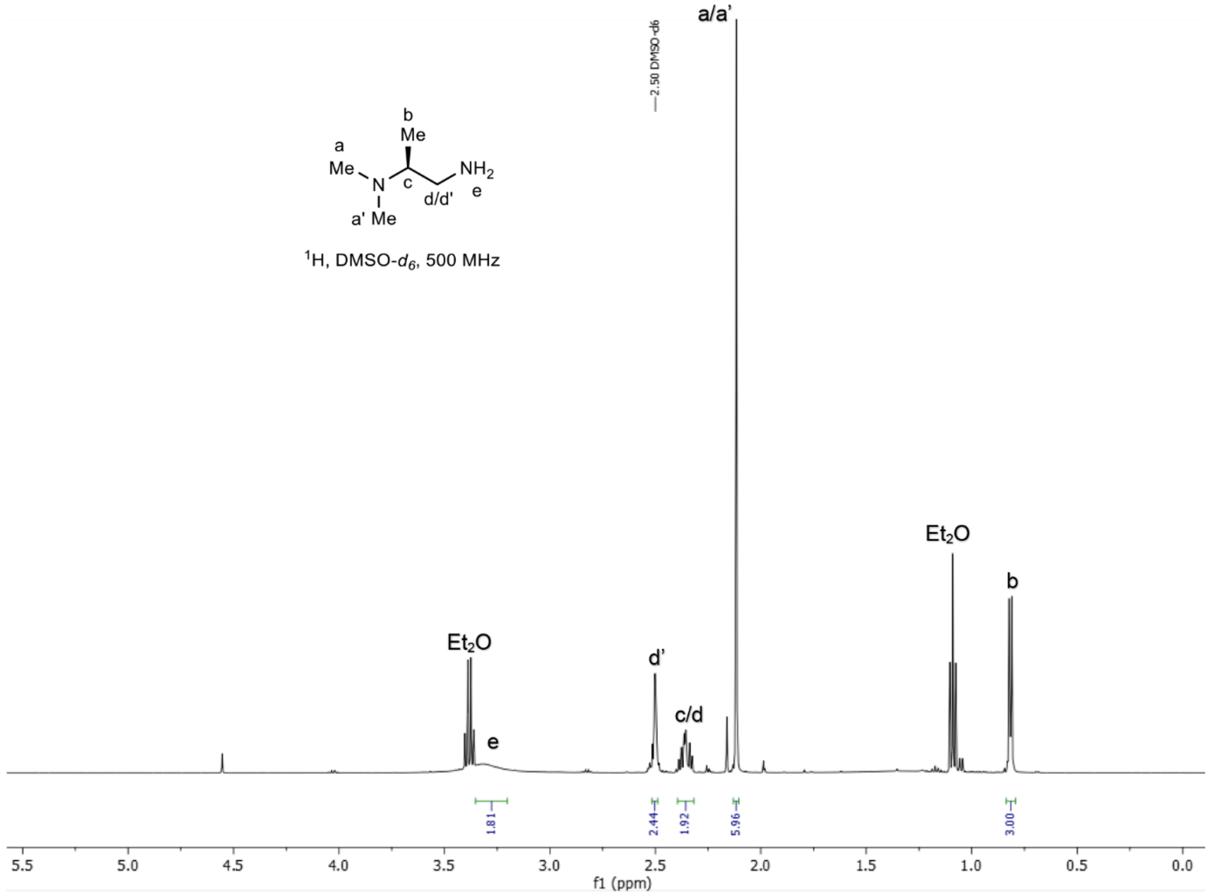

**e**

(*S*)-Dmp $^{13}C$ NMR, DMSO-$d_6$, 126 MHz.

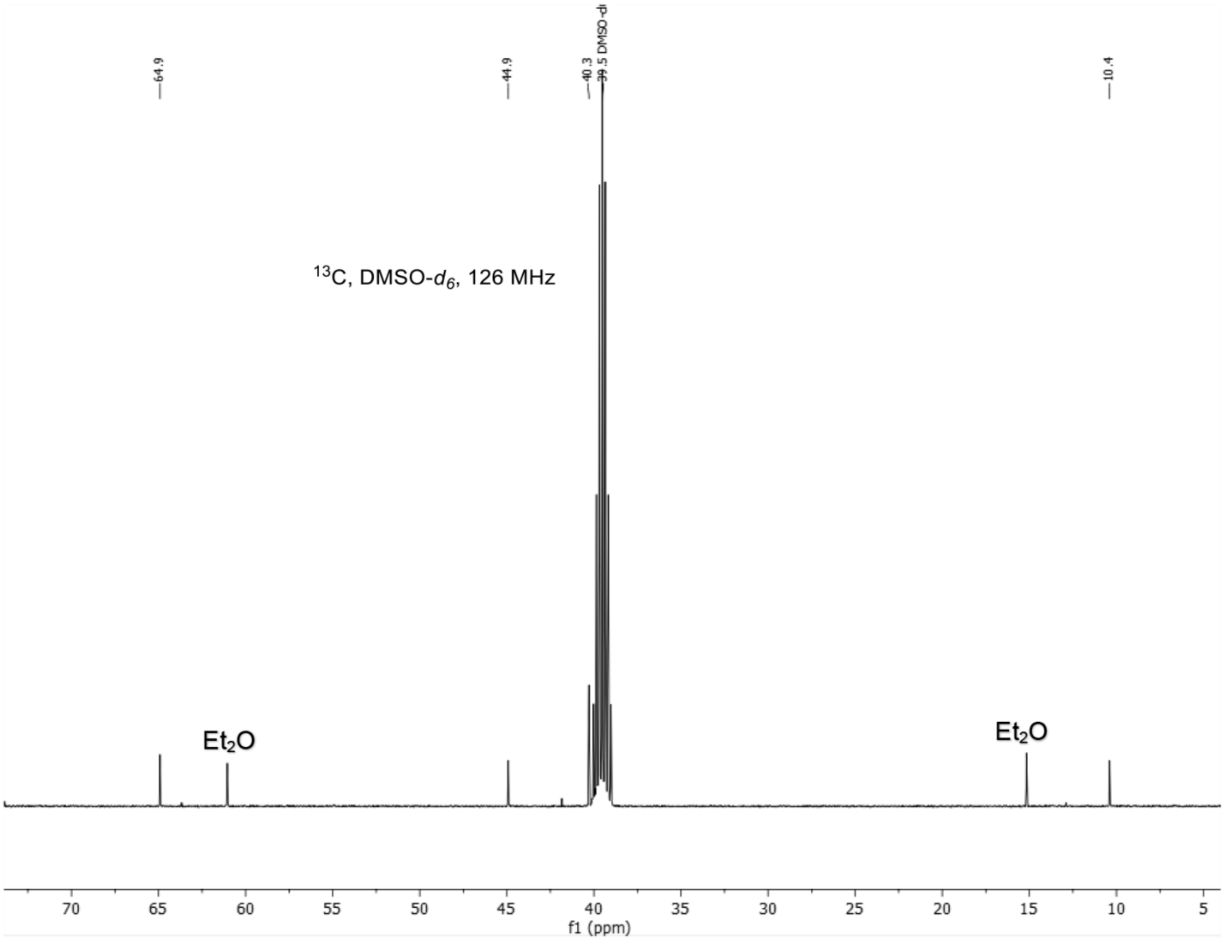

**f**

(*S*)-Dmp $^{1}$H-$^{1}$H COSY NMR, DMSO-$d_6$, 500 MHz.

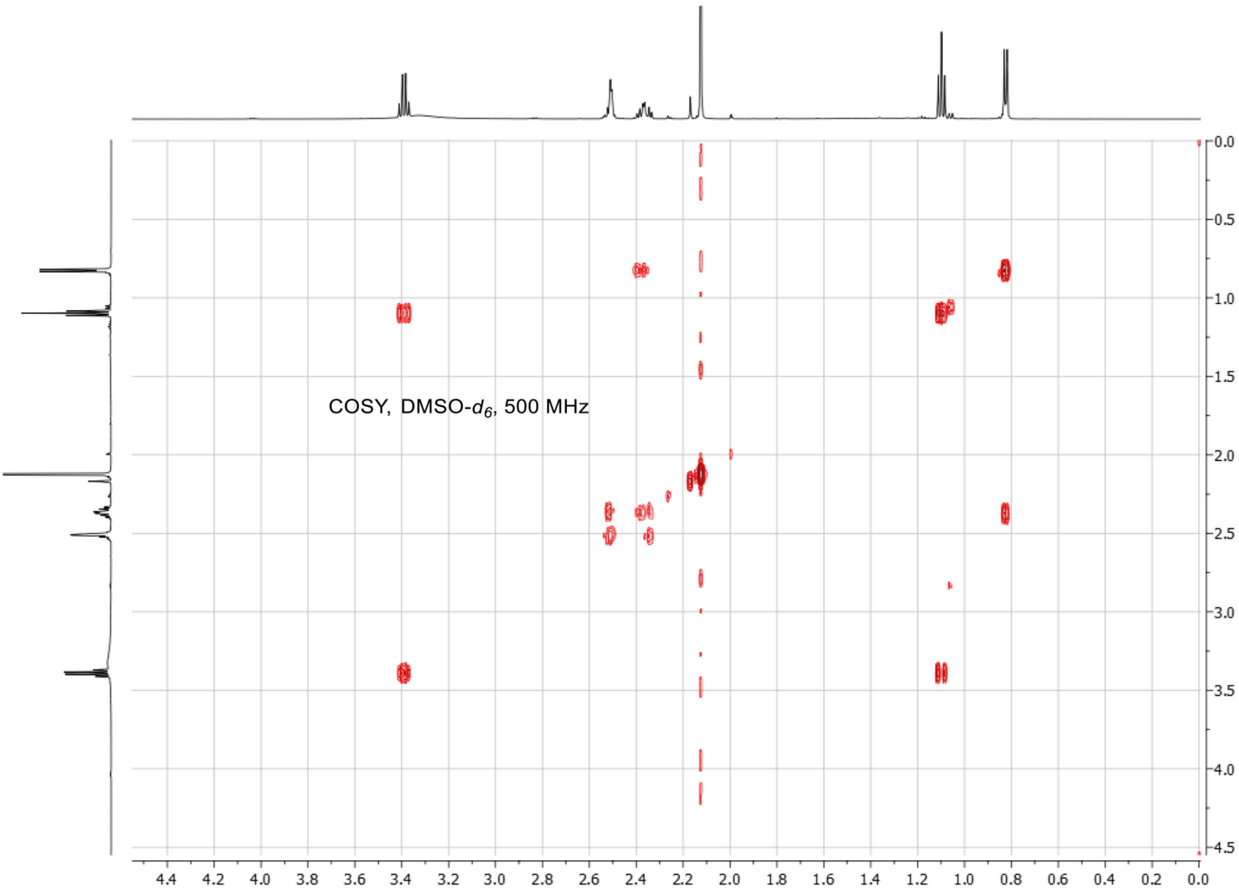

**Supplementary Figure 15: NMR spectra of authentic (*R*)-Dmp and (*S*)-Dmp standards**. Dmp: $N_2$,$N_2$-dimethyl-1,2-propanediamine. (**a**) (*R*)-Dmp $^{1}$H NMR, DMSO-$d_6$, 500 MHz. (**b**) (*R*)-Dmp $^{13}$C NMR, DMSO-$d_6$, 126 MHz. (**c**) (*R*)-Dmp $^{1}$H-$^{1}$H COSY NMR, DMSO-$d_6$, 500 MHz. (**d**) (*S*)-Dmp $^{1}$H NMR, DMSO-$d_6$, 500 MHz. (**e**) (*S*)-Dmp $^{13}$C NMR, DMSO-$d_6$, 126 MHz. (**f**) (*S*)-Dmp $^{1}$H-$^{1}$H COSY NMR, DMSO-$d_6$, 500 MHz.

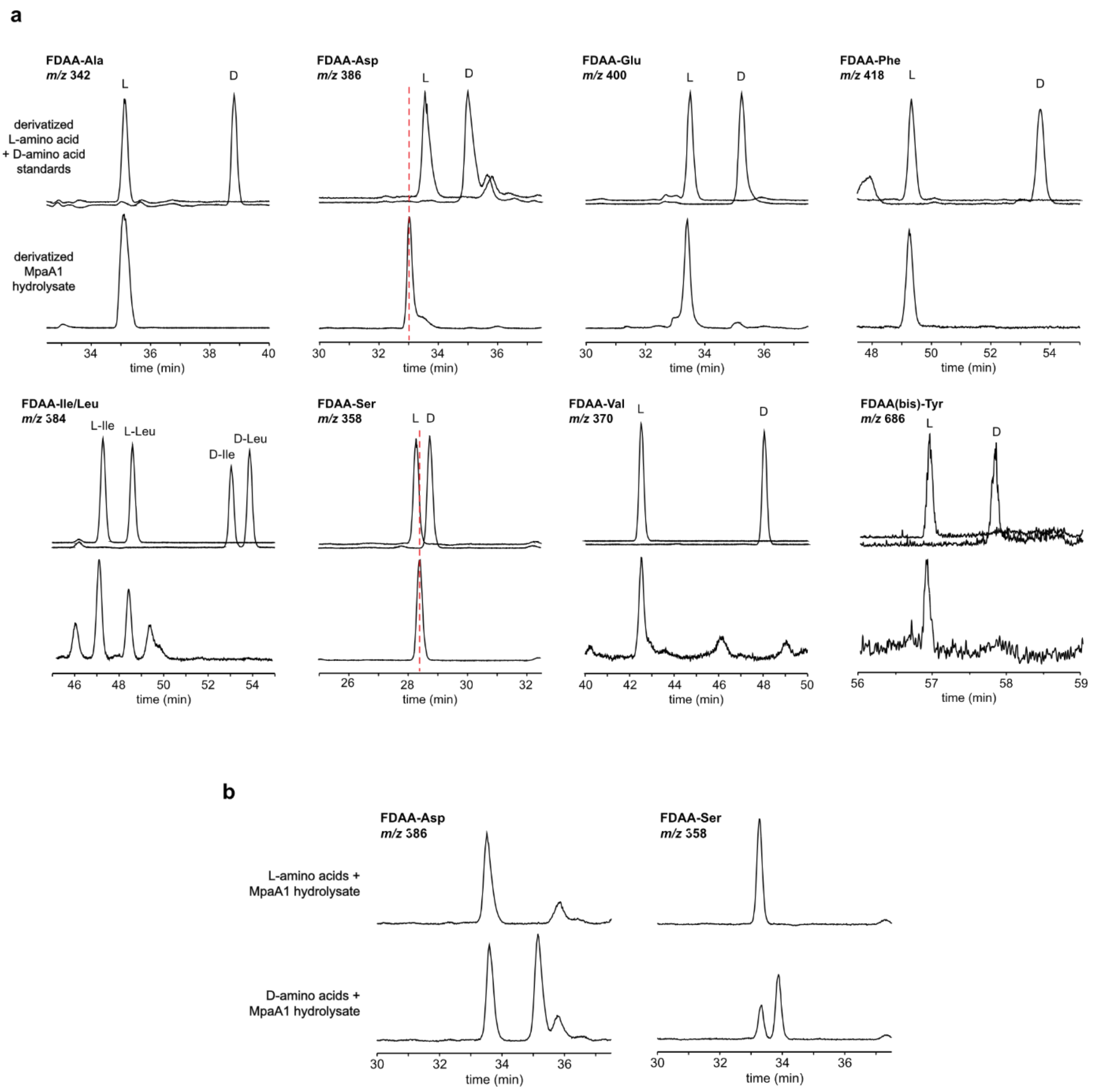

**Supplementary Figure 16: Determination of MpaA1 amino acid stereochemistry using LC-MS**. (**a**) Ion count-normalized LC-MS chromatograms are shown for derivatized amino acid standards and for 1-fluoro-2-4-dinitrophenyl-5-alanine amide (FDAA)-derivatized hydrolysate of **1**. L- and D-amino acid standards were run individually. FDAA-Trp adducts were not observed[9]. FDAA-Glu was monitored instead of FDAA-Gln due to the hydrolysis of MpaA1. Assignment of Asp and Ser was initially ambiguous owing to retention time drift. (**b**) LC-MS chromatograms for co-injections of derivatized amino acid standards with derivatized MpaA1 hydrolysate.

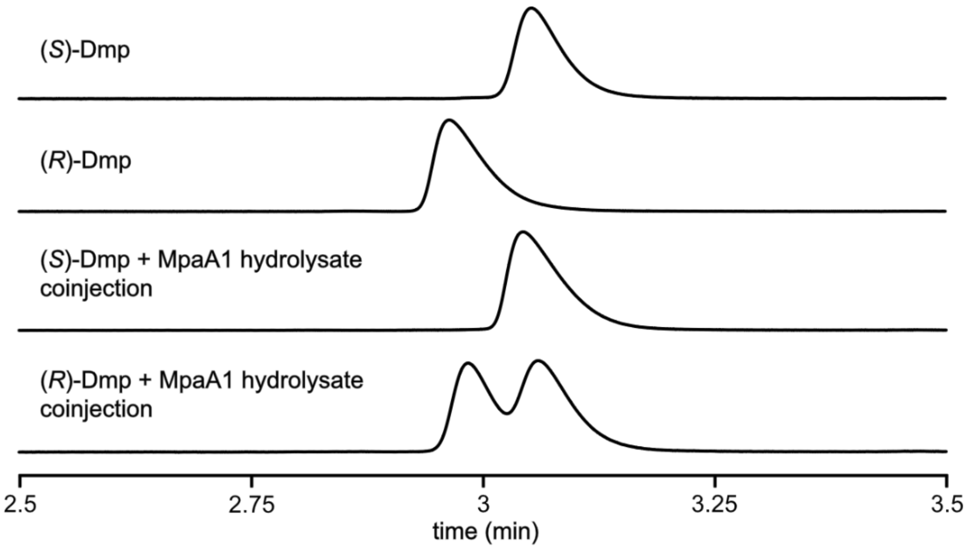

**Supplementary Figure 17: Determination of MpaA1 Dmp stereochemistry**. Absorbance-normalized chromatograms are shown for single injections of FDAA-derivatized (*R*)-Dmp and (*S*)-Dmp, along with co-injections of derivatized hydrolysate of **1**. Absorbance was monitored at 340 nm. Dmp: $N_2$,$N_2$-dimethyl-1,2-propanediamine.

**a**

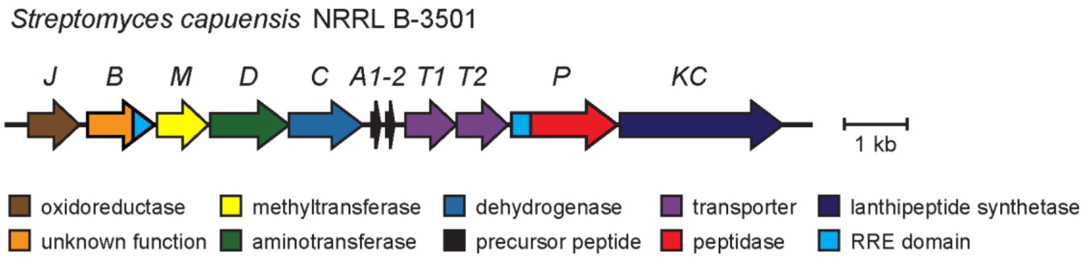

**b**

```
                Leader                              Core
        -20      -15      -10       -5      1    5        10       15       20       25       30
ScaA1  MQEIEKIQPVLELGMQELEAMDAPGWWTAAGVSAGVVVSASAAYGSAALSAAAIT
ScaA2  MQETEKIQPVLELGMQELEAMEAPGFWTGFSYGVAVSGTAAASAAVSVAASIAT
```

**c**

| Gene | NCBI Accession | Description (per NCBI listing) |
|---|---|---|
| *scaJ* | WP_003980804.1 | Luciferase-like monooxygenase (LLM) class flavin-dependent oxidoreductase |
| *scaB* | WP_003980803.1 | Hypothetical protein |
| *scaM* | WP_003980802.1 | Class I SAM-dependent methyltransferase |
| *scaD* | WP_003980801.1 | Aminotransferase class III-fold pyridoxal phosphate-dependent enzyme |
| *scaC* | WP_003980800.1 | Iron-containing alcohol dehydrogenase |
| *scaA1* | WP_165417107.1 | Hypothetical protein |
| *scaA2* | WP_003980799.1 | Hypothetical protein |
| *scaT1* | WP_003980797.1 | ATP-binding cassette domain-containing protein |
| *scaT2* | WP_003980796.1 | ABC transporter permease |
| *scaP* | WP_231512674.1 | Hypothetical protein |
| *scaKC* | WP_003980794.1 | Class III lanthionine synthetase LanKC |

**Supplementary Figure 18: The *sca* BGC identified from *Streptomyces capuensis* NRRL B-3501.** (**a**) Organization of the *sca* BGC. (**b**) Precursor peptide sequences. (**c**) Functional annotation and accession numbers of genes in the *sca* BGC

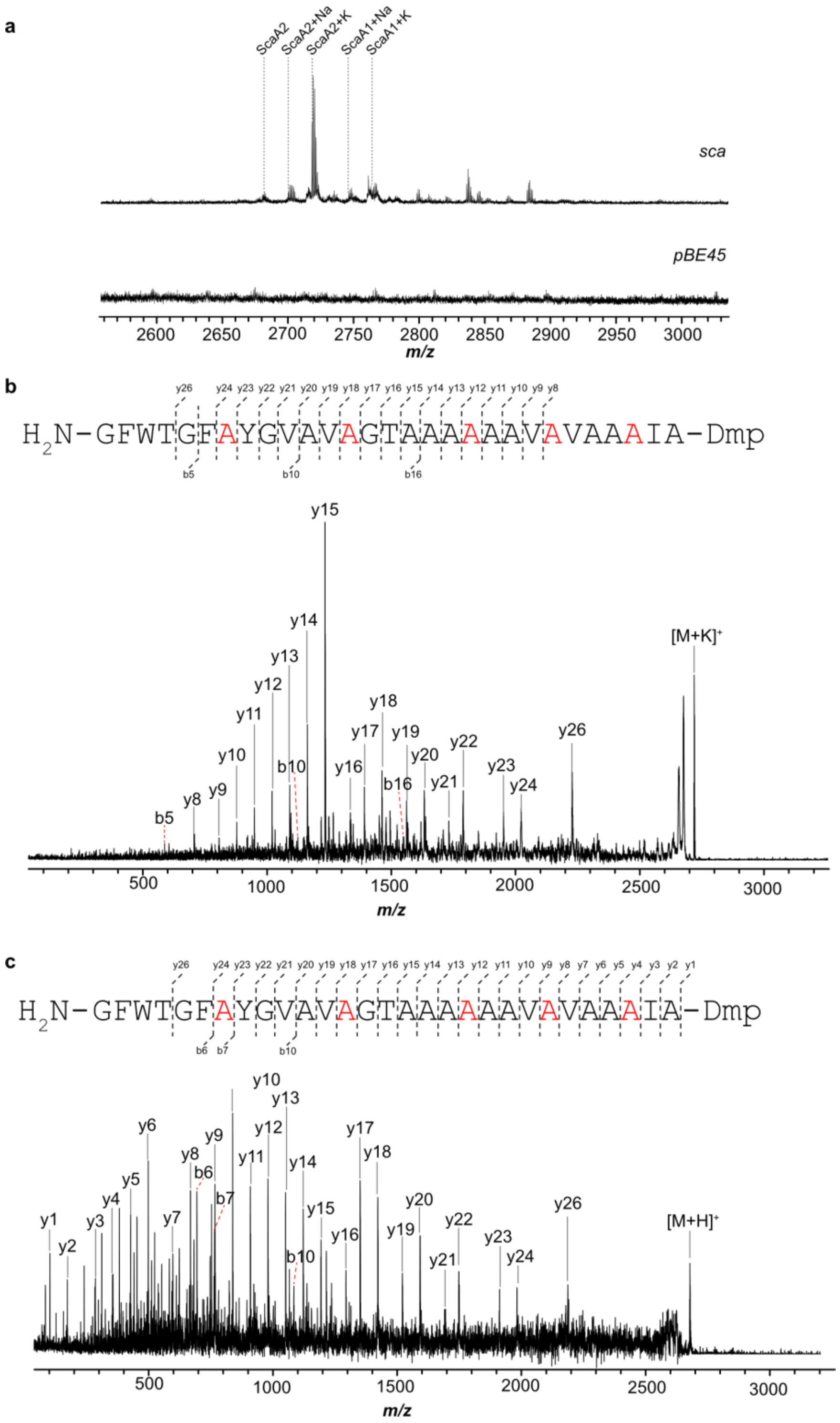

**Supplementary Figure 19: Heterologous expression and product characterization of *sca*. (a)** MALDI-TOF mass spectra of the methanol extracts of *S. albus* J1074 containing *sca* and the pBE45 empty vector. Sodium and potassium ions are denoted as '+Na' and '+K'. (**b**) MALDI LIFT-TOF/TOF mass spectrum of the modified ScaA2 core peptide acquired by the potassium adduct. Observed b- and y-ions are annotated. Ions are in the +1 charge state. Suspected sites of conversion to *D*-Ala are colored in red. Dmp: $N_2$,$N_2$-dimethyl-1,2-propanediamine. (**c**) MALDI LIFT-TOF/TOF mass spectrum of the modified ScaA2 core peptide acquired using the $[M+H]^+$ ion. Observed b- and y-ions are annotated. Ions are in the +1 charge state. Suspected sites of conversion to *D*-Ala are colored in red. Dmp: $N_2$,$N_2$-dimethyl-1,2-propanediamine.

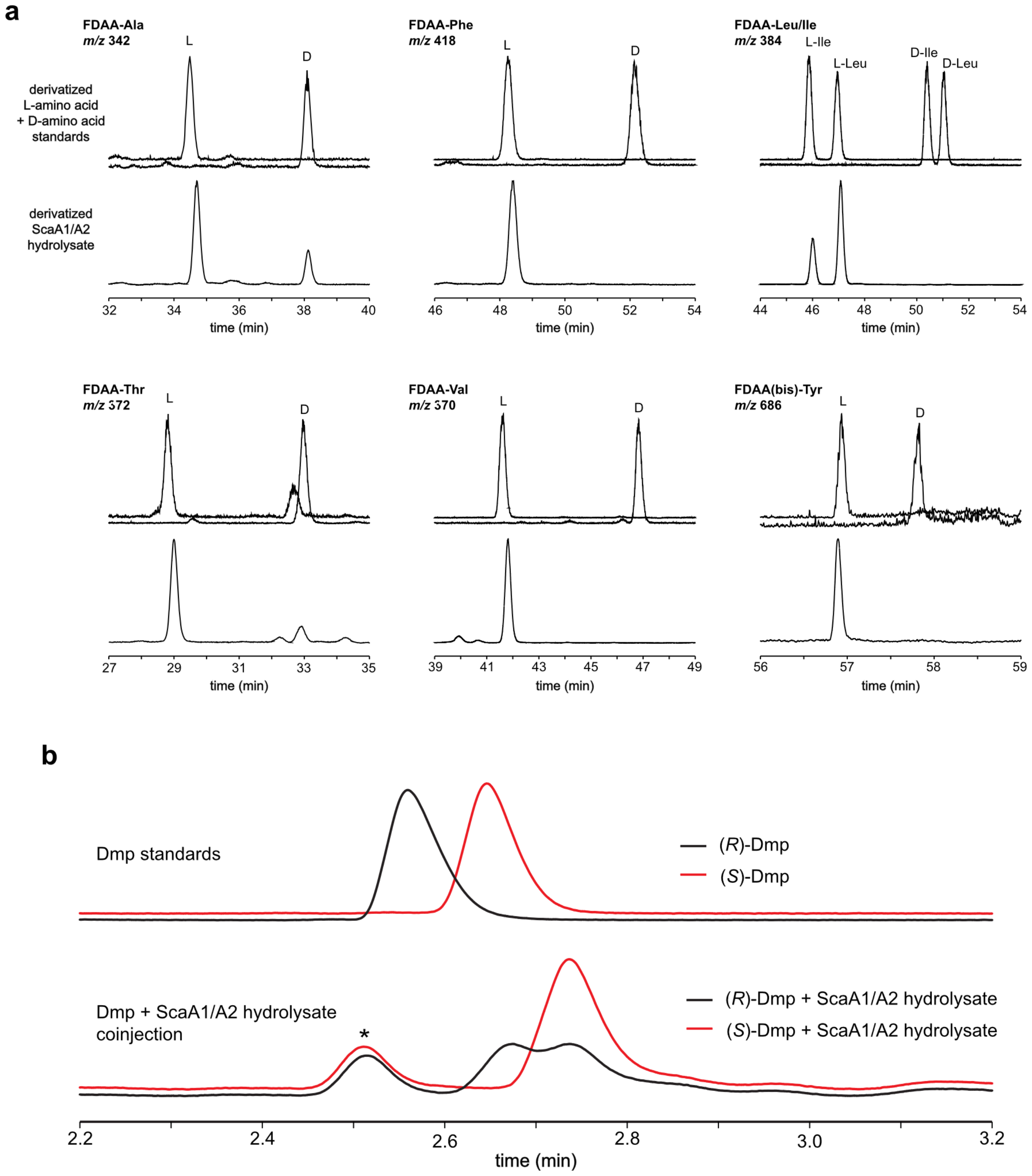

**Supplementary Figure 20: Determination of ScaA1/A2 residue stereochemistry**. (**a**) Ion count-normalized LC-MS chromatograms are shown for derivatized amino acid standards and for 1-fluoro-2-4-dinitrophenyl-5-alanine amide (FDAA)-derivatized hydrolysate of ScaA1/A2. L- and D-amino acid standards were run individually. FDAA-Trp adducts were not observed[9]. (**b**) Absorbance-normalized chromatograms are shown for single injections of FDAA-derivatized (*R*)-Dmp and (*S*)-Dmp, along with co-injections of derivatized hydrolysate of ScaA1/A2. Absorbance was monitored at 340 nm. Asterisk indicates possible FDAA adduct to unidentified hydrolysis product. Dmp: $N_2$,$N_2$-dimethyl-1,2-propanediamine.

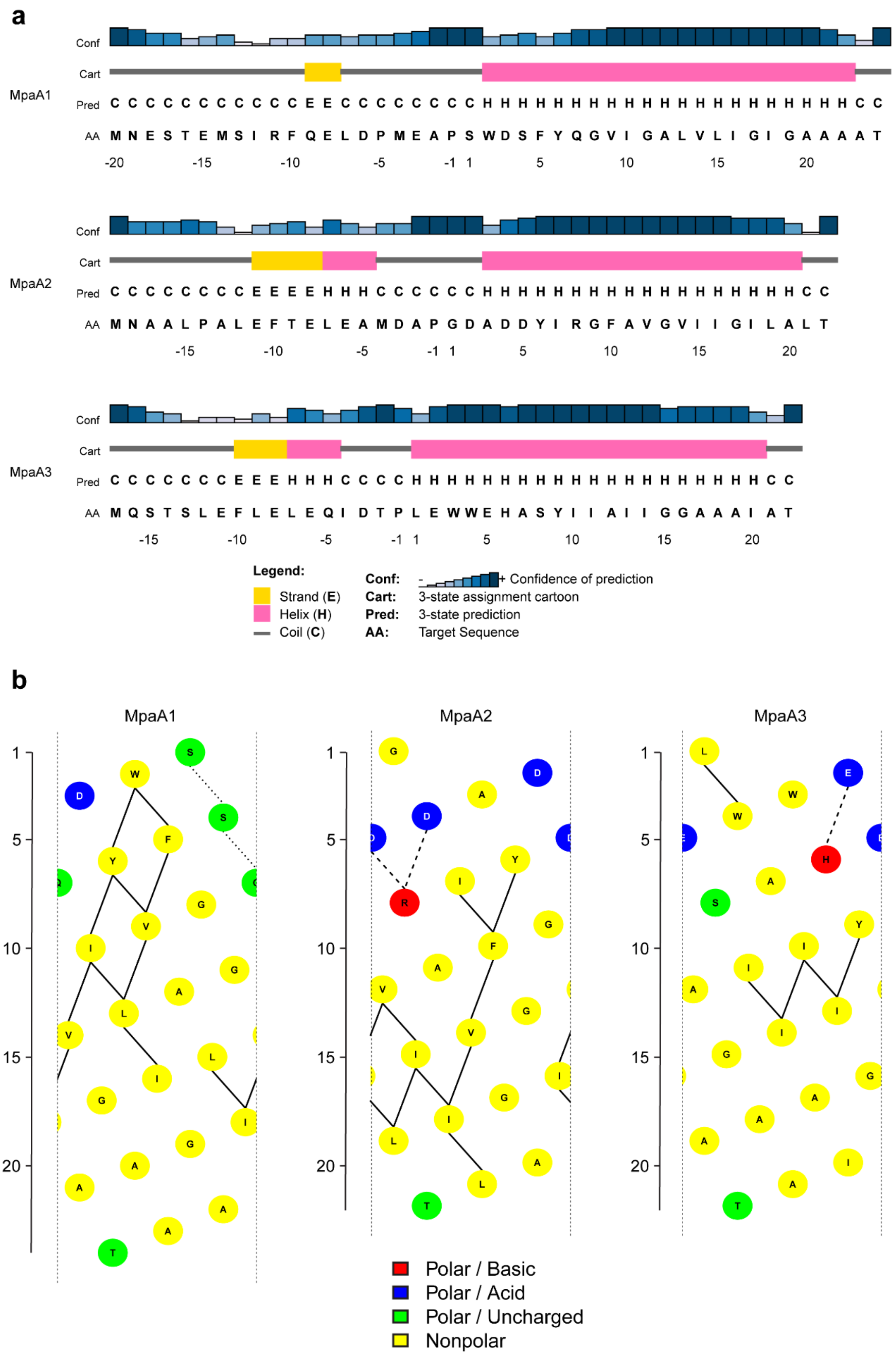
a
MpaA1
MpaA2
MpaA3
Conf
Cart
Pred
AA
Legend:
Strand (E)
Helix (H)
Coil (C)
Conf: Confidence of prediction
Cart: 3-state assignment cartoon
Pred: 3-state prediction
AA: Target Sequence
b
MpaA1
MpaA2
MpaA3
Polar / Basic
Polar / Acid
Polar / Uncharged
Nonpolar

c

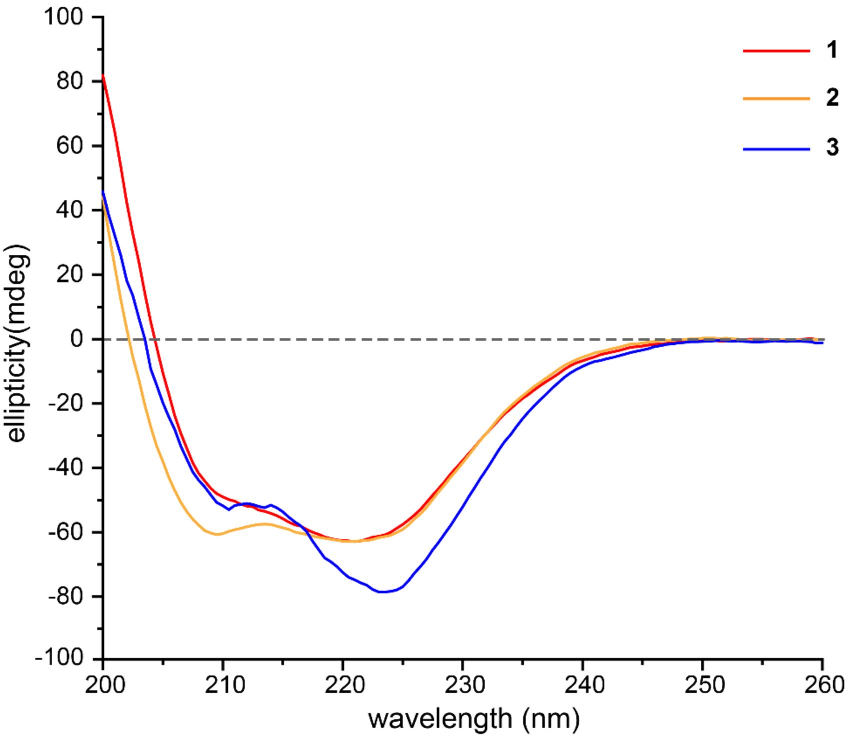

**Supplementary Figure 21: Secondary structure of daptides 1-3**. (**a**) α-helices of MpaA1-3 predicted by PSI-PRED4.0[10] (http://bioinf.cs.ucl.ac.uk/psipred/). (**b**) Helical net diagrams showing predicted intra-helix interactions for MpaA1-3 core peptides. Diagrams were generated using NetWheels (http://lbqp.unb.br/NetWheels/). (**c**, following page) Circular dichroism spectra of **1**-**3**.

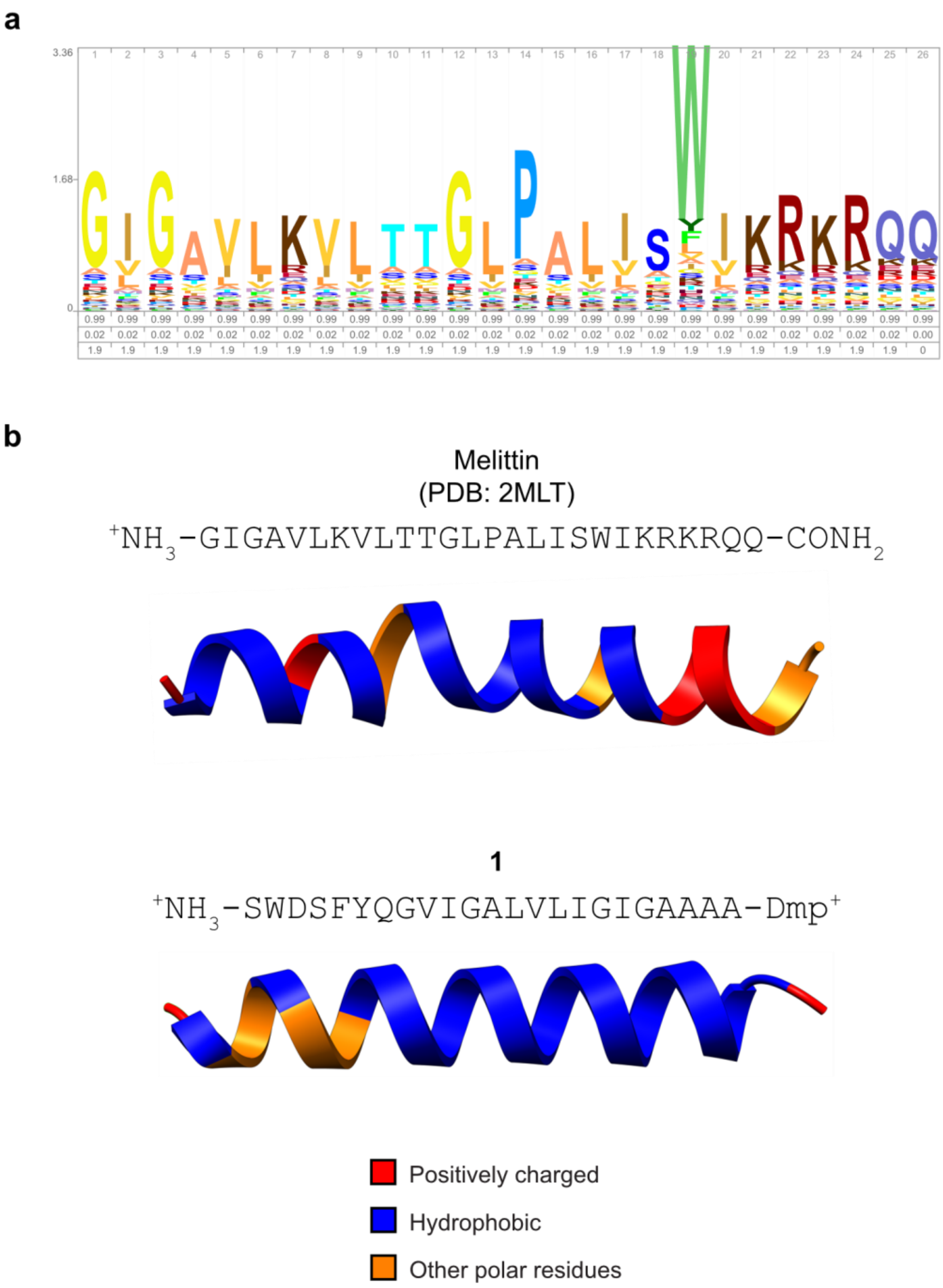

**Supplementary Figure 22: Comparison of melittin to daptide 1.** (**a**) Sequence logo for the melittin family is shown. The logo was generated using SkyLign[6] on the melittin pHMM (Pfam identifier: PF01372)[2]. (**b**) A crystal structure of melittin and predicted structure of **1** are shown. AlphaFold[11] was used to predict the MpaA1 complete sequence, and only the core region is shown. Melittin structure was re-created based on a review article[12]. Dmp: $N_2$,$N_2$-dimethyl-1,2-propanediamine.

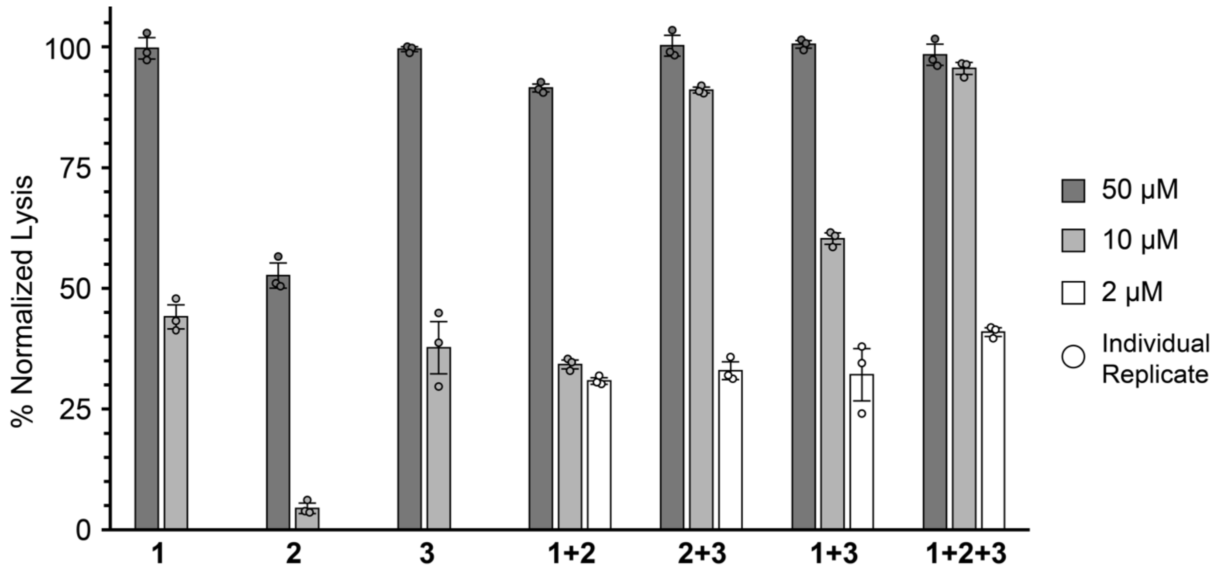

**Supplementary Figure 23: Hemolytic activity of daptides 1-3**. Hemolysis of bovine erythrocytes was normalized to the positive control [5% (*v/v*) Triton X-100]. All individual and combination daptide treatments were assessed. The percentage of hemolysis at total concentrations of 50 µM and 10 µM are shown as grey and white bars, respectively. In the combination treatments, the total daptide concentration is listed (i.e., each daptide contributes 1/2 or 1/3 of the total concentration). All data represent the mean of n = 3 biologically independent samples and error bars show standard deviation.

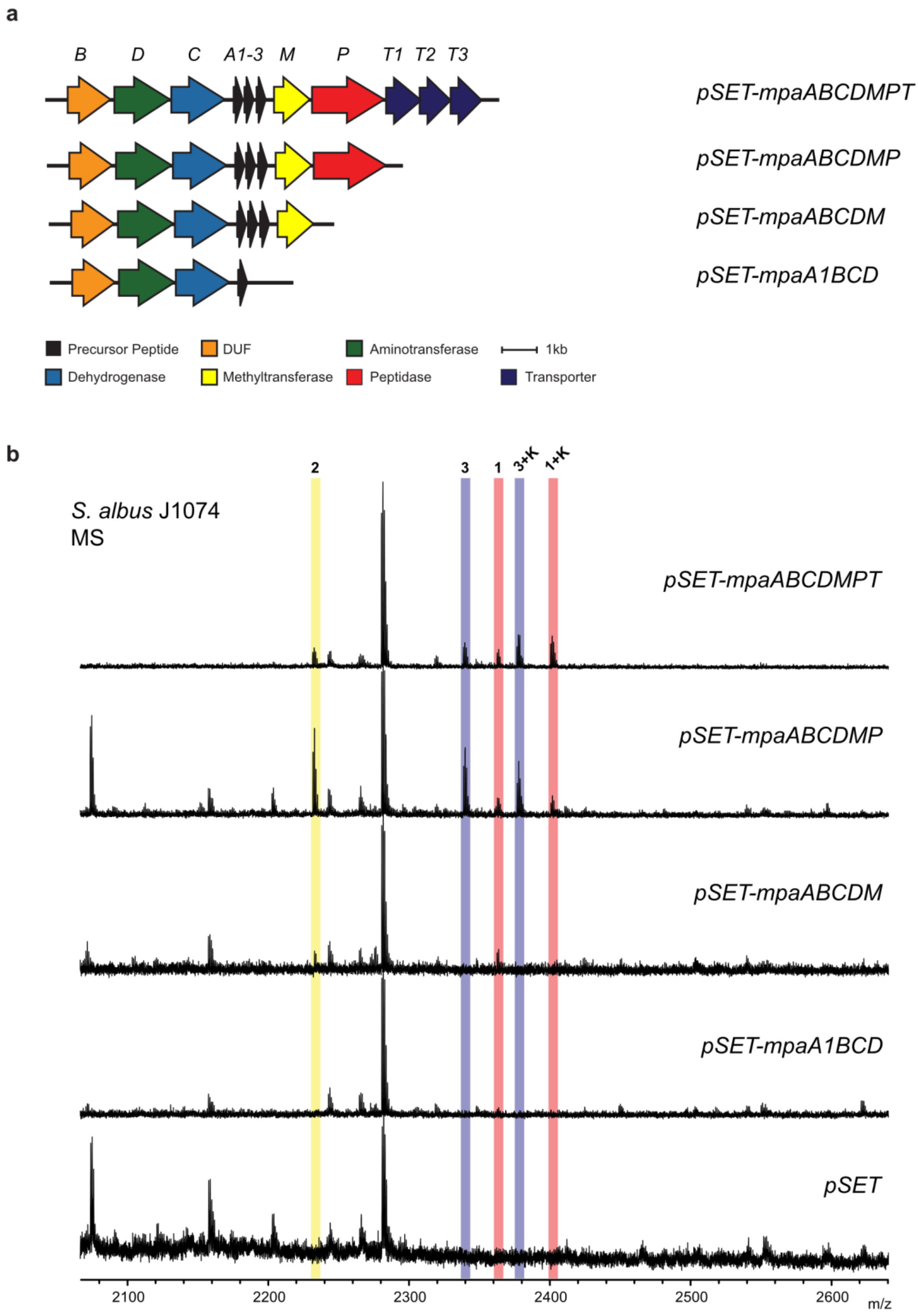

**Supplementary Figure 24: Heterologous expression and *mpa* gene omissions in *S. albus* J1074**. (**a**) Gene organization of the minimal *mpa* BGC (no flanking genes included) with examined gene omissions depicted. (**b**) MALDI-TOF mass spectra of the corresponding methanol extracts.

**a**

| Condition | Medium | Induction Density $OD_{600}$ | Expression Time (h) | Expression Temperature (°C) |
|---|---|---|---|---|
| 1 | TB | 1.0 | 20 | 18 |
| 2 | TB | 1.0 | 3 | 18 |
| 3 | TB | 1.5 | 3 | 37 |
| 4 | LB | 0.6 | 20 | 18 |
| 5 | M9 | 0.7 | 36 | 18 |

**b**

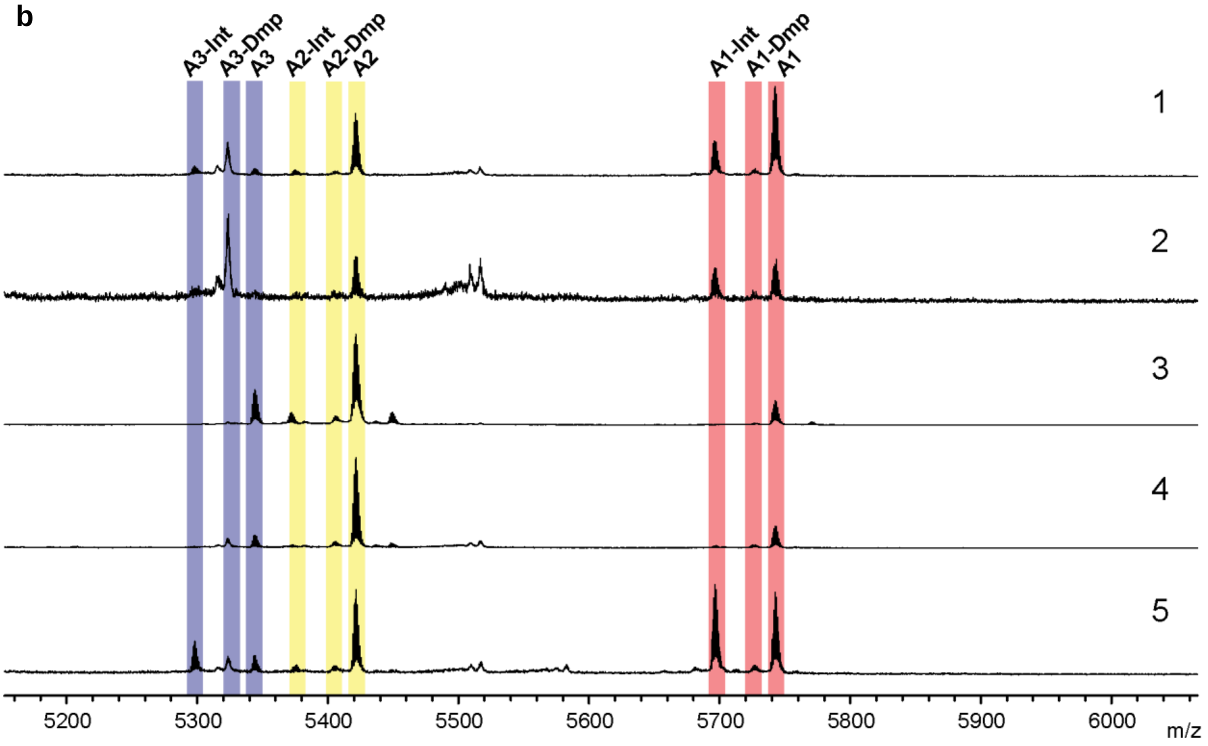

**Supplementary Figure 25: Expression of the refactored *mpaABCDE* under various cultivation conditions**. (**a**) List of variables for expression conditions 1-5, which used IPTG (0.5 mM final) for induction . (**b**) MALDI-TOF mass spectra of IMAC-purified products from expression conditions 1-5. The ketone/amine intermediates are denoted by "-Int" while the fully modified products are denoted by "-Dmp". Dmp: $N_2$,$N_2$-dimethyl-1,2-propanediamine

**a**

| Plasmid | Chaperone Genes | Promoter | Inducer |
|---|---|---|---|
| pTf16 | *tig* | araB | L-Arabinose: 0.5 mg/ml |
| pG-KJE8 | *dnaK-dnaJ-grpE*<br>*gro ES-groEL* | araB<br>Pzt-1 | L-Arabinose: 0.5 mg/ml<br>Tetracycline: 1 ng/ml |
| pKJE7 | *dnaK-dnaJ-grpE* | araB | L-Arabinose: 0.5 mg/ml |
| pGro7 | *groES-groEL* | araB | L-Arabinose: 0.5 mg/ml |
| pG-Tf2 | *groES-groEL-tig* | Pzt-1 | Tetracycline: 1 ng/ml |

**b**

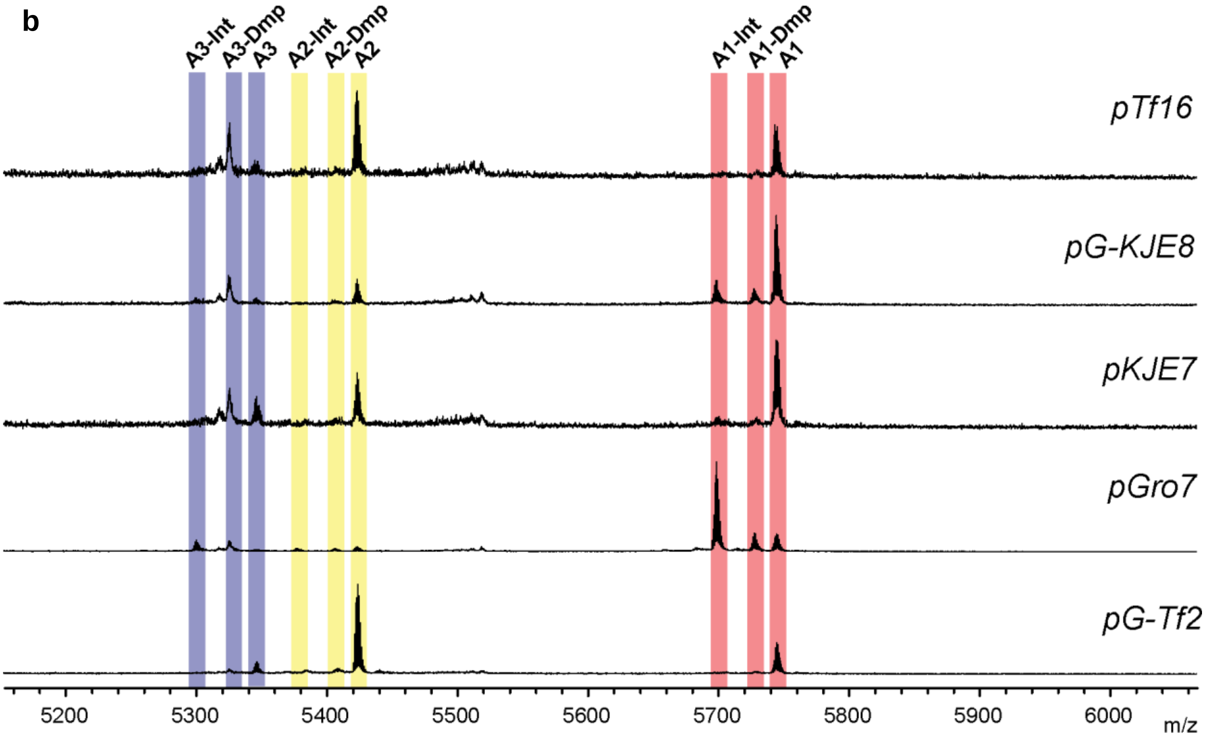

**Supplementary Figure 26: Expression of the refactored *mpaABCDE* with chaperone plasmids**. (**a**) Information for the genes involved in each plasmid. All chaperone plasmids were obtained from TaKaRa Bio Inc. Additional expression parameters were: medium, TB; induction $OD_{600}$, 0.8; [IPTG] 0.1 mM; expression temperature, 18 °C; expression time, 18 h. (**b**) MALDI-TOF mass spectra of IMAC purified products. The ketone/amine intermediates are denoted by "-Int" while the fully modified products are denoted by "-Dmp". Dmp: $N_2$,$N_2$-dimethyl-1,2-propanediamine.

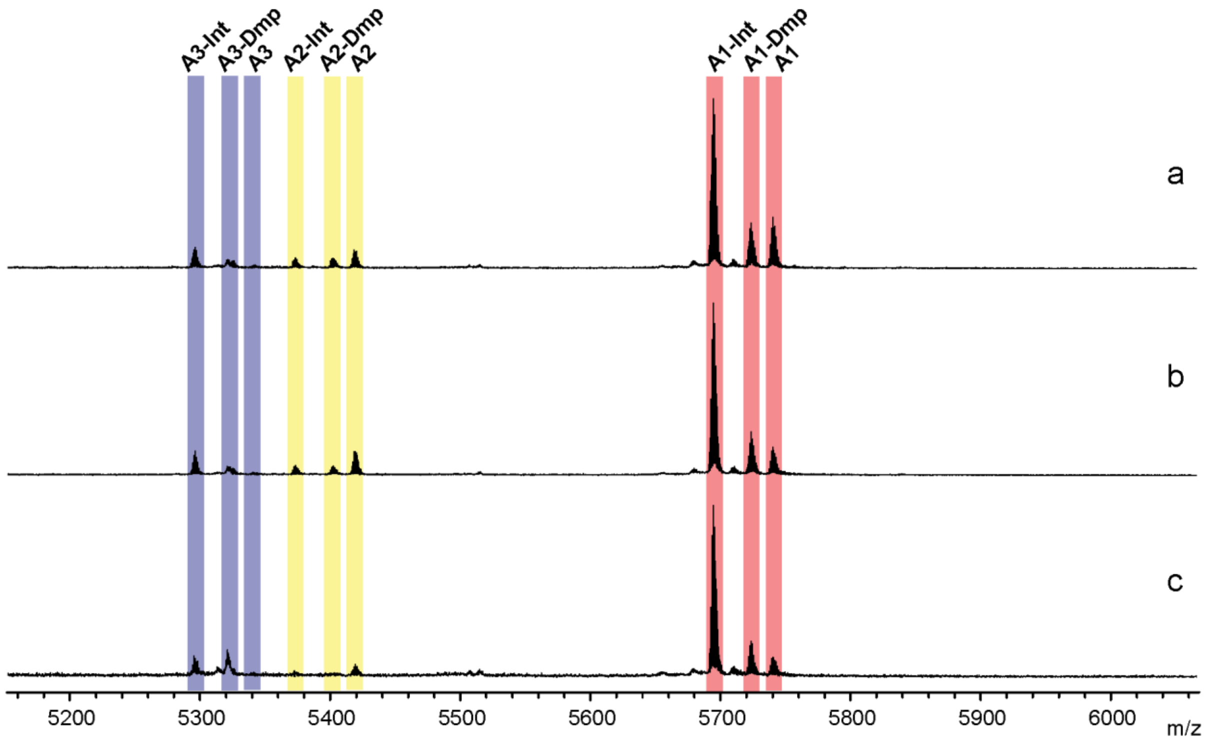

**Supplementary Figure 27: Expression of the refactored *mpaABCDE* with pGro7 chaperone plasmid under various arabinose concentrations** (a = 0.5 mg/mL, b = 2.0 mg/mL, c = 4.0 mg/mL). Additional expression parameters were: medium, TB; induction $OD_{600}$, 0.8; [IPTG], 0.1 mM; expression temperature, 18 °C; expression time, 18 h. The ketone/amine intermediates are denoted by "-Int" while the fully modified products are denoted by "-Dmp". Dmp: $N_2$,$N_2$-dimethyl-1,2-propanediamine.

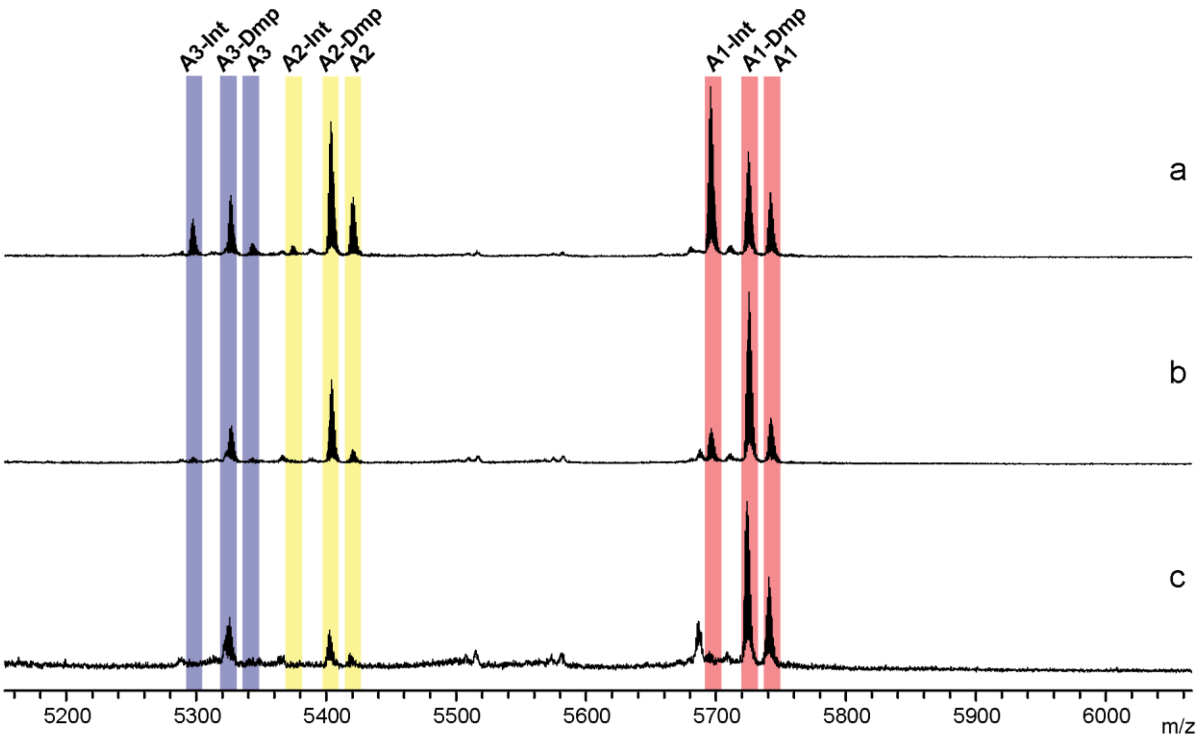

**Supplementary Figure 28: Expression of the refactored *mpaABCDE* with pGro7 chaperone plasmid with extended expression time** (a = 36 h; b = 72 h; c = 96 h). Additional expression parameters were: medium, M9; [arabinose], 0.5 mg/ml; induction $OD_{600}$, 0.7; [IPTG], 0.5 mM; expression temperature, 18 °C. The ketone/amine intermediates are denoted by "-Int" while the fully modified products are denoted by "-Dmp". Dmp: $N_2$,$N_2$-dimethyl-1,2-propanediamine.

**a**

-25 -20 -15 -10 -5 -1 1 5 10 15 20

MHHHHHHAMMNESTEMSIRFQ**ELDPME**A**P**SWDSFYQGVIGALVLIGIGAAAAT

**b**

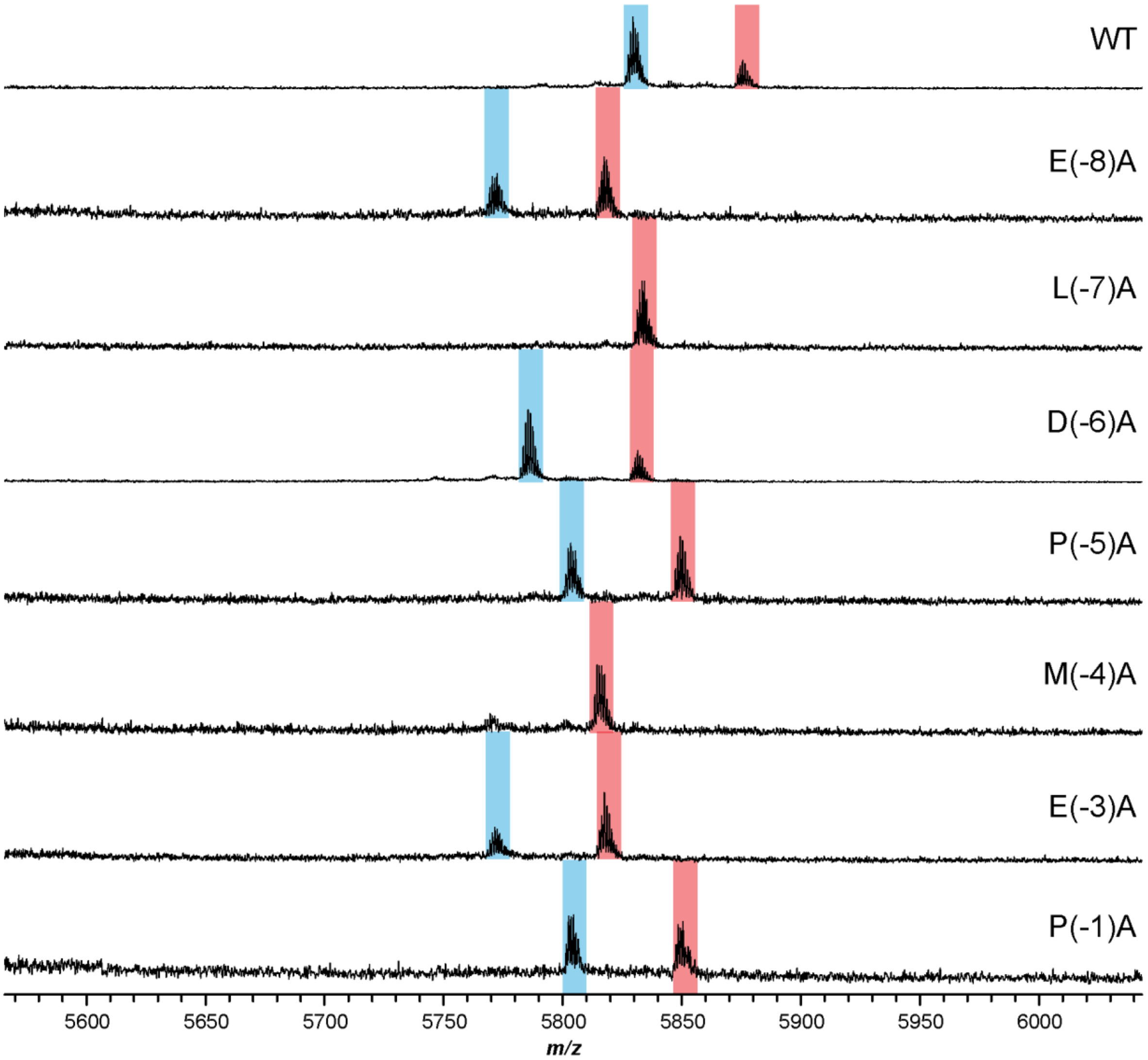

**Supplementary Figure 29: MALDI-TOF-MS analysis of MpaA1 leader region variants co-expressed with MpaB and MpaC.** (**a**) Sequence of MpaA1 with varied residues shown in bold. (**b**) MALDI-TOF mass spectra of IMAC-purified products after co-expression under the following condition: medium, TB; chaperone plasmid, pGro7; [arabinose], 0.5 mg/mL; induction $OD_{600}$, 0.8; [IPTG], 0.1 mM; expression temperature, 18 °C; expression time, 18 h. Ions for unmodified and modified peptides are highlighted in red and blue, respectively.

**a**

-25 -20 -15 -10 -5 -1 1 5 10 15 20

MHHHHHHAMMNESTEMSIRFQELDPMEAPSWDSFYQGVIGALVLIGIGAAAA**T**

**b**

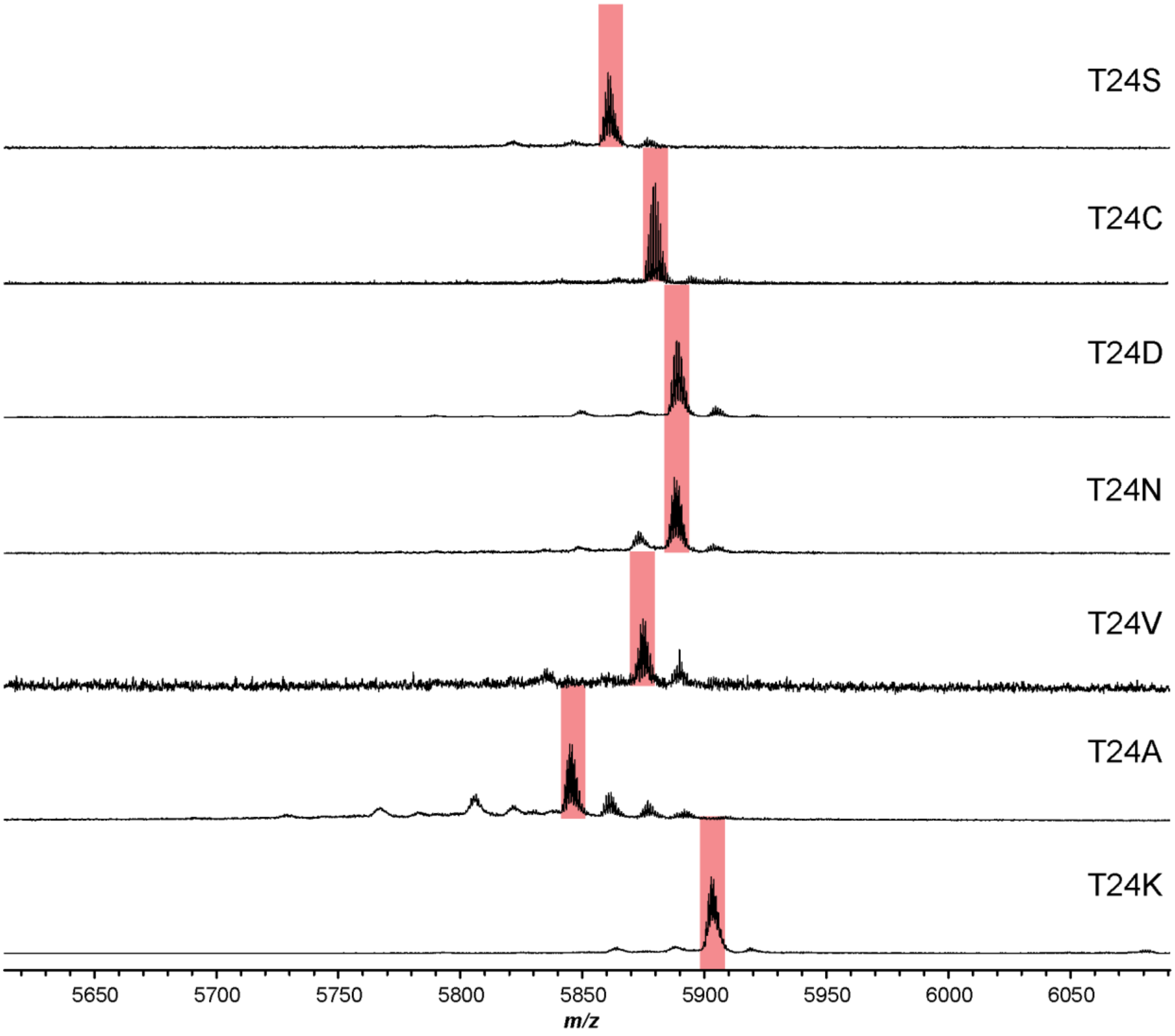

**Supplementary Figure 30: MALDI-TOF-MS analysis of MpaA1 *C*-terminal Thr variants that were co-expressed with MpaB and MpaC**. (**a**) Sequence of MpaA1 with the replaced C-terminal Thr shown in bold. (**b**) MALDI-TOF mass spectra of IMAC-purified products after co-expression under the following condition: medium, M9; chaperone plasmid, pGro7; [arabinose], 0.5 mg/mL; induction $OD_{600}$, 0.7; [IPTG], 0.5 mM; expression temperature, 18 °C; expression time, 72 h. Ions for unmodified peptides are highlighted in red. Minor peaks appearing at +16 are presumed to be from Met oxidation.

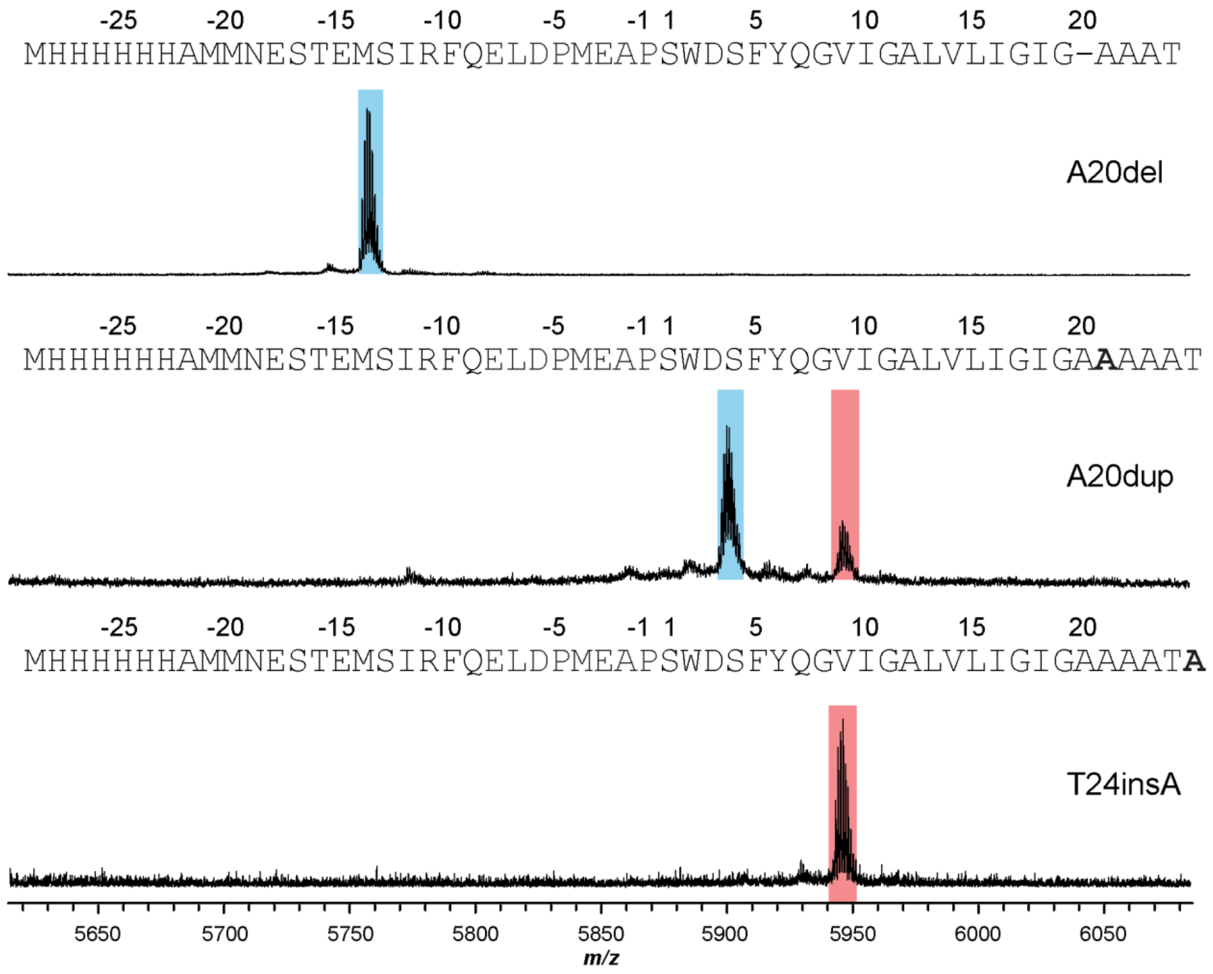

**Supplementary Figure 31: MALDI-TOF-MS analysis of MpaA1 insertion and deletion variants at the C-terminus that were co-expressed with MpaB and MpaC.** Products were purified by IMAC after co-expression under the following condition: medium, M9; chaperone plasmid, pGro7; [arabinose], 0.5 mg/mL; induction $OD_{600}$, 0.7; [IPTG], 0.5 mM; expression temperature, 18 °C; expression time, 72 h. Sequences of the MpaA1 variants are shown above their corresponding mass spectra, with the inserted/deleted residues denoted in bold and dash respectively. Ions for unmodified and modified peptides are highlighted in red and blue, respectively.

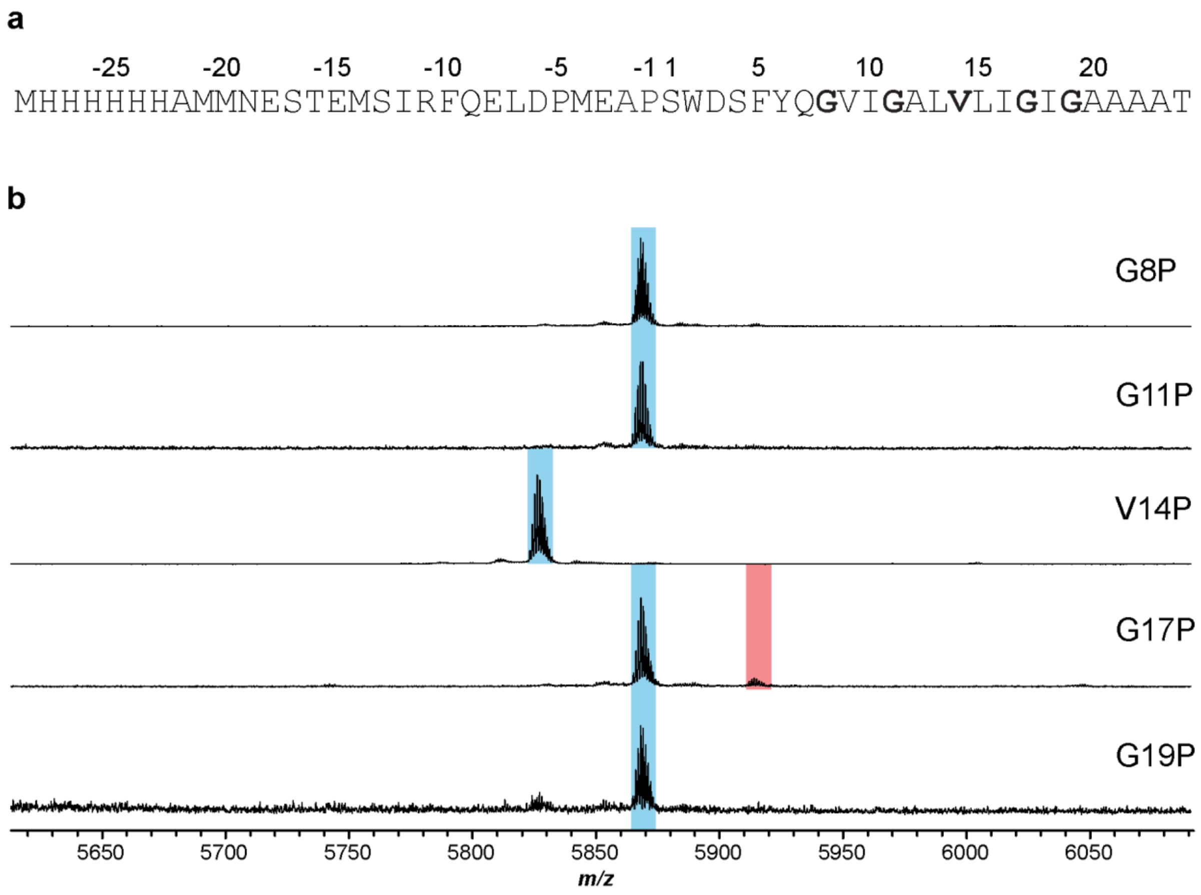

**Supplementary Figure 32: MALDI-TOF-MS analysis of MpaA1 variants in the core peptide region that were co-expressed with MpaB and MpaC**. (**a**) Sequence of MpaA1 with the mutated residues shown in bold. (**b**) MALDI-TOF mass spectra of IMAC-purified products after co-expression under the following condition: medium, M9; chaperone plasmid, pGro7; [arabinose], 0.5 mg/mL; induction $OD_{600}$, 0.7; [IPTG], 0.5 mM; expression temperature, 18 °C; expression time, 72 h. Ions for unmodified and modified peptides are highlighted in red and blue, respectively.

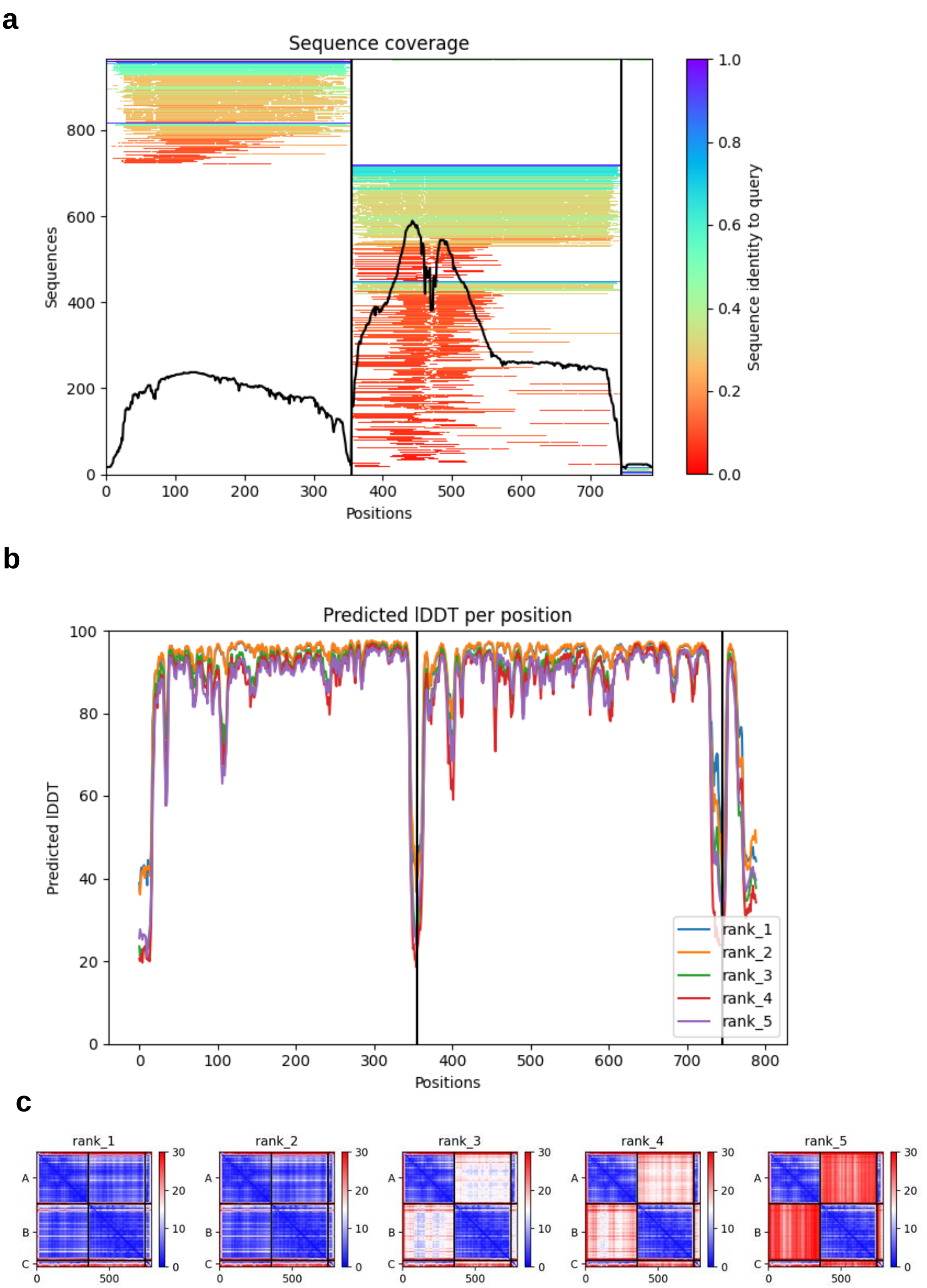

**Supplementary Figure 33: Assessment of the AlphaFold-Multimer prediction for the MpaB-MpaC-MpaA1 complex**. (**a**) The multiple sequence alignment (MSA) depth during MpaB-MpaC-MpaA1 complex prediction using AlphaFold- Multimer. (**b**) The predicted Local Distance Difference Test (pLDDT) for five predicted models. pLDDT is a per-residue measure of local confidence on a scale from 0-100. (**c**) Predicted Aligned Error (PAE) for five predicted models. The inter PAE between chains is low, indicating a confident prediction.

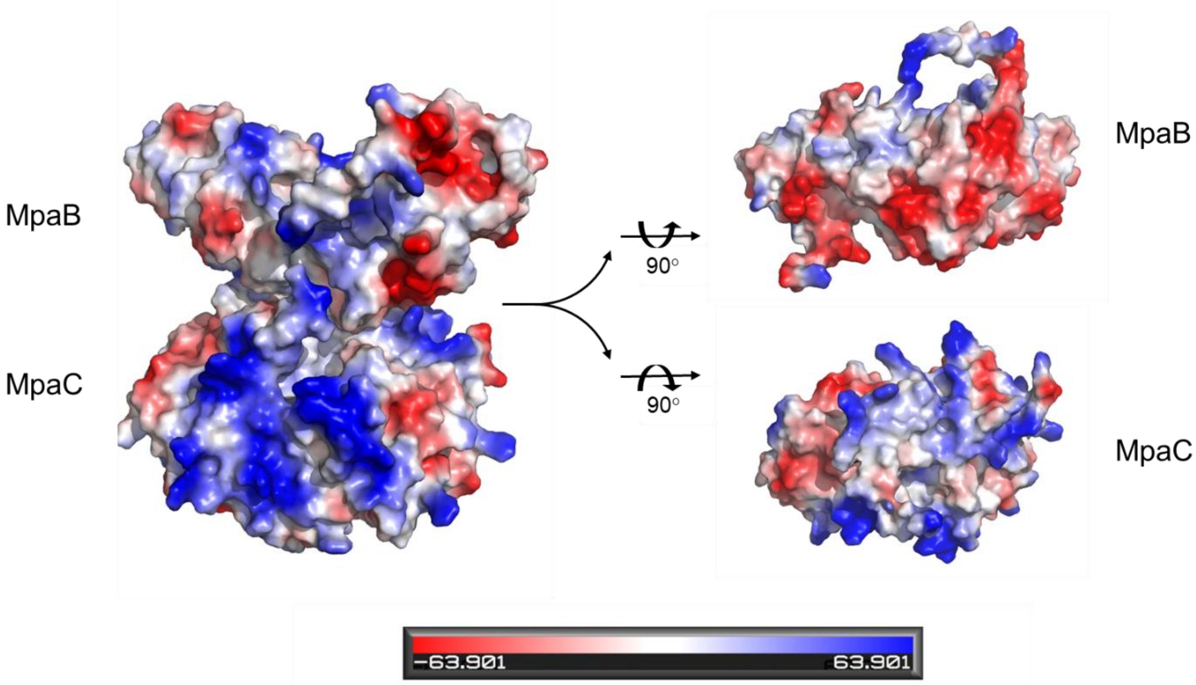

**Supplementary Figure 34: Electrostatic surface potential of MpaB and MpaC.** MpaB (pI = 4.8) and MpaC (pI = 11.2) are shown as a predicted protein complex on the left with positively and negatively charged surface shown in blue and red, respectively. Views of the interaction surfaces are shown on the right by a 90-degree rotation for each protein. The interaction surface is overall negatively charged for MpaB and positive for MpaC. The electrostatic surface potential is calculated by APBS in PyMOL[13, 14].

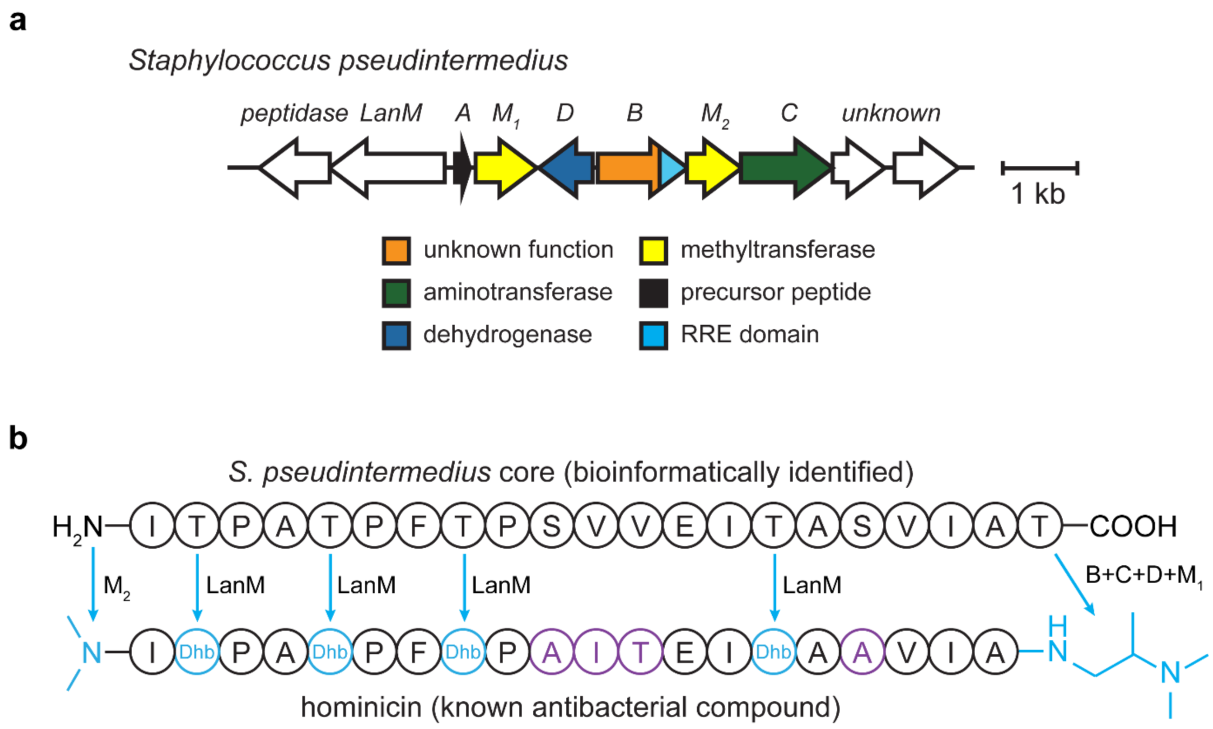

**Supplementary Figure 35: Identification of a hominicin-like BGC from *Staphylococcus pseudintermedius*.** (**a**) The BGC from *S. pseudintermedius* (DapC protein: WP_214539449.1) is shown. Additionally encoded are a LanM which is proposed to convert the four internal Thr residues to dehydrobutyrine (Dhb), a second methyltransferase (putatively dimethylating the *N*-terminus), a peptidase, and two genes of unknown function. (**b**) Comparison of the core peptide region from the *S. pseudintermedius* BGC and the structure of hominicin[5]. Sites modified in hominicin are cyan and the predicted modifying enzymes are listed. Amino acid variations from the *S. pseudintermedius* core are purple.

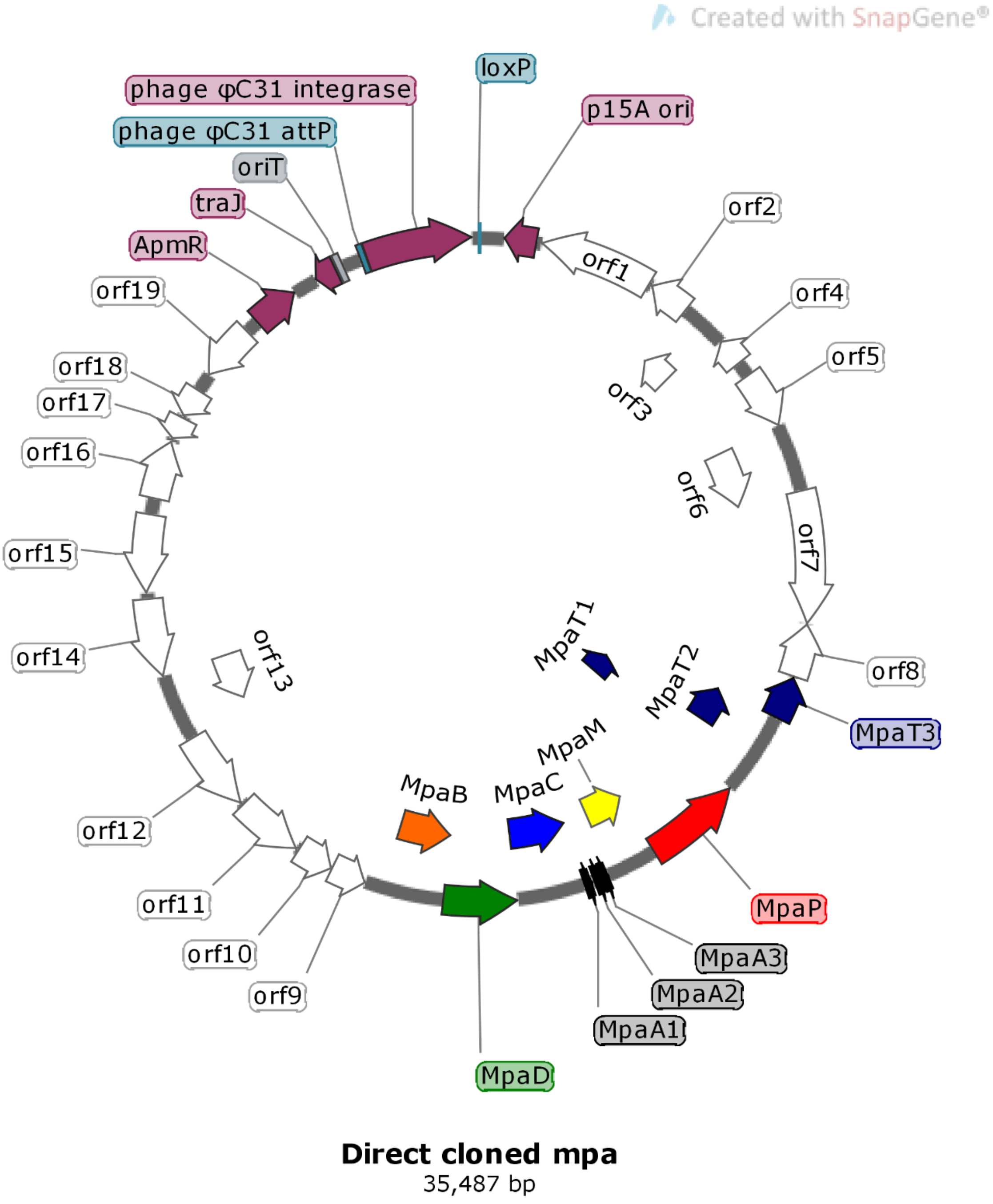

**Supplementary Figure 36: The plasmid map of directly cloned *mpa* for heterologous expression in *S. albus* J1074.** ApmR: apramycin resistance marker; traJ: conjugal transfer transcriptional regulator traJ; oriT: origin of transfer; loxP: Cre-Lox recombination site; p15A ori: p15A plasmid origin of replication. The plasmid map was drawn with SnapGene (GSL Biotech LLC). The plasmid sequence is available in **Source Data**.

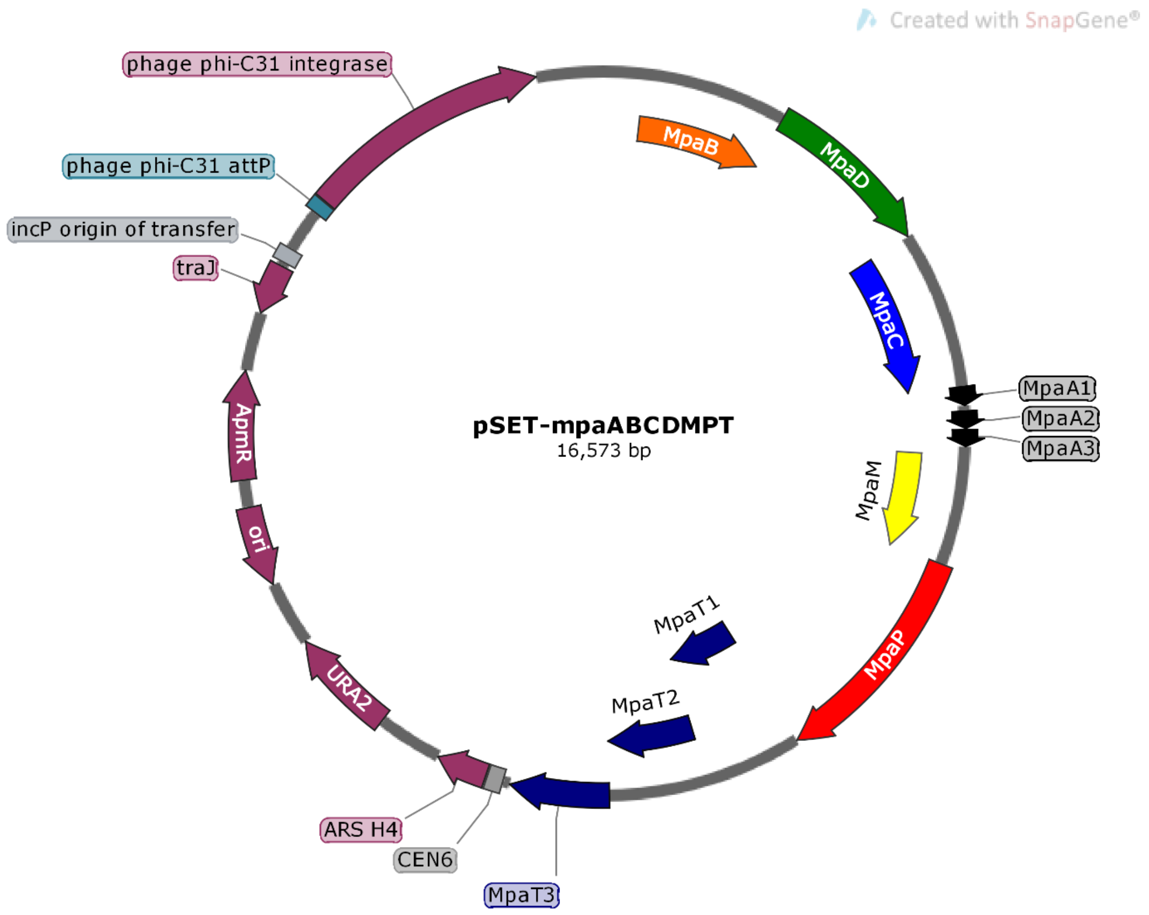

**Supplementary Figure 37: Plasmid maps of gene omission for heterologous expression in *S. albus* J1074: pSET-mpaABCDMPT.** ApmR: apramycin resistance marker; traJ: conjugal transfer transcriptional regulator traJ; oriT: origin of transfer; loxP: Cre-Lox recombination site; ori: p15A plasmid origin of replication; URA2: uracil auxotrophic marker; ARS H4: yeast artificial chromosome; CEN6: centromere in chromosome VI of *Saccharomyces cerevisiae*. The plasmid map was drawn with SnapGene (GSL Biotech LLC). The plasmid sequence is available in **Source Data**.

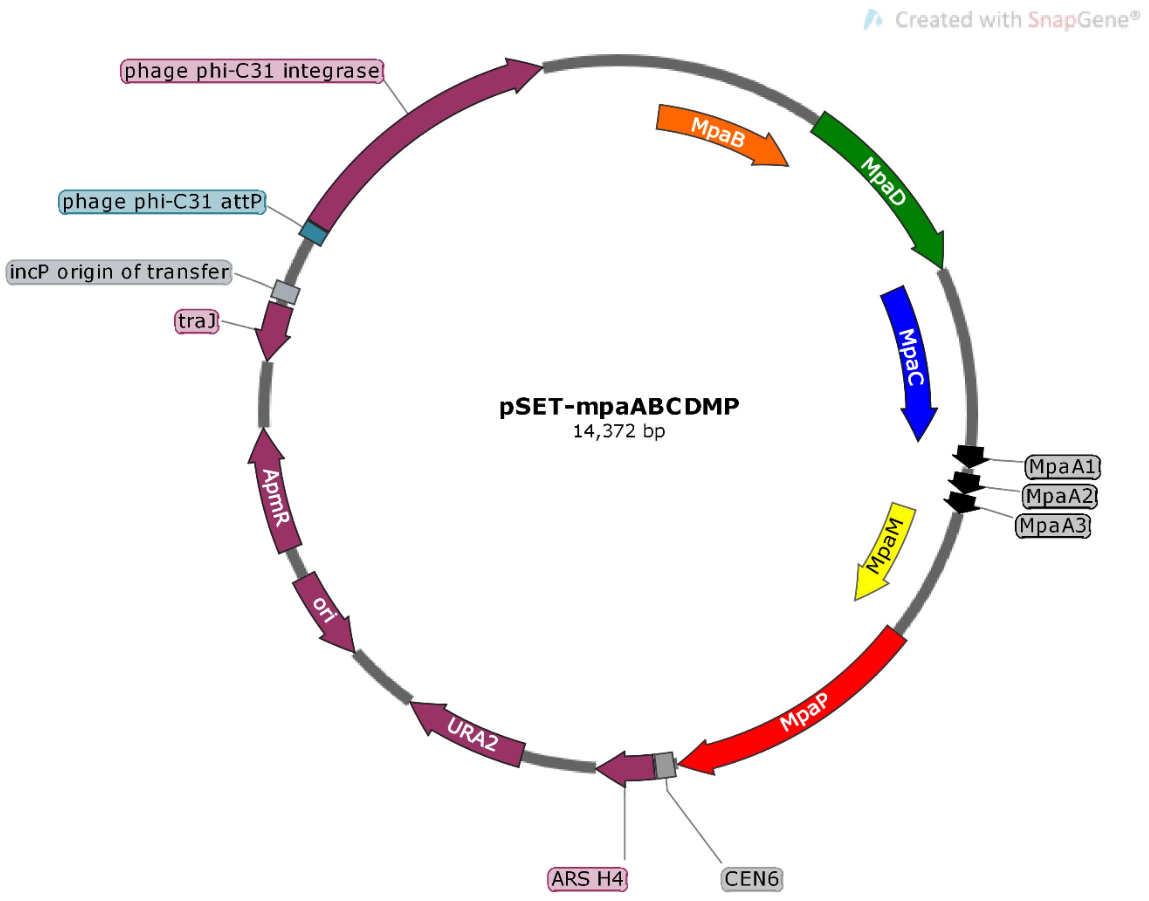

**Supplementary Figure 38: Plasmid maps of gene omission for heterologous expression in *S. albus* J1074: pSET-mpaABCDMP.** ApmR: apramycin resistance marker; traJ: conjugal transfer transcriptional regulator traJ; oriT: origin of transfer; loxP: Cre-Lox recombination site; ori: p15A plasmid origin of replication; URA2: uracil auxotrophic marker; ARS H4: yeast artificial chromosome; CEN6: centromere in chromosome VI of *Saccharomyces cerevisiae*. The plasmid map was drawn with SnapGene (GSL Biotech LLC). The plasmid sequence is available in **Source Data**.

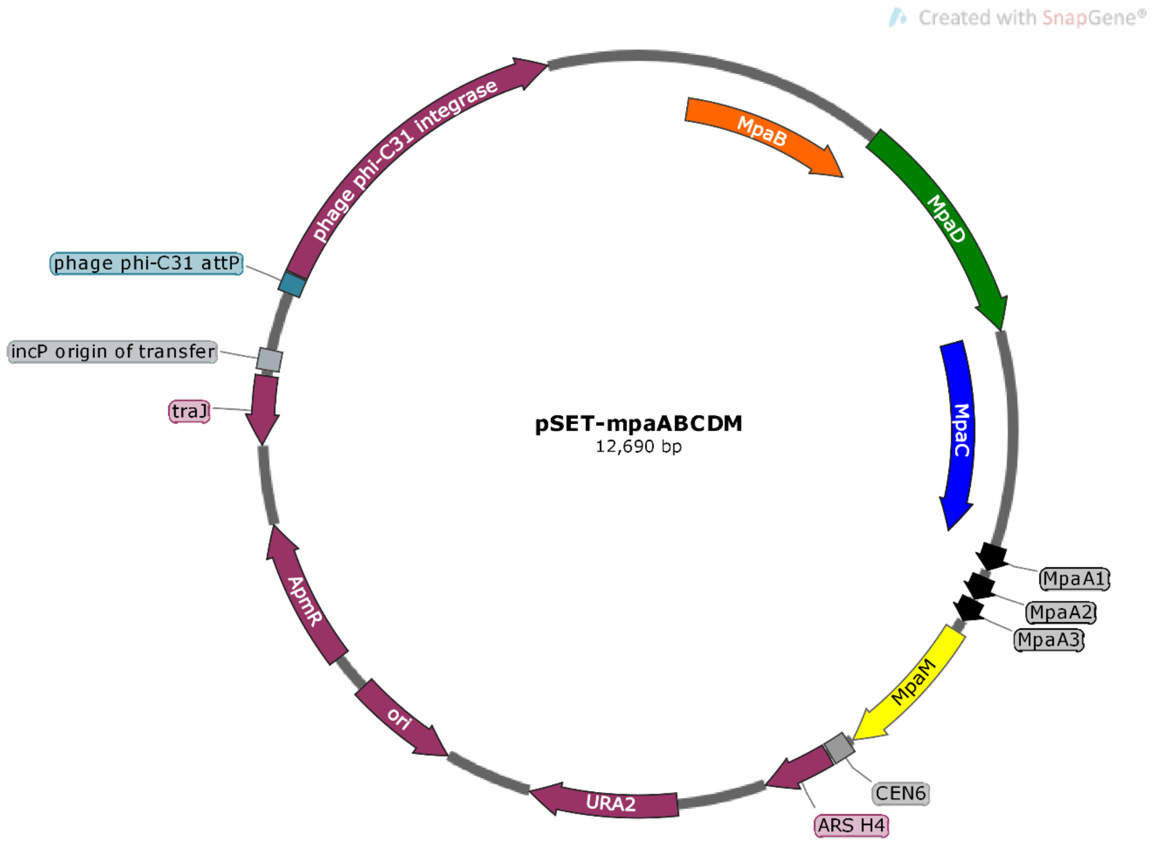

**Supplementary Figure 39: Plasmid maps of gene omission for heterologous expression in *S. albus* J1074: pSET-mpaABCDM.** ApmR: apramycin resistance marker; traJ: conjugal transfer transcriptional regulator traJ; oriT: origin of transfer; loxP: Cre-Lox recombination site; ori: p15A plasmid origin of replication; URA2: uracil auxotrophic marker; ARS H4: yeast artificial chromosome; CEN6: centromere in chromosome VI of *Saccharomyces cerevisiae*. The plasmid map was drawn with SnapGene (GSL Biotech LLC). The plasmid sequence is available in **Source Data**.

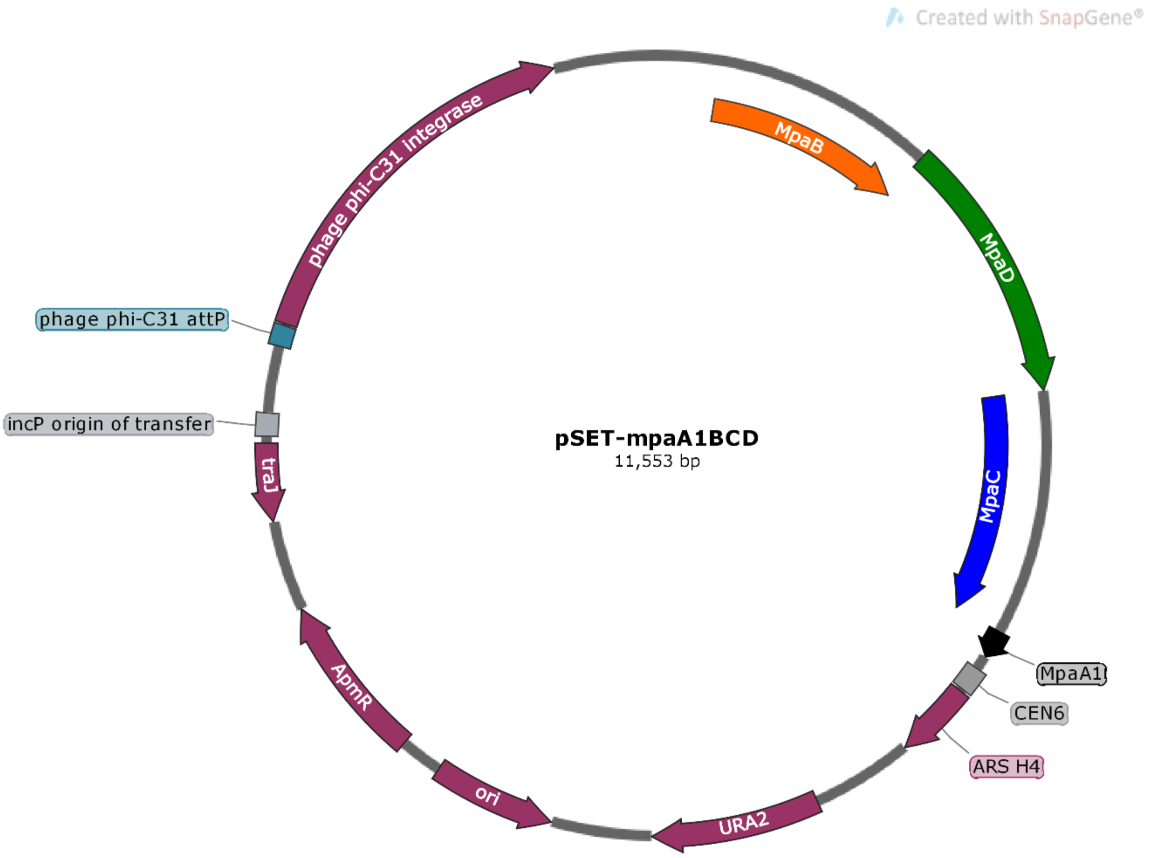

**Supplementary Figure 40: Plasmid maps of gene omission for heterologous expression in *S. albus* J1074: pSET-mpaA1BCD.** ApmR: apramycin resistance marker; traJ: conjugal transfer transcriptional regulator traJ; oriT: origin of transfer; loxP: Cre-Lox recombination site; ori: p15A plasmid origin of replication; URA2: uracil auxotrophic marker; ARS H4: yeast artificial chromosome; CEN6: centromere in chromosome VI of *Saccharomyces cerevisiae*. The plasmid map was drawn with SnapGene (GSL Biotech LLC). The plasmid sequence is available in **Source Data**.

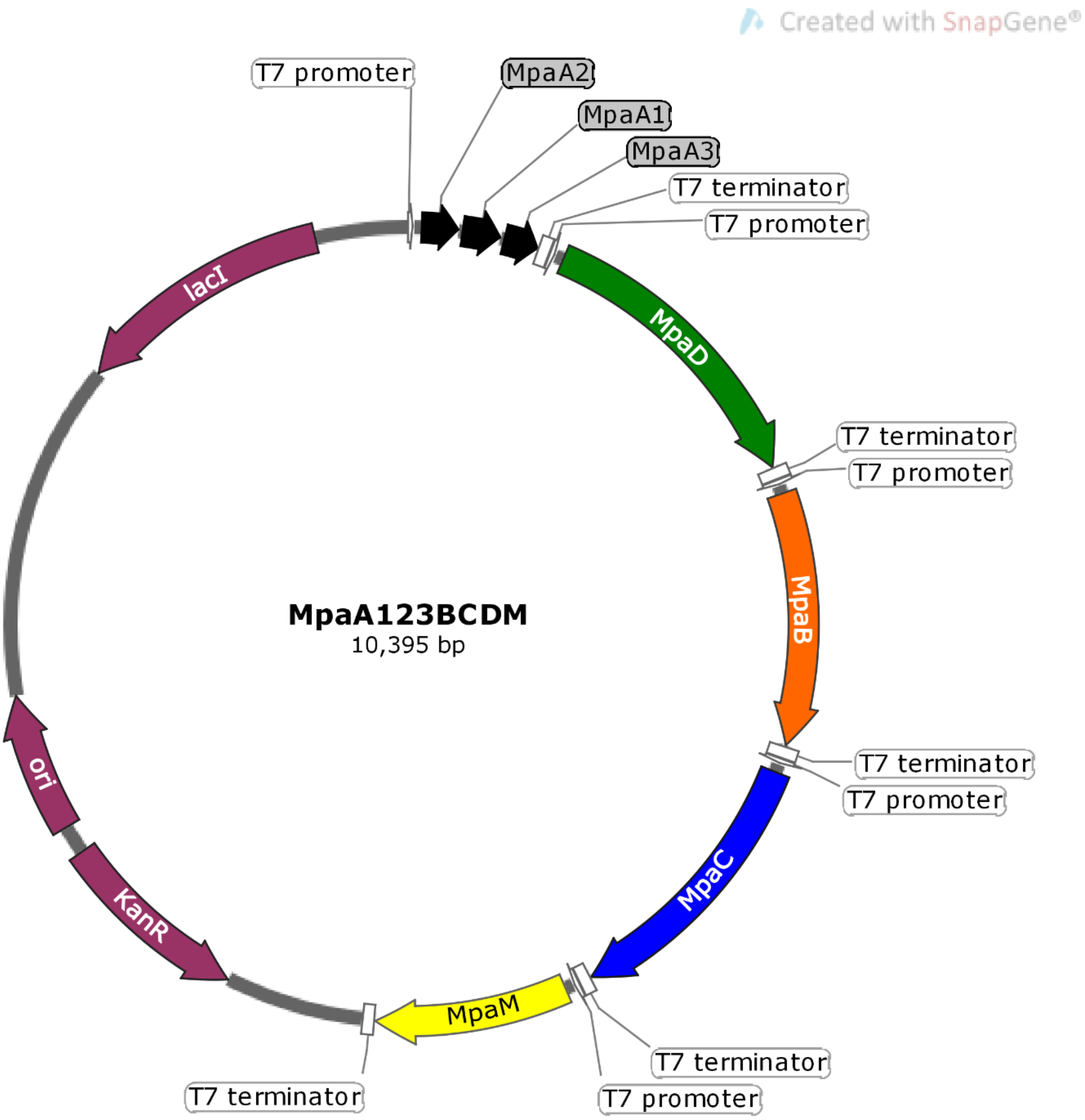

**Supplementary Figure 41: Plasmid maps of codon-optimized and refactored genes for heterologous expression in *E. coli*: MpaA1A2A3BCDM.** KanR: kanamycin resistance marker; ori: pBR322 plasmid origin of replication; lacI: lactose operon repressor lacI. The plasmid map was drawn with SnapGene (GSL Biotech LLC). The plasmid sequence is available in **Source Data**.

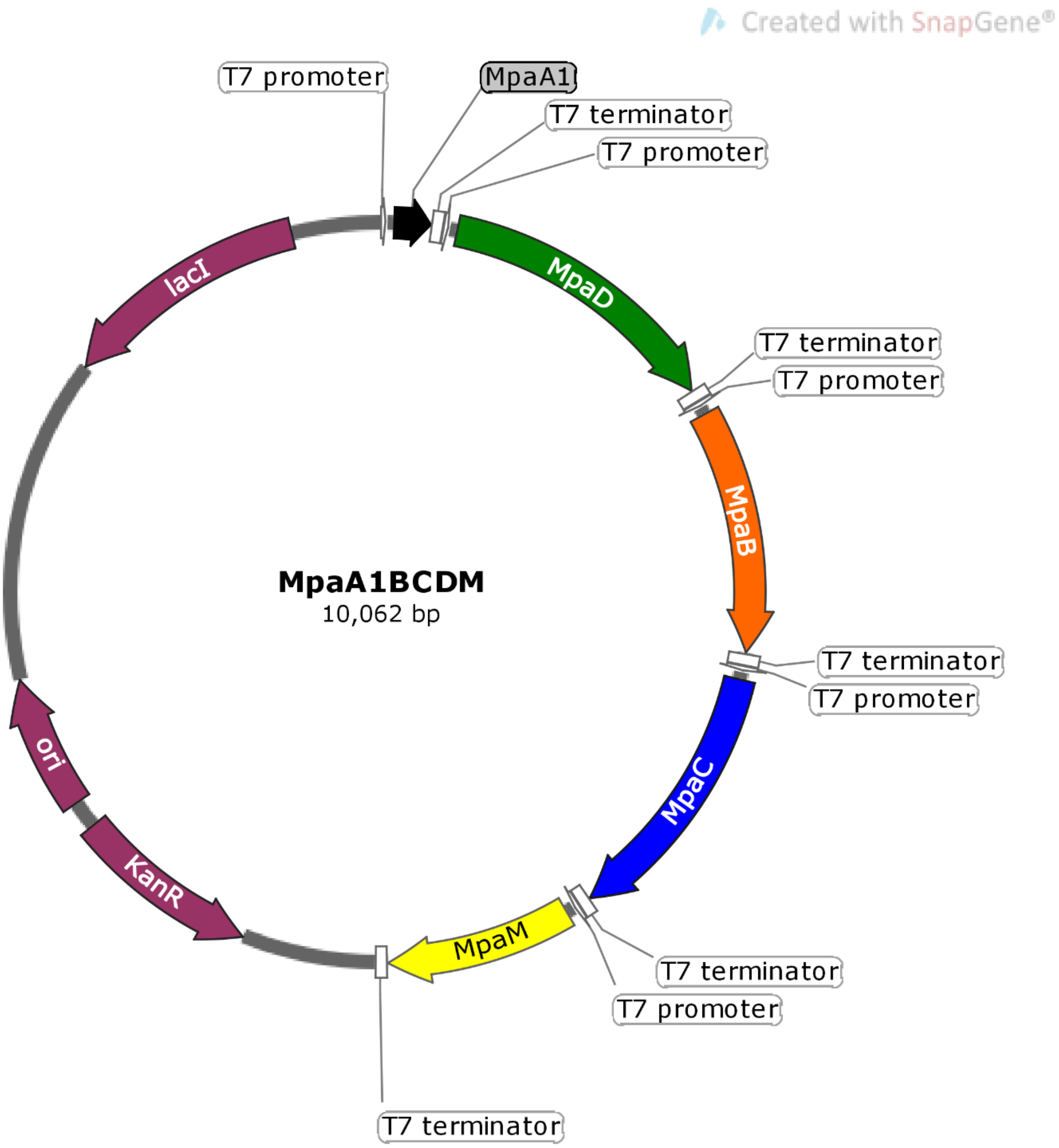

**Supplementary Figure 42: Plasmid maps of codon-optimized and refactored genes for heterologous expression in *E. coli*: MpaABCDM.** KanR: kanamycin resistance marker; ori: pBR322 plasmid origin of replication; lacI: lactose operon repressor lacI. The plasmid map was drawn with SnapGene (GSL Biotech LLC). The plasmid sequence is available in **Source Data**.

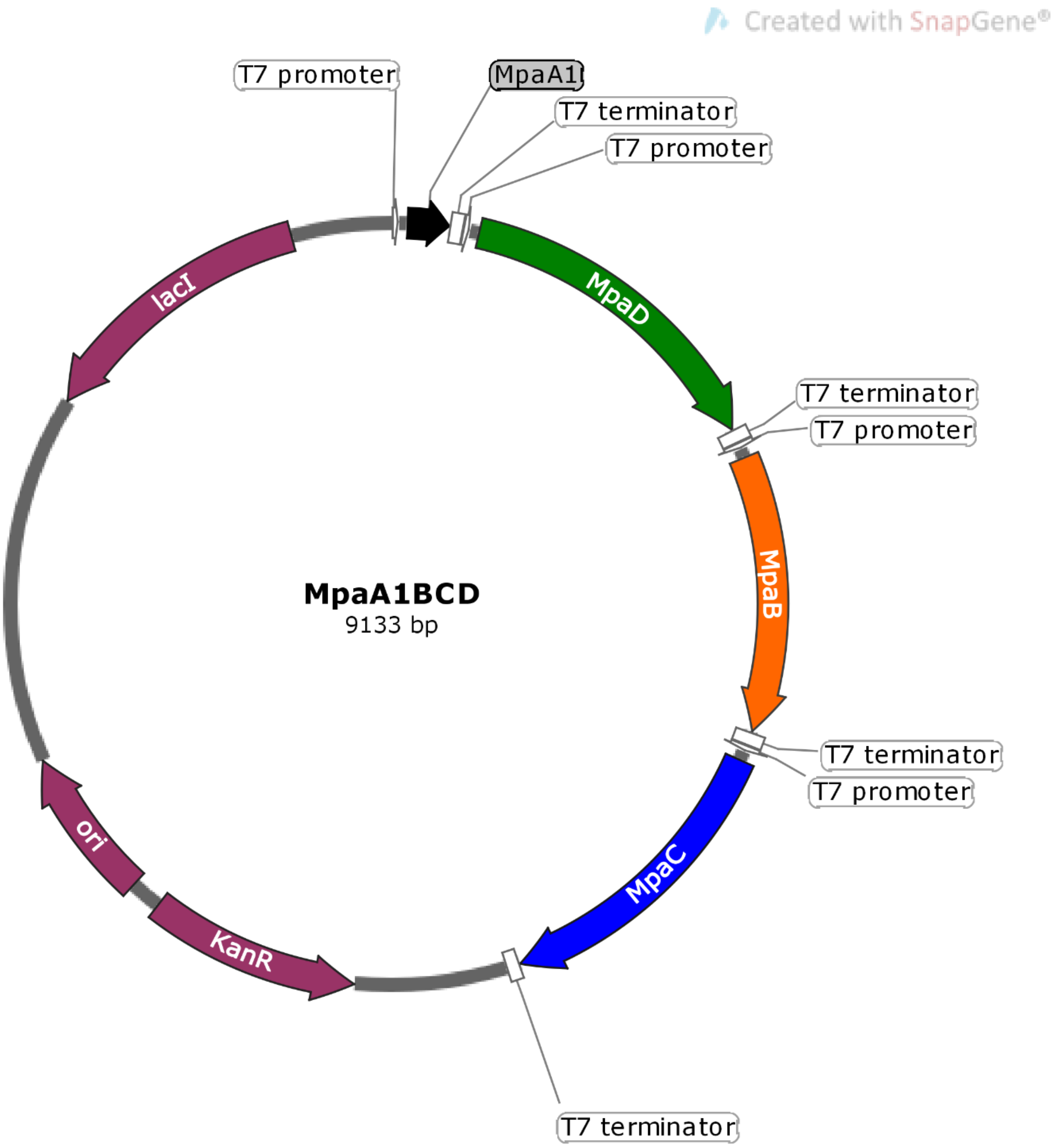

**Supplementary Figure 43: Plasmid maps of codon-optimized and refactored genes for heterologous expression in *E. coli*: MpaABCD.** KanR: kanamycin resistance marker; ori: pBR322 plasmid origin of replication; lacI: lactose operon repressor lacI. The plasmid map was drawn with SnapGene (GSL Biotech LLC). The plasmid sequence is available in **Source Data**.

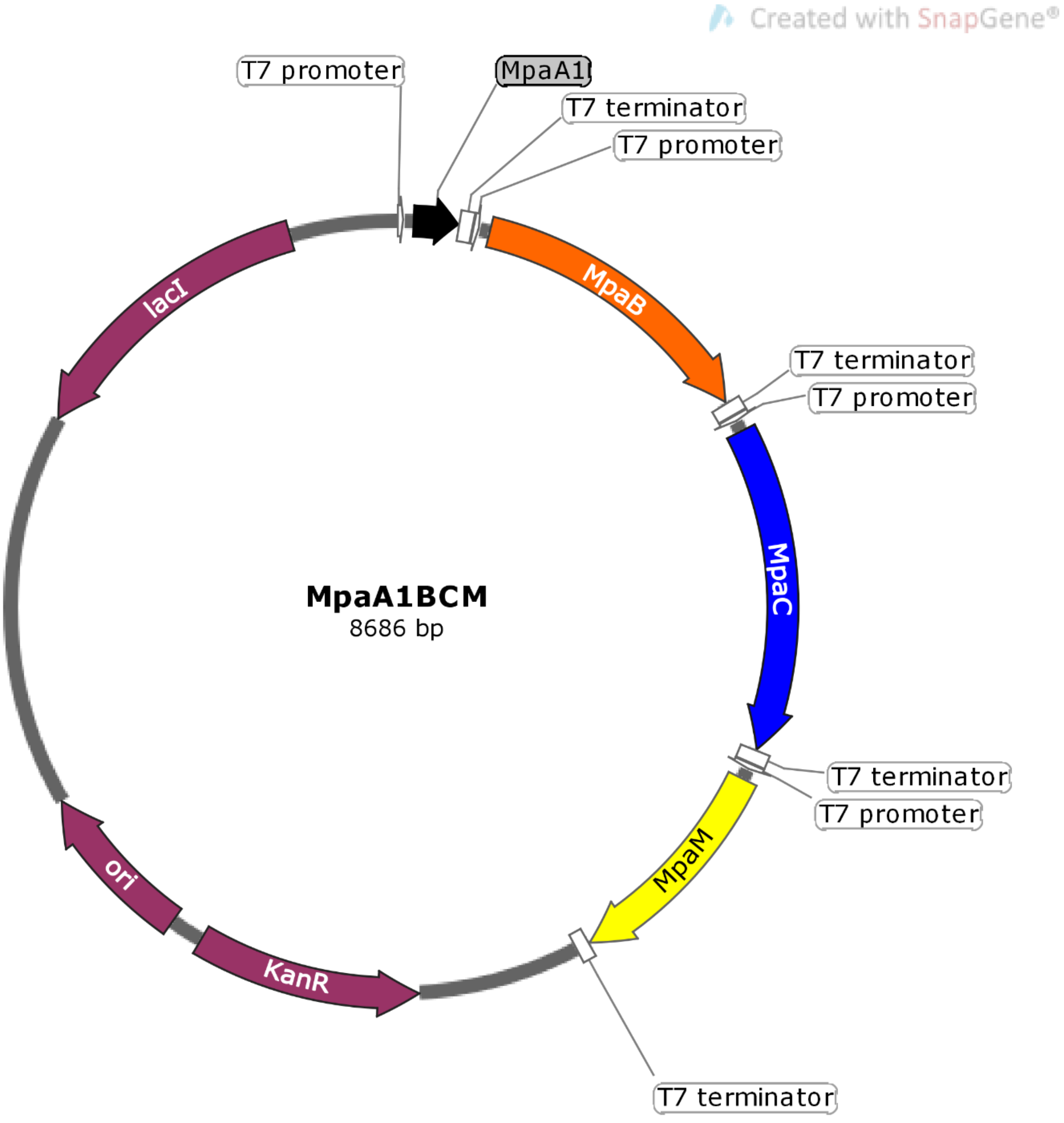

**Supplementary Figure 44: Plasmid maps of codon-optimized and refactored genes for heterologous expression in *E. coli*: MpaABCM.** KanR: kanamycin resistance marker; ori: pBR322 plasmid origin of replication; lacI: lactose operon repressor lacI. The plasmid map was drawn with SnapGene (GSL Biotech LLC). The plasmid sequence is available in **Source Data**.

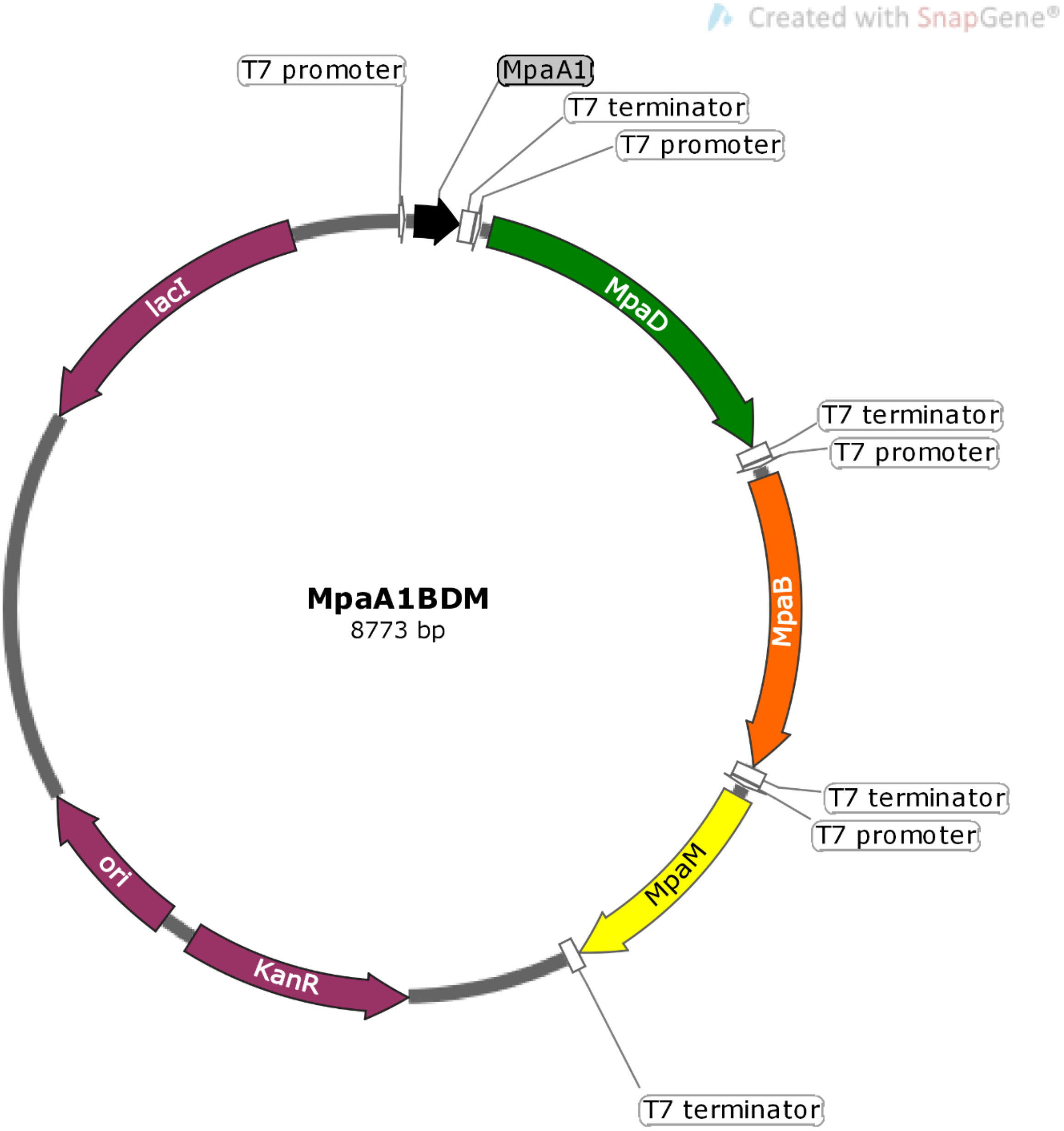

**Supplementary Figure 45: Plasmid maps of codon-optimized and refactored genes for heterologous expression in *E. coli*: MpaABDM**. KanR: kanamycin resistance marker; ori: pBR322 plasmid origin of replication; lacI: lactose operon repressor lacI. The plasmid map was drawn with SnapGene (GSL Biotech LLC). The plasmid sequence is available in **Source Data**.

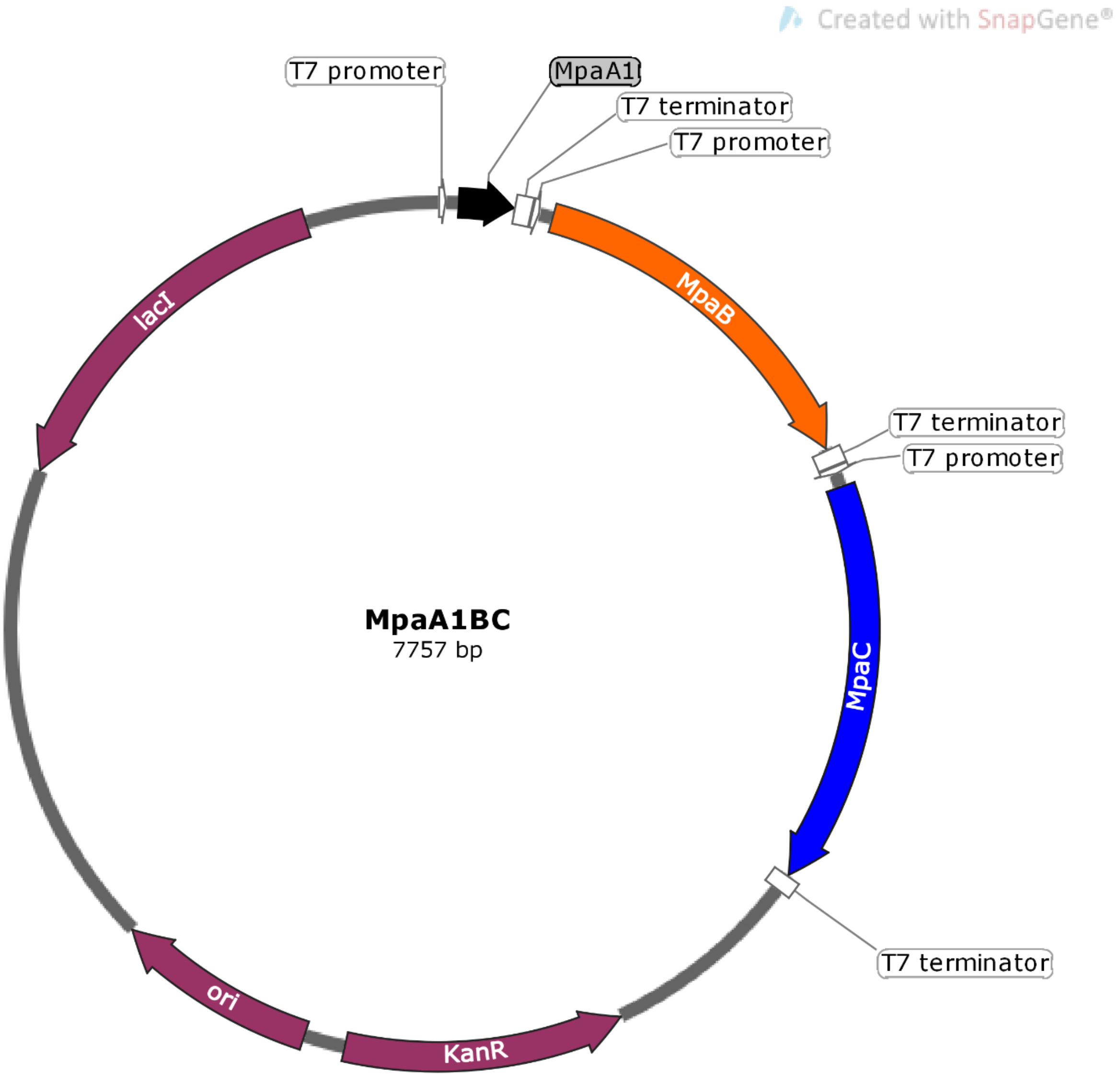

**Supplementary Figure 46: Plasmid maps of codon-optimized and refactored genes for heterologous expression in *E. coli*: MpaABC.** KanR: kanamycin resistance marker; ori: pBR322 plasmid origin of replication; lacI: lactose operon repressor lacI. The plasmid map was drawn with SnapGene (GSL Biotech LLC). The plasmid sequence is available in **Source Data**.

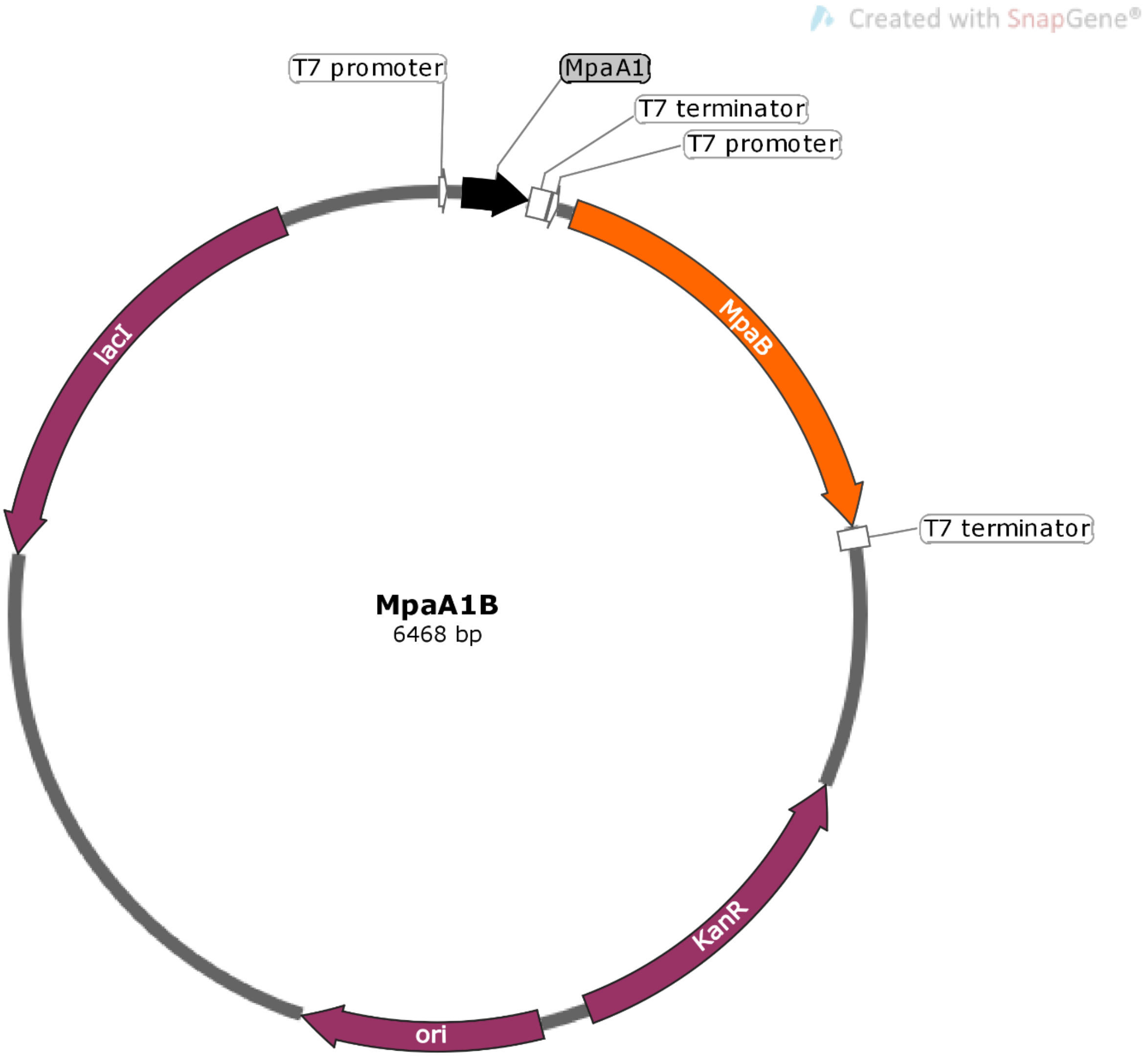

**Supplementary Figure 47: Plasmid maps of codon-optimized and refactored genes for heterologous expression in *E. coli*: MpaAB.** KanR: kanamycin resistance marker; ori: pBR322 plasmid origin of replication; lacI: lactose operon repressor lacI. The plasmid map was drawn with SnapGene (GSL Biotech LLC). The plasmid sequence is available in **Source Data**.

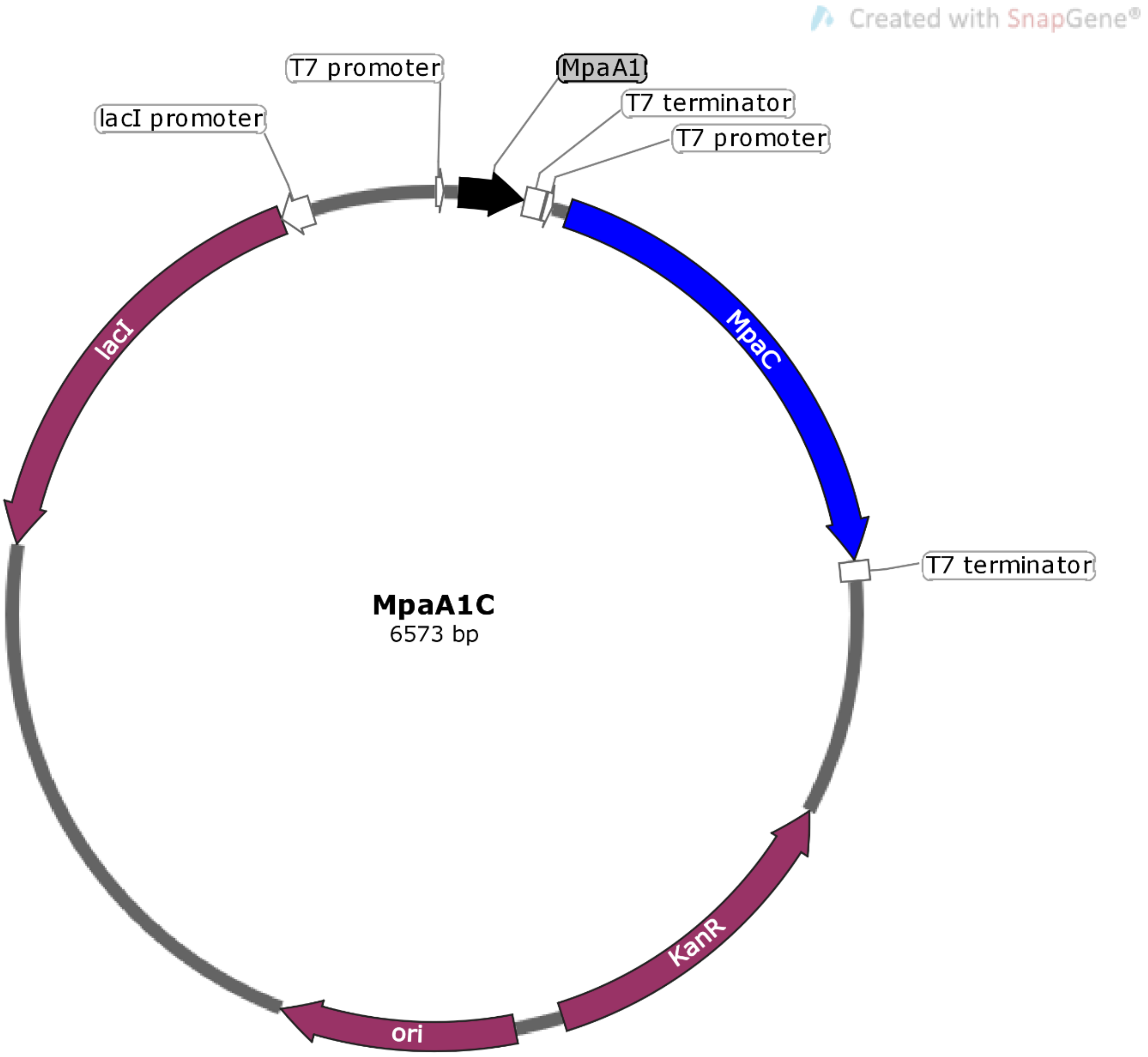

**Supplementary Figure 48: Plasmid maps of codon-optimized and refactored genes for heterologous expression in *E. coli*: MpaAC.** KanR: kanamycin resistance marker; ori: pBR322 plasmid origin of replication; lacI: lactose operon repressor lacI. The plasmid map was drawn with SnapGene (GSL Biotech LLC). The plasmid sequence is available in **Source Data**.
